# Binding of cortical functional modules by synchronous high frequency oscillations

**DOI:** 10.1101/2023.05.20.541597

**Authors:** Jacob C. Garrett, Ilya A. Verzhbinsky, Erik Kaestner, Chad Carlson, Werner K. Doyle, Orrin Devinsky, Thomas Thesen, Eric Halgren

**Author notes:** ^†^These authors made equal contributions to this manuscript.

## Abstract

Whether high-frequency phase-locked oscillations facilitate integration (‘binding’) of information across widespread cortical areas is controversial. Here we show with intracranial EEG that cortico-cortical co-ripples (∼100ms long ∼90Hz oscillations) increase during reading and semantic decisions, at the times and co-locations when and where binding should occur. Fusiform wordform areas co-ripple with virtually all language areas, maximally from 200-400ms post-word-onset. Semantically-specified target words evoke strong co-rippling between wordform, semantic, executive and response areas from 400-800ms, with increased co-rippling between semantic, executive and response areas prior to correct responses. Co-ripples were phase-locked at zero-lag over long distances (>12cm), especially when many areas were co-rippling. General co-activation, indexed by non-oscillatory high gamma, was mainly confined to early latencies in fusiform and earlier visual areas, preceding co-ripples. These findings suggest that widespread synchronous co-ripples may assist the integration of multiple cortical areas for sustained periods during cognition.

**One Sentence Summary:** Widespread visual, wordform, semantic, executive and response areas phase-lock at 90Hz during integrative semantic processing.

## Main Text

A central unanswered question in neuroscience is how information is integrated across the cortex. In psychological studies, this is most often addressed as how different visual qualities of an object (e.g., color, shape, location, texture) are associated with each other^1^. One model, Binding-by-Synchrony (BBS), posits that the constituent features comprising objects are ‘bound’ together when coherent oscillations coordinate activity of the distributed neurons encoding those features^2^. LFP oscillations reflect oscillating membrane potentials in the underlying population, and the consequent modulation of inputs would help select neurons in the currently active network. Furthermore, phase-modulation would tend to synchronize outputs which may be more effective in triggering action-potentials in target structures^3,4^, especially if they arrive at an oscillation peak^5^.

The BBS hypothesis was first supported by phase-locked unit-firing and local field potentials at 40-60Hz evoked by simple visual stimuli in the anesthetized cat visual cortex^6^. Although some further studies found similar results in other cortical areas, behaviors and species^5,7^, others have been less successful, finding that the prominent gamma oscillations evoked by visual stimuli in early visual cortical areas may be modulated by low level properties (e.g., strength, eccentricity) unrelated to binding, and dissociated from synchronized cell-firing^8–10^.

In contrast to BBS which emphasizes the importance of the timing of neuronal firing for transmission and selection of information, other theories propose that a pure rate code is adequate, with oscillations themselves being an inconsequential byproduct of increased firing^11^. Such models generally propose that information is processed via convergent conjunction of labelled lines, arranged in a sequential feedforward hierarchy, with feedback predictive and attentional projections^10,12^. Such models also face various challenges. First, there are insufficient neurons to code for all possible combinations of events and actions, so ultimately they must be encoded with a network. Second, many stimuli have not been experienced before sufficiently to have labelled lines. Third, unlike other mammals, most of human cortex is within a narrow range of the same level of hierarchy, as indicated by relative myelin content levels^13^. Thus, a hierarchical model is not applicable to the majority of human cortex. Fourth, intracranial recordings show that in cognitive tasks, stimulus-evoked activity proceeds through the hierarchically-organized cortex long before a response is produced. For example, visually presented words pass through the wordform area, at the apex of the hierarchically-organized visual cortex by ∼240ms^14–19^, at which time their semantic categories evoke distinct responses^20^, but semantically-defined behavioral responses are ∼600ms later and involve much more of the brain^14,21,22^. Fifth, stimulus-evoked activation of labelled lines does not explain neural activity underlying spontaneous behavior, which could reasonably be construed to constitute most behavior. Interestingly, such ‘uninstructed movements’ were the main determinant of cortical activity in widefield 2-photon imaging of the dorsal cortical surface^23^.

These considerations suggest that convergent hierarchical conjunction of labelled lines maybe be most effective for initial processing of familiar stimuli, but later integrative processing might be facilitated by BBS. For example, the finding that unexpected stimuli in monkey visual cortex evoke increased firing-rates whereas mutually predicted stimuli evoke increased firing-synchrony^24^ led to models wherein activity in feedforward convergent labelled lines are bound by feedback BBS into coherent patterns^25,26^. This formulation of BBS emphasizes synchronous cell-firing, and accordingly applies to only some gamma oscillations, especially short bursts that modulate cell firing and located in higher levels that support feedback processing^26^.

BBS has mainly been examined in early visual areas of lower mammals, but synchronous high gamma oscillations also occur in human cortex where they may take the form of ∼90Hz oscillations lasting∼100ms. These ‘ripples’ occur throughout the cortex, during spontaneous waking and non-rapid eye movement (NREM) sleep^27^. They are commonly phase-locked between sites in different hemispheres and lobes at long distances, with zero phase-lag in waking^28^, and phase-modulate local cortical neuronal firing^28–30^. Expressed in neurophysiological terms, the central prediction of BBS is that the firing of neurons are more integrated if their locations are oscillating synchronously, and indeed, in human superior temporal and precentral gyri neurons in different cortical locations predict each other’s firing patterns more strongly when those locations are co-rippling^30^. This co-prediction is independent of increased activation, but strongly modulated by ripple phase, as hypothesized by BBS^5^. In the one previous report of cortical co-rippling in humans during a cognitive task, co-ripples increased prior to correct recall in a paired associates task^28^. However, only five patients were studied with SEEG, and the electrode sampling was insufficient to reveal the spatio-temporal evolution of the co-rippling. Nonetheless, the efficacy of co-ripples in co-predicting neuronal firing in different locations, and their ubiquity across cortical regions and states suggest that they may contribute to binding beyond perceptual features, possibly consistent with the view of binding in philosophy where it refers to the general problem of how the various contents of a mental event are united into a single experience^31^.

In the current study we report the first detailed examination of cortico-cortical co-rippling during a cognitive task. Local field potentials were recorded from widespread cortical areas while subjects read words and made a semantic judgement. Words evoked strong co-rippling between fusiform wordform and widespread language related areas from ∼200 to 400ms, then from ∼400 to 800ms target words evoked stronger co-rippling between visual, wordform, semantic, executive and response-related areas. Transient increases in high-frequency non-oscillatory activity also co-occurred with about the same density as ripples, but they showed very little task modulation, with the exception of the shortest latencies in early visual areas and the fusiform wordform area, where they preceded ripples. Except for early visual areas, co-rippling sites were phase-locked at zero-lag. These results suggest a mechanism whereby following early perceptual processing, cognitive integration may be supported by sustained widespread bursts of synchronous high-frequency oscillations.

## Results

### A reading task evokes widespread cortical co-ripples

Subjects key-pressed to words referring to animals, while consonant strings or false fonts matched for length served as sensory controls (Fig. 1A). ECoG electrodes, placed semi-chronically on the cortical surface for diagnosis of epilepsy, were localized to language-related ROIs using superimposed MRI/CT (Fig. 1B). We predicted that ripples would co-occur (‘co-ripples’) between locations and with latencies that correspond to the sequential binding of legal letter-strings into wordforms, wordforms with their meanings, meaning with task instructions, and task instructions with the behavioral response (Fig. 1C). Ripples were detected with previously described procedures (see Methods). Their waveforms (Fig. 1DE) and characteristics (Fig. 1F-I)^27,28,30,32–34^ resembled those recorded with SEEG and Utah array electrodes within the cortex, except for 10 locations in early visual cortex with high amplitude and density. Unlike other ripples at other locations, these occurred at short latency and very high rates, especially to false fonts (Fig. 1JK). Such sites (∼1% of recordings) were excluded from analysis. Ripples in different sites had a strong tendency to co-occur (Fig. 1L), at a rate that decrements very slowly with distance (R^2^=0.021; b=-0.0025; Fig. 1M). Only left hemisphere sites were included in analyses unless specified otherwise.

**Fig. 1.**
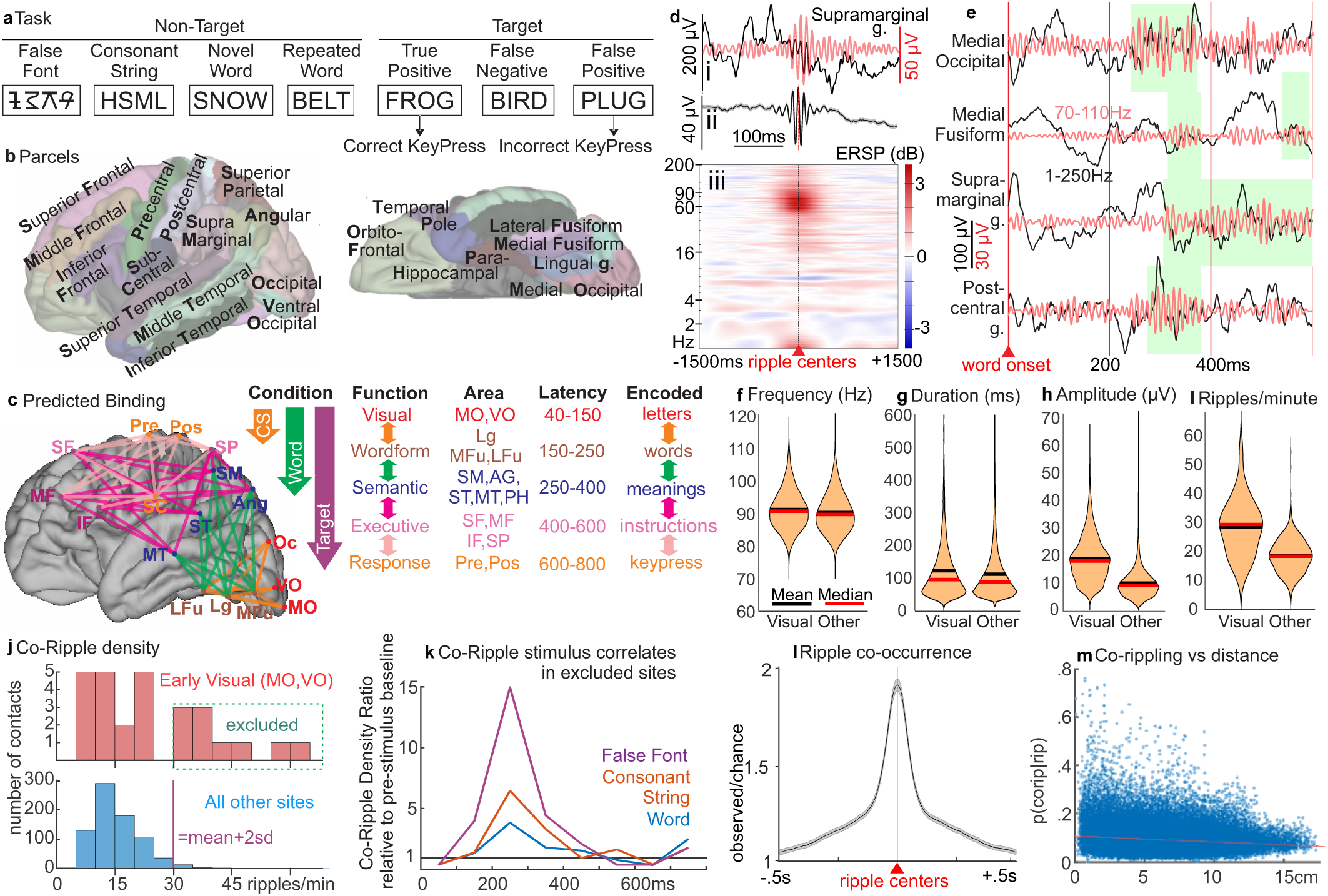
Cortical Ripple and Co-Ripple Characteristics During a Reading Task. **A**. Subjects read words and control stimuli (including consonant strings and false fonts) and key-pressed to names of animals. Stimuli were presented continuously at 600ms or 1000-1400ms intervals. **B**. Regions of Interest chosen for amalgamating responses across subjects (for details see supplementary table S2). Capital letters indicate the abbreviations used for these ROIs in this and subsequent figures. **C**. Predicted successive stages of binding from visual input to keypress response. **D**. Example high frequency oscillation (‘ripple’) from typical disc electrode on the cortical surface as single sweep (**Di**; broadband (black) or band-passed (red)), average waveform (**Dii**), or average time-frequency plot (**Diii**). **E**. Example co-ripple, single sweep to a novel word, multiple sites with overlapping ripples (green shading), broadband (black trace) and band-passed (red trace). **F-I.** In most locations (‘Other’), mean ripple frequency (**F**: 90.3Hz), amplitude (**G**: 10.6µV), duration (**H**: 113.5ms), and rate (**I**: 18.6/minute) are all similar to previously reported during spontaneous activity^27^. However, ripples in early visual cortex regions VO and MO (‘Visual’) had ∼85% higher amplitude (**H**) and ∼50% higher occurrence rate (**I**). **J**. Co-ripple density in VO and MO channels were either within the same the same distribution as the other channels, or in 10 channels were more than 2 standard deviations above the overall mean. **K**. Co-ripples in these channels increased strongly at short latency to all visual stimuli, especially false fonts, unlike other sites. ripples in these 10 outlier contacts (1.3% of total) were excluded from further analysis. **L**. Cortical ripples often co-occur within ∼1s of each other (shaded area denotes SEM), and (**M**) overlap in time (‘co-ripples’) with a probability that is mainly independent of distance (Euclidean distance; R^2^=0.021; b=-0.0025/cm [−0.0026, −0.0024]; p=10^−310^, t=-37.9, df=68634, two-tailed t-test).

### Words evoke co-ripples between wordform and widespread areas

related to language from ∼100-400ms after word onset. We grouped contacts in anatomo-functional regions (Fig. 1B) and tested for co-ripple density in 100ms epochs from 0-400ms after word onset, compared to the pre-stimulus baseline (Fig.2). Increased co-rippling is present from 100-400ms, between the fusiform wordform area and areas related to semantic (supramarginal gyrus, angular g., inferior, middle, and superior temporal gyri, and temporal pole), executive (inferior, middle and superior frontal g., and superior parietal lobule), and response (pre- and post-central g.) processing needed for task performance (Fig. 2C, 3AEF). Remarkably, Co-R between non-fusiform language-related sites does not increase. The early co-rippling to words between fusiform wordform and other language-processing areas largely ends after ∼400ms for non-target stimuli (Fig. 3AEF). Although consonant-strings also evoke fusiform Co-R, these are significantly smaller than those to words (Fig. 3B) and decline sharply from 200ms while those to words continue to increase (Fig. 3E, dotted cyan line).

**Fig. 2.**
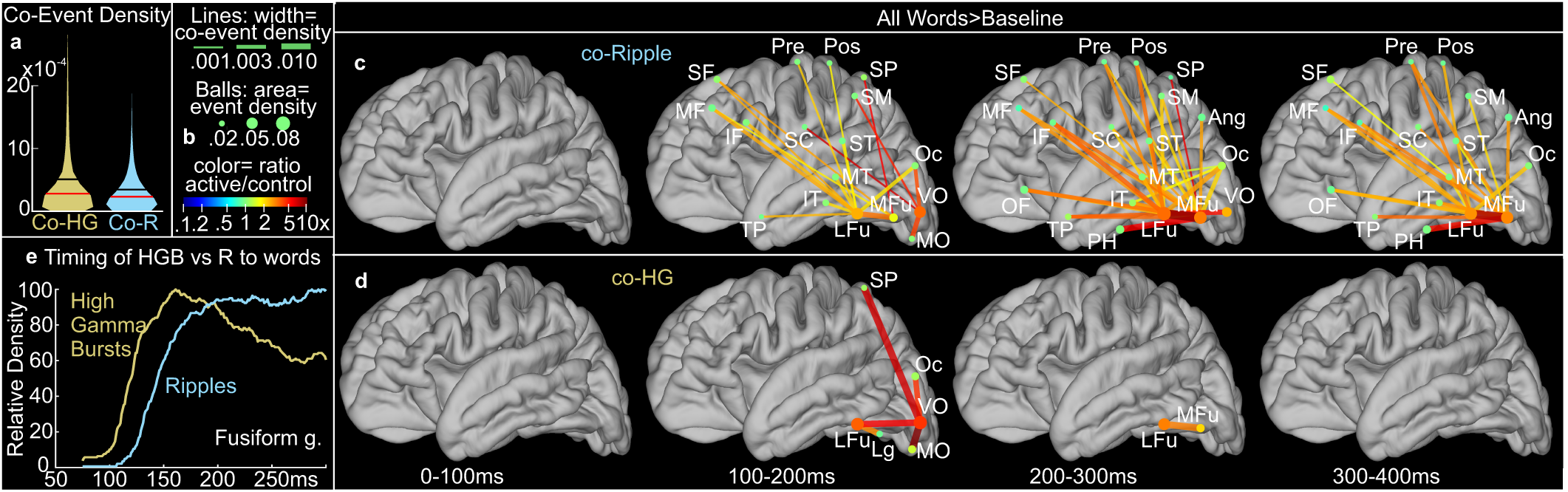
Co-Ripple and Co-High Gamma Burst Initial Responses to Words. **A**. Co-occurrence density of ripples (Co-R) and of non-oscillatory high gamma bursts (Co-HG). Co-event density is defined as the proportion of total task time that Co-R (or Co-HG) in 2 channels are both active. The means (3.69 for Co-R, 5.48 for co-HG), medians (2.48; 3.05), and distributions are similar to each other. **B**. Connection plots were constructed for task contrasts where line color indicates the ratio of co-event (either Co-R or Co-HG) density between the connected areas evoked by active versus control conditions, and line thickness is proportional to co-event density in the active condition. All lines are p<0.01, 1-tailed, permutation test, FDR-corrected. Dot color indicates ratio of event density at that site by active versus control conditions, and dot area is proportional to event density in the active condition, organized by anatomical regions (Fig. 1B, Table SX). Same legend for panels 2CD, 3A-D, 4AC, and 5A-H. **C**. Co-R significantly increase over baseline for a sustained period beginning 100-200ms following word onset with the wordform area (Lateral and medial Fusiform; LFu, MFu) projecting to virtually all left hemisphere language-related areas (areas and abbreviations mapped in Figs. 1BC). **D**. Co-HG increases strongly but more briefly and sparsely than co-ripples between early visual sites (MO, VO, Oc, Lg), with some connection to LFu at 100-200ms, and between LFu-MFu at 200-300ms. Compared to Co-R with 18, 27 and 21 significant connections in the 100-200, 200-300 and 300-400 epochs, Co-HG task-modulated co-events are more sparse and earlier (5, 1 and 0 significant connections, respectively). **E**. High gamma bursts (HGB) lead R when they both occur on the same Fusiform g contact between 75 and 300ms post stimulus onset. Of the 320 such trials, HGB preceded R in 68.1% (62.7-73.1%, p=10^−10^, two-tailed binomial test; mean latency difference 24.5ms). This test was repeated in the one channel that had sufficient trials (n=200), with HGB preceding R in 78.5% of trials (72.1-84.0%, p=10^−15^, two-tailed binomial test, mean latency difference 35.7ms).

**Fig. 3.**
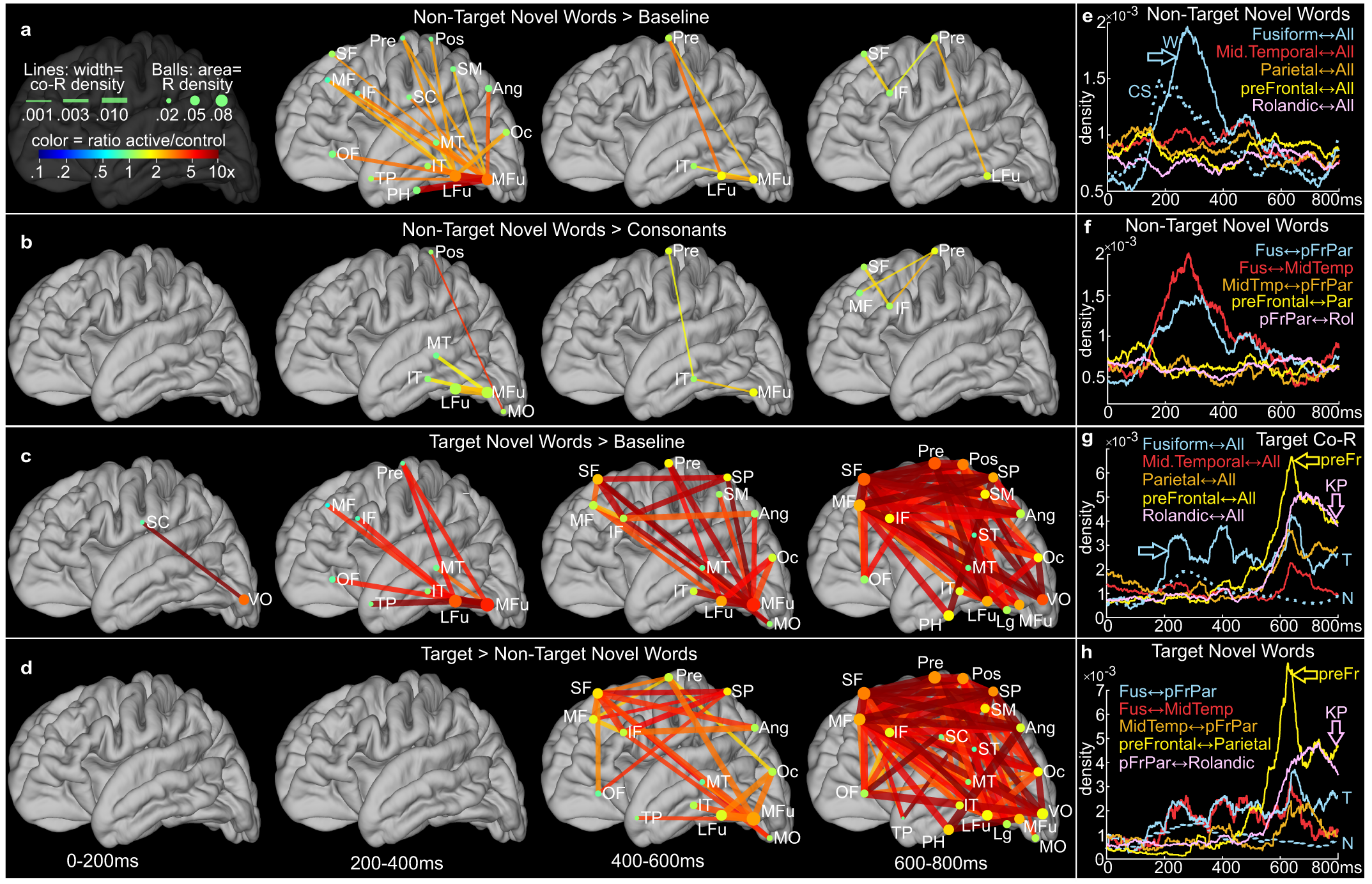
Cortical Co-Ripples Selective for Words and Semantic Decisions. **A-D**. Increased co-ripple density between left hemisphere sites from 0 to 800ms post-stimulus in 200ms increments, between (**A**) Non-Target Words versus Baseline (**B**) or Consonant-Strings, and (**C**) Target Words versus Baseline or (**D**) Non-Target Words. Compared to baseline, reading Non-Target (A) or Target (C) Words evokes from 200 to 400ms increased co-rippling between wordform (MFu, LFu) and other language-related sites, including semantic (Ang, SM, MT, IT), executive (SF, MF, IF), and response (Pre, Pos) areas, generally larger to words than consonant strings. Compared to Non-Targets, Targets evoke from 400-800ms strong co-rippling linking semantic, executive and response (Pre, Pos) areas (D). See Fig. 1BC for parcels. Line color indicates Co-R active/control ratio; all lines are p<0.01, 1-tailed, permutation test, FDR-corrected (see Fig. 2B). **E-H**. High temporal resolution co-ripple density plots from selected groups of regions to all other groups (E, G), or between selected groups (F, H). **E,F**. Non-Target words evoke increased Fusiform Co-R with many other areas (cyan arrow), beginning at ∼130ms post-stimulus onset, peaking at ∼275ms, largely resolving by∼400ms but continuing past 600ms. This broad increase is larger to non-target words than consonant-strings (cyan arrow, 3E; two-tailed t-test, 200-400ms Fus-all Co-R density, 0.7×10^−3^ [0.3, 1.2×10^−3^], t=3.39, df=5591, p=0.001). With few exceptions, only connections including the fusiform increase co-rippling to Non-Target words, possibly seeding wordform information in other language areas. **G,H**. Target words evoke strong sustained Co-R with Fusiform (cyan arrow, 3G), much greater than Non-Targets (dotted cyan line, 3G; two-tailed t-test, 200-400ms Fus-all Co-R density, 2.2×10^−3^ [0.8, 3.6×10^−3^], t=3.11, df=3162, p=0.002) and unlike Non-Targets further evoke strong Co-R between virtually all language areas, led by prefrontal at ∼300ms exponentially rising to peak at ∼640ms. Virtually all other language-related zones increase Co-R with each other beginning ∼80-200ms after the prefrontal cortex, but peak together, except Rolandic cortex which peaks shortly before the Key-Press (pink arrows).

### Co-high gamma burst increases are early, brief, and more restricted

following words, relative to co-ripple increases. In order to test the assertion that ripples do not actively facilitate transcortical integration but are simply a byproduct of nonspecific activation^10,11^, we compared co-rippling to the co-occurrence of bursts of non-oscillatory very high gamma (‘co-HG’; 120-190 Hz), a measure of nonspecific activation^35^. The mean and median density of co-occurrence of high gamma bursts was slightly higher than that of co-ripples, and their distributions were similar (Fig. 2A). Co-HG increased strongly but sparsely and briefly between early visual sites (MO, VO, Oc, Lg), with some connection to lateral fusiform at 100-200ms, and between lateral and medial fusiform at 200-300ms. Because LFu and MFu were involved at early latencies for both Co-R and Co-HG, we tested for sequential activation by identifying trials where HGB and R were detected on the same Fusiform g contact between 75 and 300ms post stimulus onset, finding that HGB preceded R in ∼70% of trials by an average of ∼25ms (Fig. 2E).

### Target words strongly and selectively evoke co-ripples

between visual, wordform, semantic, executive and response areas. Co-rippling strongly increased on trials where the subject key-pressed correctly to animate words, compared to pre-stimulus baseline or to non-target novel words (Figs. 3CDGH, S3). Increases in co-rippling density between left hemisphere areas could be greater than twenty-fold (Tables 1, S6). As described above, the initial Co-R increase is between fusiform and regions in all lobes starting at ∼130ms post-stimulus onset. This broad increase is larger to non-target words than consonant-strings (, peaks at ∼275ms, and largely resolves by∼400ms. Target words evoke strong co-R from Fusiform (Fig. 3CG cyan arrow) which is much greater than to non-targets from ∼180ms (dotted cyan line in Fig. 3G; note vertical scale). Target-evoked Co-R with Fusiform continues to >800ms, with peaks at ∼250, 400 and 640ms, but nothing notable occurs in other areas until ∼3-400ms when pre-frontal co-R with other areas (especially parietal) increase exponentially with a doubling time of 62ms to a sharp peak at ∼640ms. (exponential fit R^2^=0.98; Fig. S2). Similar but weaker sharp rises in co-R between other language-related zones follow those with prefrontal by ∼80-200ms. Co-R between these sites also sharply peak at ∼640ms (i.e., with prefrontal), except for Rolandic cortex whose broad peak at ∼730ms (pink arrows) precedes the average behavioral response at 789ms.Thus, Co-R to Target Words are centered successively on fusiform, prefrontal-parietal, widespread-language, and Rolandic cortices. Similar results were found with 100ms epochs (Figs. 2, S4).Peak co-rippling occurred in the response period (600-800ms). Correct performance in this task requires that the meaning of the word be evaluated against the criterion ‘animal,’ and if it matches to respond with a keypress. Maintenance of the response criterion and consequent action in active memory, comparison of stimulus word meaning to the response criterion, and procedural sequence control, are all critical but poorly understood ‘executive’ processes that are supported by prefrontal and parietal cortices^36,37^. In the critical response epoch (600-800ms), prefrontal and superior parietal cortices are strongly coupled with areas thought crucial for encoding word meaning, which could support comparison of stimulus word meaning to the response criterion. Prefrontal coupling is also very strong with motor areas, which could support linking the target detection to the required response. Thus, target-selective co-ripples peak in appropriate anatomical locations and post-stimulus latencies for binding stimulus word meaning to target meaning, and for binding stimulus-target matches to appropriate motor response, as predicted by our hypothesis.

### Correct responses evoke higher co-ripple density

especially between semantic, executive and response areas. Although we could not selectively disrupt co-rippling in humans to test if it is necessary for binding, we were able to examine if incorrect responses were associated with decreased co-rippling. Compared to missed responses to targets, correct responses were associated with strongly increased co-rippling between wordform, semantic (supramarginal and angular), executive (prefrontal), and response (pre- and post-central) areas (Figs. 4A; S6). A similar but weaker pattern of increased co-rippling occurred in the 200ms immediately preceding True Positive versus False Positive keypresses (Fig. 4BC, S6; this comparison was only possible in the 6 subjects with sufficient key-presses to non-target words). Note that on both True and False Positive trials, the subject read a word and made a keypress; thus sensory and motor components of the trial were matched. Thus, correct detection of target words was differentiated from incorrect detections by increased co-rippling between executive, semantic and response areas, the critical interactions where binding is predicted to be necessary for correct performance.

**Fig. 4.**
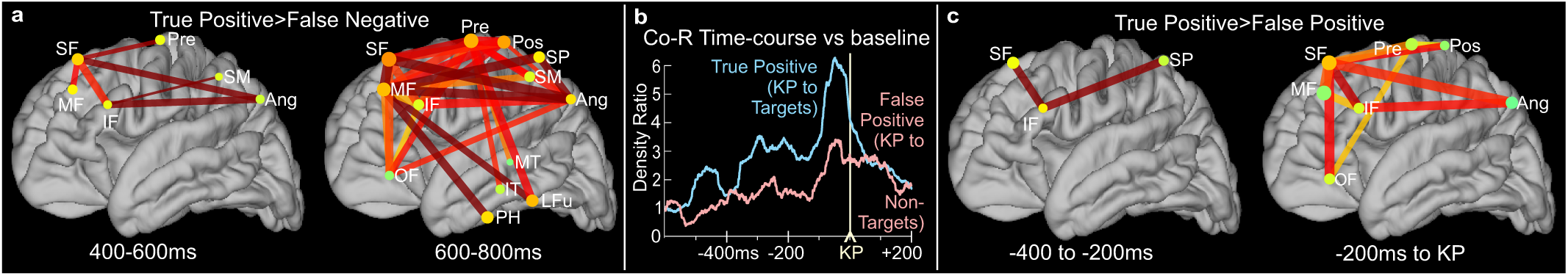
Response Accuracy. Correct keypresses elicit greater co-rippling than omitted (**A**) or wrong keypresses (**C**) mainly between semantic (SM, Ang), executive (SF, MF, IF) and response (Pre, Pos) areas. For legend see Fig. 3E. **B**. Co-rippling density averaged across all left hemisphere sites is larger prior to correct key-presses, compared to incorrect.

### Task-modulation of co-rippling is robust

Similar effects can be observed in individual subjects, and overall results are robust to analysis method, recording reference, and interstimulus interval. We repeated the analyses just described while varying experimental and analysis parameters and methods. In Fig. 5A, the pattern of increased co-rippling density to correct target vs. nontarget words from 600-800ms is reproduced from Fig. 3D, for comparison with subsequent panels 5B-H which each show the effects of changing a single parameter. In panel 5B, the number of co-ripples between the indicated areas rather than their density is used to test statistical significance and calculate ratios between conditions (please see Fig. S11 for a more complete presentation of latencies and conditions). Accordingly, the duration of target-evoked ripples did not differ from that of baseline ripples (Fig. S16). Similarly, although the observed target-evoked co-ripple counts did vary across trials and electrodes, their distribution was unimodal, suggesting that they were drawn from a single population (Fig. S17). In panel 5C (and Fig. S8), instead of analyzing monopolar LFPs recorded between cortical surface contacts and a distant extracranial reference, we analyzed the bipolar signals obtained by taking the difference of monopolar LFPs recorded from adjacent contacts located in the same anatomical parcel. In panels 5E and 5F (and Fig. S9) we analyze separately the task when the ISI was a fixed 600ms (7 participants) vs a random 1000-1400ms (4 participants, one took both versions of the task; Table S1). In panel 5G, we show the standard analysis applied to the data from a single subject. Additional examples of single subjects with more latencies and conditions are shown in Fig. S10. In all cases, the co-rippling distribution and timing is the same with the caveat that analyses involving fewer trials revealed fewer significant connections but nonetheless with the same overall pattern.

**Fig. 5.**
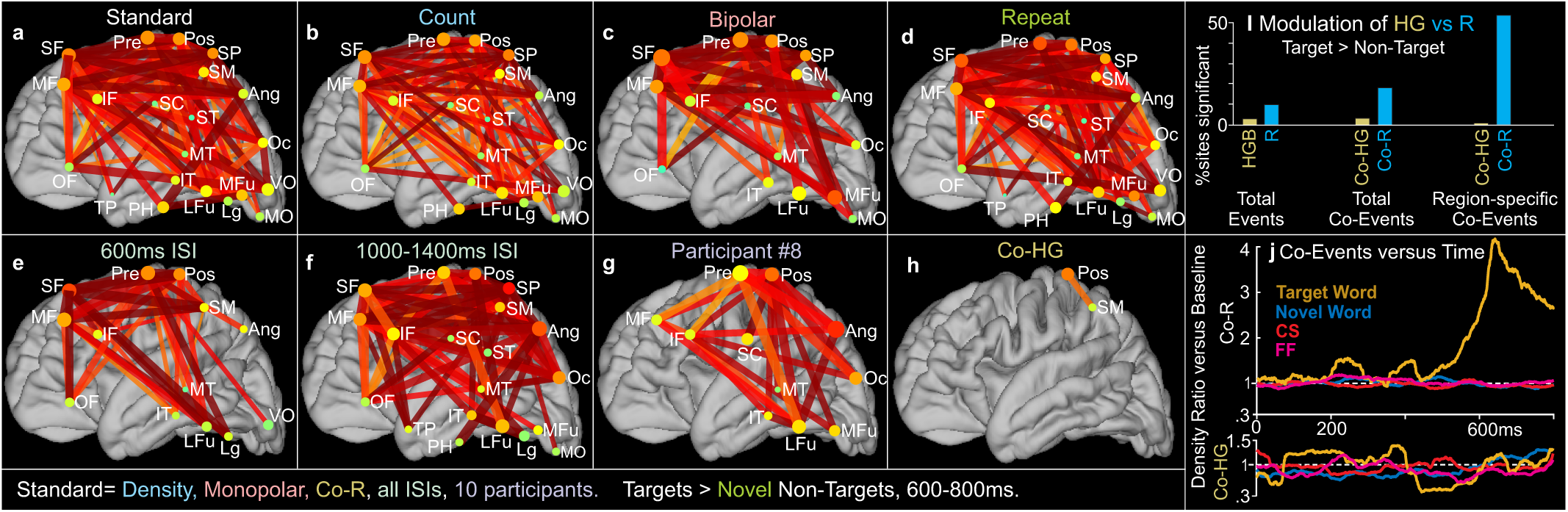
Comparison of Controls to Co-R Target Response. **A.** Co-ripple density to Target versus Non-Target words from 600-800ms post-word-onset, using monopolar recordings from 10 participants consolidated into regions of interest. Identical to the right panel of Fig. 3D, provided here for comparison to subsequent panels which are identical to A except: (**B**) the number of co-ripples rather than their density is plotted; (**C**) Ripples are detected in bipolar derivations rather than monopolar; (**D**); Targets are compared to Repeated Non-Targets rather than Novel Non-Targets; (**E**) only data from trials with a fixed 600ms ISI are used; (**F**) only data from trials with ISI randomly varying from 1000 to 1400ms are used; and (**G**) only data from a single participant are used. The similarity of the plots in A, B, C, E, F and G shows that the effects are robust to the manner of quantifying ripple co-occurrence (B), recording derivation (C), non-target repetition (D), and presentation speed (E,F), and are sufficiently robust to be observed in single subjects (G). **H**. Unlike Co-R (A), Co-HG in the same comparison change only minimally between conditions. **I**. Comparison of the percentage of sites or connections between left hemisphere sites that show significant Target>NonTarget effects for HGB and ripples. From left to right, pairs of bars show any target>non-target events, co-events, or region- and epoch-specific co-events (illustrated in panels A vs D). While HGB and Co-HG events occur and can differ between target and non-target words, they are much less sensitive to task modulation than Co-R, especially that specific for region and epoch. **J**. Time-course of co-event density to Target and Non-Target Words compared to baseline, summed across all sites, for Co-R (upper) versus Co-HG (lower).

### Novel and repeated co-ripple responses are almost identical

Binding wordforms to their meanings, or their meanings to task demands and response performance, are not hippocampus-dependent^38^, because they do not *require* reinstatement of a recent declarative memory trace. However, our task included repeated words which presumably formed a trace associating the word with the task context, and which could be reinstated when the words are repeated. If such reinstatement were to occur, it would be hippocampal-dependent due to the long distractor-filled delay between repetitions. Thus, if the co-rippling we observe is due to reinstatement then it would be selective for repeated words. However, repeated non-target words evoked essentially the same task-modulation of co-rippling as novel non-target words (Fig. 5D, S7; note that novel nontarget words were used for the standard comparison in Figures 3 and 5, because the targets were also all novel). Indeed, repeated non-target words evoked slightly *less* task-modulated co-rippling than novel non-target words, in a direct comparison (Fig. S7), or when comparing each with baseline (Fig. S3, S7). These effects are weak, with only one connection (medial fusiform gyrus – inferior temporal gyrus at 400-600ms) significant in the direct comparison. Nonetheless, this decrease is consistent with the oft-observed phenomenon of repetition-suppression due to facilitated lexical access and cognitive integration^39^, and thus provides further support for our hypothesis that such processes, rather than reinstatement, account for the cortical co-rippling we observed.

### Target-related non-oscillatory co-high-gamma bursts are rare

As described above, and shown in Fig. 2, we compared the task-modulation by words of co-HG to co-R. We also compared their task-modulation by target words. The contrast between co-ripples and co-HG bursts became much more pronounced when comparing the proportion of inter-areal connections that were significantly task-modulated. Increased connectivity to targets versus non-targets occurred in 111 inter-areal connections with co-ripples, compared to only 1 with co-HG (compare Fig. 5A to 5H, S3 to S5; Table 1). Similarly, the comparison of target words to baseline evoked no significant increases in Co-HG bursts between any region at any latency (Fig. S5). This was a general finding across task comparisons (Table 1). As indicated above, the lack of task-modulation of co-HG is not due to a lack of co-HG occurrences inasmuch as the mean and median density of co-HG were slightly higher than co-R (Fig. 2A). Furthermore, like ripples, more high gamma bursts were evoked by target than non-target novel words, although the number of left hemisphere sites where this increase was significant was much greater for ripples (14 versus 4%; Figs. 5I, S12). Similarly, more co-HG bursts were evoked by targets than non-targets, but this effect was again seen in more sites for co-ripples (29 versus 7%; Figs. 5I, S13), and overall co-HG density decreases before responses to targets, unlike the large increase seen in co-R density (Fig. 5J). Thus, while HG bursts and co-HG bursts do increase during reading, they are less specifically focused than co-R to the particular task conditions, latencies, and pairs of anatomical areas that are engaged by critical cognitive processes. These findings indicate that co-ripple effects are unlikely to be secondary to high gamma activation.

**Table 1:**
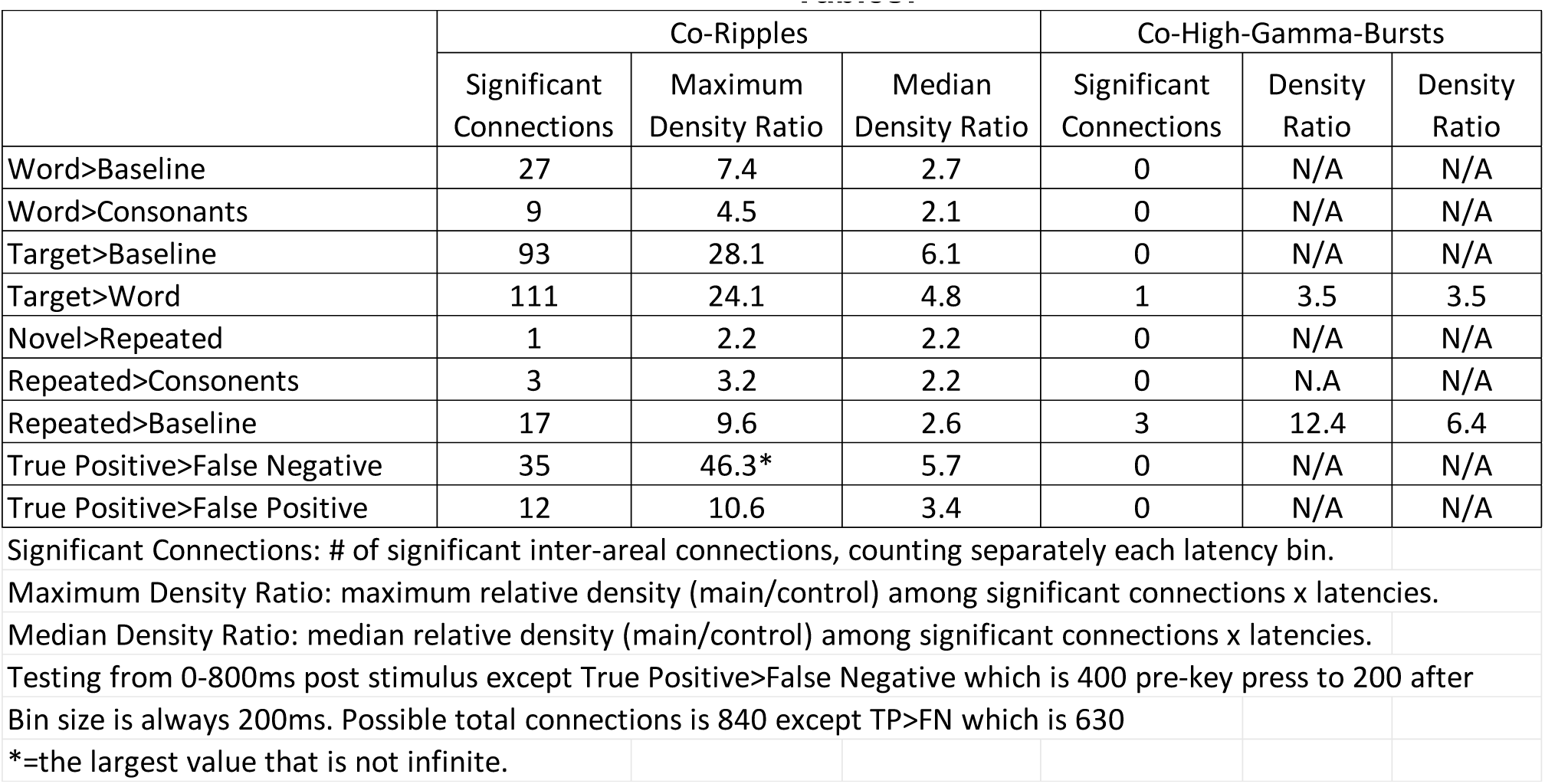
Co-Ripple and Co-High-Gamma-Burst Responses.

### Co-rippling sites phase-lock at zero phase-lag

If co-rippling between cortical locations facilitates binding of their encoding unit-firing patterns, then there should be indications of non-random interaction of the ripples in those locations. One such indication is the increased occurrence of co-ripples above what would be expected by chance (Figs. 2-5; Table 1). Another indication would be a tendency of such co-occurring ripples to oscillate with a consistent phase-lag relative to each other. The consistency of phase across co-ripples was quantified for each electrode pair as the Phase Locking Value (PLV). PLV was found to be significant between 89.0% of the 210 region-pairs (p<0.01, Raleigh’s circular uniformity test, FDR-corrected; Fig. 6A, Table S9), with the exceptions mainly in early visual areas (Figs. 6AC, S15). PLV amplitude was largely independent of the distance between recording electrodes (R^2^=0.009; b=-0.0076/cm; Fig. 6B), but increases with the proportion of recorded sites co-rippling, with PLV = 0.16 when ∼6% of sites co-rippling and increasing to 0.82 when ∼34% are co-rippling (Fig. 6FG). Thus, co-ripples between left hemisphere locations generally phase-lock with each other during the reading task, especially when a large proportion of the sites are co-rippling.

**Fig. 6.**
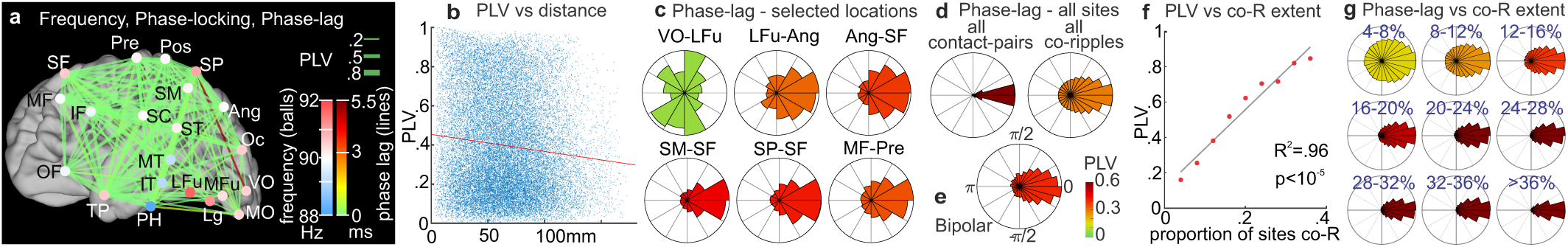
Widespread Left Hemisphere Phase-locking. **A**. Areas with significant PLV (Raleigh’s circular uniformity test, p<0.01, FDR corrected) during the task are connected with lines; line color indicates average phase-lag (±5.5ms =±π at 90Hz); dot color indicates average ripple frequency. **B**. PLV decreases slowly with distance, but distance explains only ∼1% of PLV variance. Each dot is an electrode pair with >20 co-ripples. b=-0.0075/cm [−0.0085, −0.0066], 5 R^2^=0.0085 p=10^−51^, t=-15.2, df=26869, two-tailed t-test. **C**. Phase-lag plots of all co-ripples between the indicated example structures (see also Fig. S15). Note the concentration of phase-lags near zero lag at the right side of the rosette, for all except VO-mFu (early visual sites have low phase synchrony with other areas). **D**. Phase-lag of all co-ripples (left) and average phase-lag for all significant contact-pairs (right, >20 co-R). **E**. Replication of phase concentration around zero-lag, when comparing bipolar derivations (contacts spaced 1cm apart on the cortical surface). **F**. PLV versus percent of contacts co-rippling. (p=10^−5^, b = 2.2 [1.8-2.6], t=13.5 df=7, two-tailed t-test) **G**. Phase of individual co-ripples concentrate more strongly at zero lag with increasing percentage of recorded contacts that are co-rippling. PLV is the vector-sum of phase-lags, ranging from 0 (no consistency) to 1 (perfect consistency).

The phase-lag between phase-locked oscillations can suggest the flow of information between them. The phase-lag between sites was significantly concentrated around zero in 97.8% of the connections with significant PLV (Figs. 6A, S15; Table S9). This is apparent in the phase histograms of all co-ripples between individual areas, or of all areas and contact-pairs (Fig. 6CD, S15). Zero-lag suggests that the communications between areas occurring during co-ripples are not being driven by one of the areas but are interactive.

The synchronization of distant locations as quantified by PLV and zero-lag is facilitated by the high consistency of ripple frequency across areas, whose mean frequency varies by a maximum of <2Hz (Fig. 6A), and individual ripples by ∼10Hz (Figs. 1F, Table S3). Across contact pairs, PLV was negatively correlated with difference in mean ripple frequency (p=10^−14^, b=-0.024/Hz [−0.030, −0.018], t=-7.9, df=5653, two-tailed t-test). Further, as more sites co-ripple, their ripple frequencies become more similar (R^2^=0.75; p=0.002, b=-8.5Hz [−12.5, −4.4], t=-5.0, df=7, two-tailed t-test Fig. S14). Thus, as co-ripples engage more sites, they tend to become more synchronous and oscillate at similar frequencies.

### Zero phase-lag is observed between bipolar derivations

While all possible phase-lags were observed between co-rippling sites, zero lag was clearly and consistently the most common, even at long distances. This is an important observation because it implies a network mechanism akin to coupled oscillators rather than sequential driving as the mechanism of phase-locking. It is therefore critical to confirm that the reported phase synchronization between two sites is genuine, and does not reflect artifactual common activity due to the common reference electrode, which in any case is unlikely because it was placed outside of the skull at a distance from the active electrodes which are lying on the pial surface. It is doubtful that brain activity in the 90Hz range can be recorded outside the skull, and we have previously found zero-lag phase-locking between ripples during waking in focal bipolar transcortical SEEG recordings, i.e. in the absence of a shared reference^28^. Furthermore, in the current study, we found that if a ripple occurs in a channel, the probability that any other given channel will ripple is <10% (Fig. 1M), whereas synchronous co-ripples from reference bleed would be close to 100%. Conversely, reference crosstalk could not reasonably account for the very high level of selectivity of co-rippling for particular epochs and between particular locations engaged in the task.

In addition to these considerations, we conducted a supplemental analysis with bipolar derivations where the common reference was eliminated as a factor. In this analysis, left hemisphere contacts were divided into pairs such that the distance between the contacts in each pair was 10mm (Euclidean distance), and ripples were identified in the potential difference between the contacts. We then identified co-rippling, phase-locking and phase-lag between all pairs of such contact-pairs, with the proviso that no two of the four contacts could be <40mm apart. Instances where ripples were present on both contacts of the bipolar pair were rejected, to assure that only one of the pair was ‘active.’ The phase-lags between all such pairs of contact-pairs clearly replicate the pattern of zero-lag phase we report in the referential recordings during co-ripples (Fig. 6E). In summary, the use of an extracranial distant reference electrode, the previous finding using bipolar transcortical electrodes of zero phase-lag, the low probability of co-rippling, the high task-modulation of co-rippling timing and location, and demonstration of zero phase-lag using local bipolar derivations in the current dataset, all strongly point to the zero phase-lag reported here representing the distant interactions of locally-generated activity.

## Discussion

In the current study we tested the hypothesis that co-ripples facilitate the integration (‘binding’) of information encoded in different cortical locations. This hypothesis predicts, and we found, that co-rippling is elevated in task conditions requiring binding, between cortical areas that encode information that needs to be bound, and at latencies when binding needs to occur. Our recordings revealed five epochs between stimulus and response (Table 2). In the first 200ms, all stimuli evoke co-HG bursts and atypical co-R in early visual areas, consistent with convergent visual processing using labelled lines in order to encode visual stimuli as wordforms. Local fusiform HG-bursts are followed by fusiform ripples. Beginning ∼150, and reaching peak amplitude at −240ms, all words evoke co-R between fusiform wordform areas and all other language-related areas, but not between those other areas themselves. This distributed co-R is larger to targets and sustained to the end of the trial. We hypothesize that it functions to distribute the encoded wordform for semantic processing. Starting ∼400ms, correct targets evoke an exponential rise in co-R centered on prefrontal cortex, which after ∼100ms becomes widespread between language-related areas. These strong distributed Co-R all peak at ∼640ms, except for Rolandic co-R which continue broadly until immediately preceding the response.

**Table 2:** Hypothesized Processing Stages.

| epoch (ms) | neurophysiological process | cognitive process | measure | location | task correlates | figures |
| --- | --- | --- | --- | --- | --- | --- |
| 40-150 | convergent visual processing using well-learned labelled lines | wordform encoding | co-HG, atypical co-R | early visual areas | all stimuli, esp. bizarre | 1K, 2DE |
| 150- 240 (first peak)- 800 | binding by synchrony | broad distribution of wordform | co-R | fusiform g. with all language areas | all words | 2C, 3A |
| 400- 640 (peak)- 800 | binding by synchrony | binding of semantic info with task requirements | co-R | prefrontal with all language-related areas | targets, esp. when correct | 3CDGH, 4AC |
| 500- 640 (peak)- 800 | binding by synchrony | contribution alignment | co-R | all language areas with each other | targets, esp. when correct | 3CDGH, 4AC |
| 500- 730 (peak)- 800 | binding by synchrony | response specification/ execution | co-R | Rolandic with language areas | targets, esp. when correct | 3CDGH, 4AC |

We find these observations consistent with initial convergent/sequential visual processing using well-learned labelled lines through the early visual hierarchy followed after ∼150ms through 730ms by sustained processing supported by BBS throughout (at least) the left hemisphere that integrates the wordform’s meaning with its context including target specification and response organization. This BBS-assisted processing has successive spatiotemporal stages from ∼150-730 focused on fusiform, prefrontal, all-language-related, and Rolandic areas These effects are robust to the manner of quantifying ripple co-occurrence, recording derivation, non-target repetition, and presentation speed, and are sufficiently robust to be observed in single subjects. Overall, our results support both types of global models for cortical integration, but suggests that they are dominant in different epochs and locations, and contribute to different essential cognitive processes, as well as relying on different neurophysiological mechanisms.

In the first phase of reading a word, lines are bound into letters, and letter-strings into wordforms. These processes are highly overlearned from millions of repetitions over decades. Extensive animal studies have delineated the hierarchical convergent activation of labelled lines that similar stimuli evoke, and human studies of reading are consistent with sequential processing culminating in the fusiform wordform area with responses selective for words (compared to false fonts and consonant-strings) shortly after 200ms post-word onset^16,40^. Due to clinical constraints, our sampling of early retinotopic areas was limited. In addition, the ripples we did record in these areas were sometimes unlike those in other areas, being larger, occurring more often, less phase-locked with other ripples, and with different task correlates and timing from those recorded elsewhere. This is important for distinguishing our results from most previous studies, because the binding of different sensory characteristics of visual stimuli is the most commonly studied form of binding in psychological experiments in humans^1^, as well as the most common model for studying BBS with neurophysiological recordings from the visual cortex of animals^26^. In our recordings co-R were relatively weak before 200ms, in stark contrast to the large sustained task-related increases for hundreds of ms after 200ms. Conversely, co-HG peaked prior to 200ms, with little or no task-related increases following 200ms. Furthermore, HG bursts led ripples in the fusiform when both occurred on single trials in the same contact. Thus, our recordings are consistent with previous studies supporting sequential convergent processing of labelled line activation for the initial well-learned steps of word processing prior to ∼200ms.

Beginning at ∼240ms, visual and auditory words evoke the N400 component of the EEG and MEG^41^. The N400 is attenuated if the current semantic or mnestic context facilitates the cognitive integration of the presented word. The visual N400 is absent to consonant-strings but is maximal to non-pronounceable nonwords, implying an antecedent process that encodes letter-strings as potentially meaningful and triggers an N400^21^. The N400 is widely distributed across the cortex^14,22,42^, as is semantic encoding^43,44^. Event-related intracranial potentials in a similar task also find that, following an initial sweep from visual to wordform areas, activation is simultaneously present in widespread cortical sites up to and beyond the behavioral response^22,45^.

The fusiform wordform encoding shortly after 200ms has been proposed to provide the ‘trigger’ marking the transition from sequential hierarchical labelled-line activation to widespread interactive processing which encodes the meaning of the word within its cognitive context^16,21^. Our findings provide strong support to this proposal in that, starting at ∼130ms, and peaking at ∼240ms, co-R increased between the fusiform and virtually all language-related areas. This occurred to all word categories, and was selective compared to consonant-strings.

Remarkably, the widespread areas co-rippling with the fusiform to words did not co-ripple with each other, unless the word was identified as referring to an animal, i.e., as a behavioral target. Beginning at ∼400ms, such words evoked co-rippling between language areas, peaking immediately before correct responses at ∼800ms, with the strongest modulation by far observed in this task. Initially, co-rippling to targets (beyond that with fusiform g.) are between parietal and prefrontal sites from ∼400ms, after which these expand to include co-rippling between most areas from 500, All peak together at ∼640ms. This suggests that the transition from the fusiform↔all_language to all_language↔all_language may grow out of co-rippling engaged when the semantic meaning of the word (parietal) resonates with the semantic definition of the response target in the task instructions (prefrontal). This possibility is supported by our observation that parieto-frontal co-rippling is also selectively increased prior to correct, compared to incorrect responses to target words, beginning in the 400 −600ms epoch, ∼400ms prior to the response. Since these are the critical interactions where binding is predicted to be necessary for correct performance, this finding also suggests that information integration during co-ripples contributes to successful task performance.

During this extended period of co-rippling between language sites, the ripples in different sites were strongly phase-locked at a phase that was indistinguishable from zero. This is most consistent with interactive processing rather than uni-directional information transfer between areas. Accordingly, in humans, cortical neurons in different locations reciprocally predict each others firing when their locations are co-ripples^30^. Given that cortico-cortical conduction times are ∼10m/s, the distances between co-rippling sites would result in phase-delays of >15ms at widely separated sites if communication were unidirectional. However, phase -delays between all sites were <1ms, and were not related to distance. The possibility that cortical synchrony is due to ripple-frequency driving by a subcortical structure is unlikely because phase-locking has not been observed between cortex and hippocampus^28^ or thalamus^46^. It is also unlikely that zero-lags were due to the shared distant extracranial reference because they were replicated using local bipolar derivations. Rather, computational modeling indicates that zero-phase gamma oscillations can arise in neural networks with dense interconnections^47^. The mechanism seems to involve bidirectional entrainment of ongoing activity by loosely coupled oscillators. Such a mechanism would be facilitated by the similarity of ripple frequency in different locations, and indeed, phase-locking was stronger when ripple frequencies were closer. Both higher phase-locking and similarity of ripple frequencies are strongly enhanced as a larger proportion of cortical sites are co-rippling, suggesting an interactive network mechanism that intensifies as more elements are engaged.

Several properties of the co-R reported here are inconsistent with the assertion that they are inconsequential byproducts of local activation^10,11^. The experiments supporting these assertions mainly involve early processing in vision cortex. Indeed, we found that a common marker for generic activation, non-oscillatory high gamma, tended to co-occur in visual areas at short latencies. However, overall, the co-activation of non-oscillatory high gamma bursts was much less related to task variables than were co-ripples, and never occurred during cognitive processing periods between semantic and executive areas. It is expected that ripples and high gamma would sometimes co-occur because ripples are associated with increased neuronal firing as well as increased high gamma during waking^27^. However, it is unlikely that the strong and specific association of co-rippling with functional areas and epochs which encode elements that need to be bound in our task is secondary to non-oscillatory high gamma co-activations which generally lack such associations. Furthermore, even if such associations were present, the facilitation of co-prediction during co-rippling by neurons in different cortical locations referred to above does not occur if firing increases without co-rippling^30^. Finally, the interpretation of co-R as local phenomena is inconsistent with the strong phase-locking they consistently demonstrate between distant structures.

The role of brief high frequency oscillations in behavior have also been intensively studied in the rodent hippocampus, where they occur on the peaks of sharp-waves during NREM sleep. These ‘sharp-wave ripples’ mark the replay of sequences of encoded spatial locations that were previously traversed during waking, and are critical for memory consolidation^48–51^. Ripples have also been recorded in the human hippocampus which are similar to those in rodents except for frequency^27,52,53^. Human ripples also occur during hippocampus-dependent memory retrieval during waking, and are associated with replay of previously learned firing sequences during both waking and NREM sleep^28,32–34,54^. Successful recall was associated not only with increased Rippling, but increased cortico-cortical and cortico-hippocampal co-rippling^28,33^. However, the task used in the current study did not require recent memory, and the task contrasts, latencies and structures associated with increased co-rippling were those expected for reading and semantic judgment, which would not be sensitive to hippocampal damage^38^. Conversely, a task manipulation (delayed repetition of some words) that rendered the word capable of reinstatement had very little effect on co-rippling. Furthermore, cortical ripples are more likely to co-occur, and much more likely to phase-lock, with ripples in other cortical areas than with hippocampal ripples^28^. Since memory is ubiquitous, it is not possible to definitively reject the possibility that the intense prolonged distributed co-rippling we observed was due to occult recall, for example of task instructions on each target trial. However, this seems less likely than that the co-ripples which we observed during reading reflect directly the task’s demands.

Our findings support predictions of the Global Neuronal Workspace (GNW) model wherein sequential-hierarchical focal processing eventually trigger a radiation of activation to widespread association cortex areas and sustained interaction ensues^55^. We suggest co-R as providing a mechanism supporting these interactions, noting that the entire brain is not co-R. From the mean ripple duration and density, total cortical pial surface area, and estimated size of functional modules, and noting that locations tend to co-ripple about twice as often as would be expected with transient increases to three times baseline, we roughly estimate 3,000-10,000 modules may be co-rippling at any given time (see Methods for details).

In summary, we show that ripples co-occur between early visual, wordform, semantic, executive and motor areas in the epochs and task conditions where binding between letters, words, meanings, instructions and actions need to occur for successful semantic judgements in a reading task. This supports a general role for co-rippling in binding cognitive elements encoded in different cortical areas. We found a sequence of neuronal activity patterns corresponding to different processes necessary to perform the task. In the first 200ms, limited evidence was consistent with sequential hierarchical convergent activation of labelled lines culminating in wordform encoding by the fusiform area. From ∼200-400ms, co-rippling connected fusiform with widespread language areas. These areas were not connected by co-ripples unless they were evoked by target words, when co-rippling between frontal and parietal sites from 400-600ms was followed by widespread co-rippling between nearly all language areas. Fronto-parietal and then Rolandic activation was further increased prior to correct responses from 600-800ms. Overall, our findings provide a framework for how BBS may support cortical integration during a cognitive task.

## Methods

### Participants and recordings

Subjects gave written informed consent to participate in this study, and the study was approved by the New York University Medical Center (NYUMC) and University of Californa’s Instituional Review Boards (UCSD IRBs) in accordance with the Declaration of Helsinki. Subjects were compensated hourly for their participation. Electrocorticographic (ECoG) recordings were obtained from 13 patients (10 females, mean age 36 (range 16-51), mean onset age 17 (range 3-38); patient information contained in Table S1) undergoing intracranial EEG monitoring to treat drug-resistant epilepsy. All patients were either right-handed, confirmed to have left-hemispheric language lateralization by Wada, or both. No patients exhibited clinically significant reading or language impairment. All procedures were approved by the Institutional Review Board at New York University and written informed consent was obtained from all participants. Electrode placement was determined by clinical criteria with the objective of identifying the seizure onset zone and eloquent tissue. Each patient was implanted with subdural platinum-iridium electrode arrays embedded in silastic sheets (AdTech Medical Instrument Corp). Data included arrays of grids (8−8 contacts) and strips (1−4 to 1−12 contacts). Contacts had a diameter of 4mm with 2.3mm exposure. Center-to-center spacing between contacts was 10mm for grids. In total, 8 patients had predominantly left-sided implantations, 4 patients had predominantly right-sided implantations, and 1 had bilateral implantations (i.e., strips distributed around both hemispheres). Connection plots included data from all 9 subjects with left-hemispheric sampling (Table S1). All other plots not specified as left-hemispheric included data from all 13 subjects. Recordings were acquired using a Nicolet One EEG system sampled at 512 Hz (i.e., a temporal resolution of 1.95 ms) and bandpass filtered between 0.5 and 250 Hz.

### Semantic judgment task

(Fig. 1A). Stimuli were white on a black background, in Arial font, with an approximate visual angle width of 4°, comprising novel real words (N), previously presented ‘repeat’ words (R), non-pronounceable consonant letter strings (CS), and target words (T). False font stimuli (FF) were not analyzed in the current study except to characterize outlier early visual ‘ripples.’ All stimulus categories were balanced in the average number of characters. Subjects pressed a button in response to target words representing animals. Target words were relatively infrequent in the task, representing ∼5% of all words and ∼9% of all stimuli. N, R and T were 4-8 letter nouns, with a written lexical frequency of 3-80 per 10 million^56^. There were 10 R words, each repeated 20 times. Repetition of a given word was separated by at least 15s (median 25s) and 24 (median 42) intervening trials. Tasks were programmed using Presentation software (Neurobehavioral Systems, Inc).

In each trial block we presented 200 each N, R, CS, and FF, and 40 T pseudo-randomly with the constraint that each condition was preceded by every other condition with equal likelihood. Subjects completed 1-4 trial blocks (median 2). Two versions of the task were used, differing only in inter-stimulus interval (600 or randomly from 1000-1400 ms). Throughout the experiment, each CS and FF stimulus was only presented once. Subjects detected 80.6% (stdev: 10.0%) of the targets (chance = 9.1%) range 62.7% to 94.9%. Subjects d’ was 3.37 (stdev: 0.73), range 2.23 to 4.49. Feedback was not provided. The mean false alarm rate was 1.3% (standard deviation of 1.8%, range 0.1-6.9%). For the set of subjects used for True Positive > False Positive contrasts, the mean false alarm rate was 2.2% (standard deviation 2.4%, range 0.6-6.9%). The mean reaction time was 789ms (stdev of subject means: 56 ms). This calculation excluded RTs >1500 ms. Since the Reaction Time often exceeded the inter-stimulus interval, the trials following targets were excluded from physiological analysis.

### Electrode localization and regions of interest

Electrode localization was done through co-registration of pre- and post-implant MRI images, followed by manual and automatic localization of electrodes^57^. Three-dimensional reconstructions of cortical surfaces in figures were created using FreeSurfer^58^. Anatomical parcellations (Table S2) were determined in most cases by using a modified Destrieux atlas^59^ performed in each subject’s native brain. In addition, the V1 and V2 parcels as defined in the HCP-MMP1 parcellation^60^ were used to place early visual cortex within the Medial Occipital Area. The fusiform gyrus was divided into medial and lateral portions, following the HCP parcellation based on their very distinct activation and myelination profiles^60^. For example, the lateral portion mainly corresponds to the Region termed FFC (fusiform face contrast)^60^ and also includes a portion of area TF that lies on the anterolateral fusiform gyrus in the HCP atlas*. All regions were sampled by at least 3 subjects and 8 electrode contacts*.

### Data processing

Data were preprocessed using MATLAB (MathWorks), the Fieldtrip toolbox^61^, and custom scripts. We applied a bandstop around line-noise and its harmonics (60,120,180Hz). Power spectra were generated for each subject, and narrowband noise phenomena were identified and removed using narrow bandpasses (bandwidth/base frequency range 0.0002-0.01). Data were epoched from the onset of stimulus presentation for 1000 ms, or relative to response events from −400 to 200 ms. Trials containing artifacts were identified by outlier amplitude, variance, z-score, or kurtosis in broadband LFP or high gamma (70-190 Hz) amplitude. For subjects P6-P8, independent components analysis as implemented by the Fieldtrip toolbox was used to identify and remove gross artefactual components, corresponding either to electronic noise or movement artifacts. Components were extracted using either Infomax^62^ or FastICA^63^ algorithms. Components were removed only if clearly artifactual and lacking physiological contribution.

### Channel selection

Channels were excluded which were identified as seizure origins, early sites of seizure spread, or which displayed substantial interictal epileptiform activity (See section “Rejection of ripples possibly related to epilepsy”). Channels were excluded if significant artifacts were present during the task period. For the baseline comparison connection plots only (Fig. 4) we also eliminated channels with very high ripple density (>30 ripples/min) in regions VO (6 sites) and MO (4 sites). VO and MO were clearly different from other areas in their amplitude, density, and lack of phase-locking (Figs. 2d, 5AC, S15; Table S9). In particular, these areas contained channels with high ripple densities that were evoked by all visual stimuli, potentially resulting in high levels of coincidental ripple overlap, as well as resulting from non-representative ripples. Similar issues did not arise in the condition comparison plots (e.g., Fig. 3) because the visual-evoked activity was present in both conditions.

### Time-frequency analyses

Average time-frequency plots of the ripple event-related spectral power (ERSP) were generated from the broadband LFP using EEGLAB^64^. Event-related spectral power is calculated from 1Hz to 200Hz with 1Hz resolution with ripple centers at *t*=0 by computing and averaging fast Fourier transforms with Hanning window tapering. Each 1Hz bin was normalized with respect to the mean power at −2000 to −1500 ms.

### Ripple (R) detection

was based on a previously described method^27,28,52,53,65^. Z-scores of the 70-110Hz bandpassed analytic amplitude, smoothed using a Gaussian kernel with a standard deviation of 20 ms, were computed on a channel-by-channel basis. R events were selected by identifying regions of the recording that exceed a z-score of 2.5 relative to a pre-stimulus baseline and contain at least 3 distinct oscillation cycles of relatively consistent amplitude in the 70-100Hz signal, determined by zero crossings. R centers were determined as the time of the maximum positive peak. In order to ensure that transient phase-shifts do not inappropriately terminate R-detections, a two-step algorithm was used to compute R start- and end-points. Approximate boundaries were determined using the time that the Z-score of the 70-110 Hz bandpassed epoch, smoothed using a Gaussian kernel with a standard deviation of 36 ms, drops below a value of 1. Then, precise boundaries were determined by applying the same criterion to the bandpass smoothed using a Gaussian kernel with standard deviation 20 ms. Ripples occasionally display transient drops in amplitude lasting less than one cycle in otherwise strong oscillations. Visually, these appear to be shifts in phase, rather than true changes in ripple-band activity. The 36 ms kernel was chosen empirically as it is relatively insensitive to these phenomena, and detected approximate ripple starting and ending points that agreed with examination of the raw LFP. The second stage using the 20 ms kernel is then applied to find the precise boundaries of the ripple within the approximate ranges determined by the 36 ms kernel. The 20 ms kernel size was also chosen empirically, as it reliably selected the edges of the first and last cycle of the ripple. This approach of using different kernels for first identifying a graphoelement and then determining with greater precision its onset and offset has been used several times in our laboratory successfully, as applied to sleep spindles^53,66,67^.

For each channel, the mean R-locked LFP and mean R bandpass were visually examined to confirm that there were multiple prominent cycles at R frequency (70-110Hz) and the mean time-frequency plot was examined to confirm there was a distinct increase in power within the 70-110Hz band. In addition, multiple individual R in the broadband LFP and 70-110Hz bandpass from each channel were visually examined to confirm that there were multiple cycles at R frequency without contamination by artifacts or epileptiform activity. Sites in early visual regions (Medial Occipital and Ventral Occipital) often displayed R properties unlike other regions, as observed in terms of R amplitude and rate (Fig. 1HI). Early visual sites included a subpopulation which exhibited R densities consistent with the distribution of values observed elsewhere, and another with R densities which were substantially higher than expected (Fig. 1K). Therefore, sites in early visual regions which displayed a z-score R rate of greater than 2 (corresponding to 30 R/second) were excluded from further analysis.

### Rejection of ripples (R) possibly related to epilepsy

Epileptiform activities or artifacts were excluded if the absolute value of the unfiltered data in two successive samples exceeds 40µV, or 3 successive samples 60 µV. They were also excluded if the absolute value of the unfiltered data at any point in the R exceeds 500µV. R were also excluded if they fell within ±500 ms of putative interictal spikes, detected with templates developed for each channel individually and visually confirmed. To exclude events potentially coupled to epileptiform activity on another channel, we exclude R that coincided with a putative interictal spike on any cortical or hippocampal channel included in the analyses. Events were also excluded if the largest absolute amplitude excursion in the broadband LFP was 2.5 times greater than the third largest excursion.

### High-gamma-burst (HGB)

detection uses a modified version of the method used to detect ripples. LFP is run through a 120-190 Hz bandpass filter, and candidate events are selected by identifying time periods where the z-score of the Hilbert analytic amplitude exceeds 2. Candidate events are further selected for non-oscillatory activity: cycle lengths are measured on the LFP waveforms bandpassed at 100-200 Hz, and events included only if the standard deviation of cycle lengths exceeds 1.1 ms.

### Co-ripples (Co-R) or co-high-gamma-bursts (Co-HGB)

were defined as ripples or high gamma bursts occurring in two channels with an overlap of at least 25 ms. The criterion of 25 ms was chosen to ensure that ripples overlapped for at least two full cycles. Co-R density was defined as the total number of time-points occupied by Co-R, summed across subjects, channel-pairs, and trials, divided by the theoretical maximum of this value. For a given connection between ROIs, all possible pairings between the two relevant sets of channels were included in calculation of density. Where *p, c,* and *t* denote the index of participant, channel-pair, and trial, respectively, and *C_p_* and *T_p_* denote corresponding totals in participant *p*, the total quantity of observed Co-R was defined as:

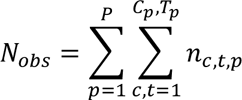

where *n_c,t,p_* denotes the number of time-points occupied by Co-R in the specified channel-pair and trial. The theoretical maximum of *N* is calculated:

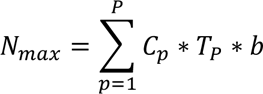

where b denotes the number of time-points in the latency window considered. Density is simply the ratio:

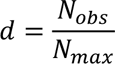

The same procedure was applied to quantify Co-HGB density. The analogous procedure calculated over individual channels rather than pairs was used to calculate ripple and high gamma burst density.

### Connection plots

Channels were pooled into regions, with regions assigned as described above. A co-event between two regions was defined as a co-event occurring between two channels, with one in each region. For each contrast, the ratio of (co-)ripple densities for each region(-pair) was calculated, and tested against a null distribution generated by randomizing trial identities (described in detail below). P-values thus generated were corrected over all region-pairs, subjects, and latency bins (200 ms bins from 200 ms to 1000 ms post-stimulus, or from −400 ms to 200 ms relative to task responses, FDR correction, alpha=0.01), and only significant connections are shown. For baseline contrasts, each latency was tested relative to the −100 to 0 ms pre-stimulus interval, in 100ms bins from 0 to 1000 ms post-stimulus. All stimulus types were included in the baseline; however, trials which followed a target stimulus were excluded. Density values described above were used to determine the visual width of lines and nodes in connection plots, and the ratio of density values in the experimental condition and control condition was used to determine coloration. Plots were displayed over the pial surface of the fsaverage template brain, projected orthographically on a view 20° anterior and 30° inferior from lateral.

### Reverse contrasts

All of the above comparisons were made single-tailed, active condition vs control, because these were the only ones of interest, and the only ones expected to occur. Nevertheless, for completeness and to fully characterize our data, reverse contrasts (e.g. Novel NonTarget>Target, Baseline>Repeat) were tested for both co-ripples and Co-HGB, using the same methods as described above. These contrasts include 16 condition comparison time-courses, or a total of 5,880 co-ripple area-area comparisons by latency bins, and an equal number of co-HGB comparisons. Of these, only 1 total significant region-to-region co-event density decrease was observed (i.e., 5 of 11,760), found in only 4 of the 16 time-courses. Novel showed 1 and Repeat words showed 2 significant co-ripple density decreases relative to baseline, Repeat words showed 1 significant Co-HGB density decrease relative to baseline, and Novel words showed higher Co-HGB density than Target words for 1 region pair.

### Ripple (R) density or high gamma burst (HGB) contrasts

by region were calculated using a procedure analogous to that described above (Methods: Co-ripples (Co-R) or co-high-gamma-bursts (Co-HGB)). Briefly, to calculate R or HGB density for a given region, the durations of all R or HGB which occurred in any relevant channel during the analyzed condition and latency were summed. This was then divided by the theoretical maximum of this value, which was the length of the latency window multiplied by the number of such channels and trials, summed over each participant. As with Co-R and Co-HGB, condition contrasts were the ratio of density in the main condition to that in the control condition. R and HGB density values are displayed in Connection Plots (Fig. 2CD, 3A-D, 4A,C, 5A-H, S1-11) and unlike Co-R and Co-HGB, the R and HGB density values were not statistically tested or their display thresholded; rather, values were displayed for all regions involved in significant Co-R or Co-HGB density contrasts.

### Co-ripple density timeseries between zones

(Fig. 3E-H) were calculated using the same methods detailed above, with the modification that co-ripple density was quantified at each time-point (ms), and regions were amalgamated into Zones in order to obtain sufficient SNR. Zones were composed of ROIs used in connection plots, as follows: Fusiform: MFu and LFu; Mid. Temporal: MT; Prefrontal: SF, MF, and IF; Parietal: SP, SM, and Ang; pFrPar: SF, MF, IF, SP, SM, and Ang; Rolandic: Pre and Pos.

### Bipolar re-referencing

for task modulation of co-ripple density analyses employed the set of high-quality referential channels and artifact-free trials used for all other analyses. The contacts forming each bipolar pair spanned no more than 30mm, and were located within the same region of interest. Ripples were detected on the resultant channels using the same methods described above. This bipolarization scheme reduced the number of contacts from 781 to 558, and the number of co-ripples from 113,276 to 79,214. The above procedure pertains only to co-ripple density analyses shown in Figs. 5C and S8.

### Bipolar re-referencing for phase-locking analysis

(Fig. 6E) was also conducted on the set of high-quality referential channels, but used an alternate bipolarization scheme. In this case, all contact-pairs forming bipolar channels were 10mm apart, and all pairs of bipolar ‘*channels’* used for phase analyses were separated by at least 40mm (minimum Euclidean distance between any two constituent contacts). Finally, ripples were excluded if present in both referential channels constituting a bipolar pair, so as to avoid the possible consequent phase-shifting the ripple-band signal. Polarity was assigned to bipolar pairs such that for each co-ripple, the positive poles of the bipolar pairs always corresponded to the contacts at which ripples were detected in the referential data. The total number of bipolar channels in this scheme was 318, and the total number of analyzed channel pairs was 11,728, vs. 33,787 referential channel-pairs.

### Site-wise analysis

For plots of significance by single site (Figs. S12 and S13 only), R or HGB density was calculated as above for each site separately in each latency bin. For each site, Co-R density was calculated for Co-R with any other site. Densities were tested using the same permutation testing procedure described below (Methods: Statistical analyses). P-values were corrected across sites and latency bins, and sites were labeled significant if p<0.01 in any latency bin.

### Ripple phase-locking value (PLV) analyses

PLV is a measure of phase-locking^68^, which unlike coherence is not affected by shared amplitude modulation. PLV was computed using the analytic angle of the Hilbert transformed 70-110Hz bandpassed (zero-phase shift) signals of each channel pair when there were at least 20 co-ripples with a minimum of 25 ms overlap for each. In order to stabilize phase estimates, the difference in ripple phase angles in radians was calculated over the duration of each ripple overlap period, then converted to complex phase values and averaged, to compute a single phase-delay value associated with each co-ripple event. Except where otherwise noted, phase distributions and PLV estimates presented are derived from the distributions of these stabilized phase-delay values, which correspond one-to-one with co-ripples. For such a distribution, PLV is calculated as the magnitude of the vector mean.^68^ A similar approach has been used previously by our group to demonstrate phase-locking in co-ripples^28,69^. PLV significance was calculated using FDR correction and a uniform null distribution (Raleigh’s circular uniformity test)^70^ of the angular difference between the band-passed ripples during their overlap.

The PLV and number of channel-pairs (#CP) or co-ripples (#Co-R) for the panels of Figure 6 were as follows: Panel C: VO-lFu: PLV=0.06 #Co-R=92, lFu-Ang: PLV=0.27 #Co-R=408, Ang-SF: PLV=0.37 #Co-R=2610, SM-SF: PLV=0.45 #Co-R=2901, SP-SF: PLV=0.42 #Co-R=1947, MF-Pre: PLV=0.34 #Co-R=5457. All except VO-mFu have both significant PLV and zero-latency phase locking at p<0.01. Panel D left: PLV=0.95, #CP=4021; right: PLV=0.31, #Co-R=615441. Panel E: PLV=0.95 #CP=2161. Panel G: 4-8% PLV=0.29, #Co-R=448329; 8-12% PLV=0.26, #Co-R=75125; 12-16% PLV=0.38, #Co-R=46180; 16-20% PLV=0.52, #Co-R=24706; 20-24% PLV=0.63, #Co-R=20634; 24-28% PLV=0.71, #Co-R=17569; 28-32% PLV=0.72, #Co-R=18814; 32-36% PLV=0.82, #Co-R=11263; >36% PLV=0.85, #Co-R=7559; (all significant at p<0.001).

### Statistical analyses

Ripples and co-ripples relative to behavioral events were compared between task conditions. Due to inherent correlations in co-rippling as a bivariate measure, we employed permutation tests to assess significance. In connection plots and site-wise plots, our measure of interest was ‘relative density’, defined as the proportion of time occupied by co-ripples in the main condition divided by that in the control condition. We tested these observed values against a distribution of relative density values which would be expected if the task condition did not influence co-ripple density. This distribution was generated by resampling the data while randomizing the task condition between the main condition and control, thereby computing hypothetical relative densities. This was performed 10,000 times to generate a null distribution with as many samples for each comparison (contrast x region x region x latency). Thus, this procedure provides the expected distribution of relative density under the assumption that the main and control conditions produce the same density, and therefore the likelihood that the observed difference in densities would occur due to chance. Analogous procedures were used to test for significance of region-to-region co-high-gamma-burst density modulation, and also for site-wise modulation of (co-)ripples or (co-)high-gamma-burst density. FDR corrections were applied across channels or region-pairs and latencies, as appropriate. All statistical tests were evaluated with α=0.01. All p-values involving multiple comparisons were FDR-corrected^71^. FDR corrections across channel pairs were done across all channels pairs from all patients included in the analysis. Category and order distributions were tested with two-sided binomial or χ^2^ tests. To determine if a circular distribution was non-uniform, the Rayleigh test was used. Zero-latency phase locking was tested using a right-tailed binomial test of the proportion of co-ripple phase angles in the interval [−π/8, π/8] against a null proportion of 1/8.

### Calculation of number of cortical modules co-rippling

We used the mean ripple duration (113.5ms) and mean density (18.6/min) from this paper to estimate that a given location is Rippling 3.5% of the time. We used typical literature value for total cortical pial surface area = 2.18e5 mm^272^, and a 1.7mm estimate of functional module diameter in human superior temporal gyrus from microgrid recordings^73^, to estimate the approximate number of cortical modules (∼96,000) and the average number of co-rippling locations (3,381). Overall, locations tend to co-ripple about twice as often as would be expected (Fig. 1I), and transient increases to three times baseline (Fig. 1L). We thus arrive at a rough estimate of 3,000-10,000 modules co-rippling at any given time.

## Data Availability

Data underlying the present study including electrode localization, ripple detections, phase data, and other ripple characteristics are publicly available on Zenodo (density analyses: https://zenodo.org/records/8396957, phase analyses: https://zenodo.org/records/12520199). Imaging and raw intracranial recordings are available upon reasonable request to the corresponding author.

## Code Availability

MATLAB code underlying the present study are available on Zenodo, including co-ripple density analyses (https://zenodo.org/records/8396957) and phase-related analyses (https://zenodo.org/records/12520199). Code implementing previously published methods for ripple detection is available on GitHub (https://github.com/iverzh/ripple-detection).

## Acknowledgments

We thank Charles Dickey, Burke Rosen, Leo Breston, Eran Mukamel and Sophie Kajfez for their support. Funding: National Institutes of Health grant MH117155 (EH); National Institutes of Health grant T32MH020002 (JCG); Office of Naval Research grant N00014-16-1-2829 (EH)

## Author Contributions Statement

Conceptualization: EH; Data Curation: JCG, EK, IAV, TT; Formal analysis: JCG, IAV, EH; Funding acquisition: EH, TT, OD; Investigation: TT, CC, WKD, OD, EH; Methodology: JCG, EH; Project administration: EH, TT, OD; Resources: OD, WKD, CC; Software: JCG, IAV; Supervision: EH, TT; Visualization: JCG; Writing – original draft: EH, JCG; Writing – review & editing: EH, JCG

