## Supplementary Materials for "Binding of cortical functional modules by synchronous high frequency oscillations"

**The PDF file includes:**

Supplementary Figures


S1. Ripple characteristics by patient.

S2. Exponential increase in Prefrontal-Parietal co-ripple density following Target stimuli.

S3. Spatiotemporal evolution of co-ripple density evoked by Novel Words compared to Baseline or Consonant Strings, and Target words compared to Baseline or Novel Nontarget Words, with 200ms bins.

S4. Spatiotemporal evolution of co-ripple density evoked by Novel and Target words, with 100ms bins.

S5. Co-high-gamma burst density is not modulated in main contrasts and epochs.

S6. Selective executive-semantic co-ripple engagement during correct task performance.

S7. Repeated and Novel nontarget words evoke largely equivalent co-ripple density patterns.

S8. Major task effects are robust to bipolar re-referencing.

S9. Spatiotemporal patterns of co-rippling are consistent between trials with 600 ms and 1000-1400 ms inter-stimulus intervals. A. 600ms duration trials in 7 subjects. B. 1000-1400ms duration trials in 4 subjects. C. 600ms duration trials in subject P8. D. 1000-1400ms duration trials in subject P8.

S10. Major co-ripple density modulations resolve within individual subjects. A. Subject P1. B. Subject P4. C. Subject P7. D. Subject P8.

S11. Quantification employing counts of co-ripple or co-high-gamma burst events yield similar results to density-based analysis. A. co-ripples. B. co-high-gamma bursts.

S12. Ripple density and high gamma burst density show limited site-wise modulation by Target words.

S13. Site-wise co-ripple density modulation by Target words is more widespread than that of co-high-gamma bursts or ripples.

S14. Widespread co-rippling is associated with more consistent ripple frequency across sites.

S15. Co-ripple phase distributions by region pair.

S16. Target-evoked co-ripple durations do not differ from Baseline.

S17. Distributions of Target-evoked co-ripple occurrence across both trials and channels displayed unimodal distributions.


Supplementary Tables


S1. Patient characteristics.

S2. Regions of interest.

` S3. Ripple characteristics by region of interest.

S4. Region-region co-ripple relative density in Novel>Baseline contrast, 200 – 400 ms post-stimulus.

S5. Region-region co-ripple relative density in Target>Baseline contrast, 400 – 600 ms post-stimulus.

S6. Region-region co-ripple relative density in Target>Novel contrast, 600 – 800 ms post-stimulus.

S7. Region-region co-ripple relative density in True Positive>False Positive contrast, -200 – 0 ms peri-response.

S8. Region-to-region co-ripple counts.

S9. Region-to-region co-ripple PLV and zero-latency bias.

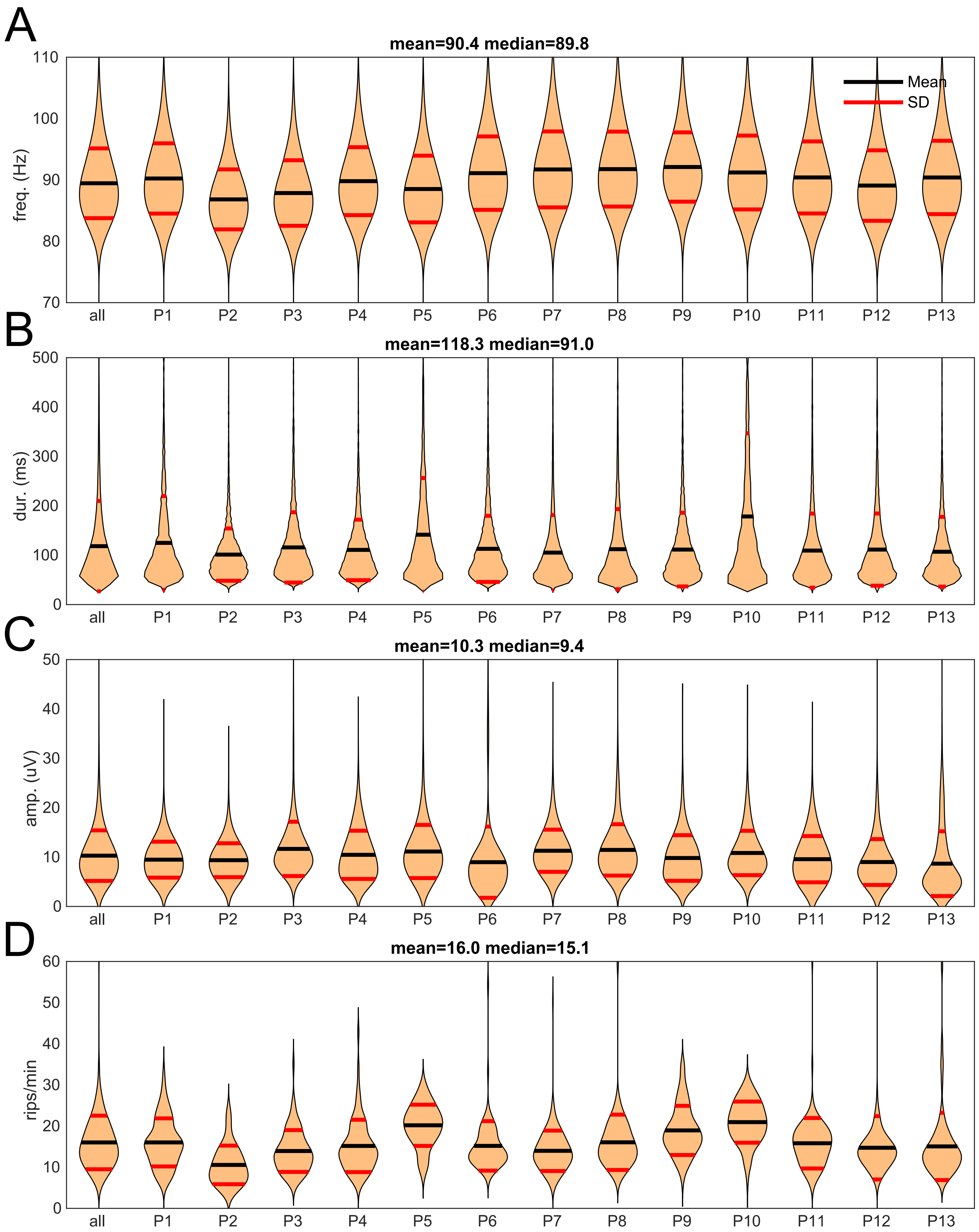


**Fig. S1. Ripple characteristics by patient.**

**A**. Frequency distribution of Ripples, by patient. Frequency was measured by the spacing of zero crossings in 70-110 Hz bandpass. **B.** Duration distribution of ripples. **C.** Peak amplitude distribution of ripples, as baseline-to-peak of the Hilbert analytic amplitude. **D.** Ripple rate distribution of channels.

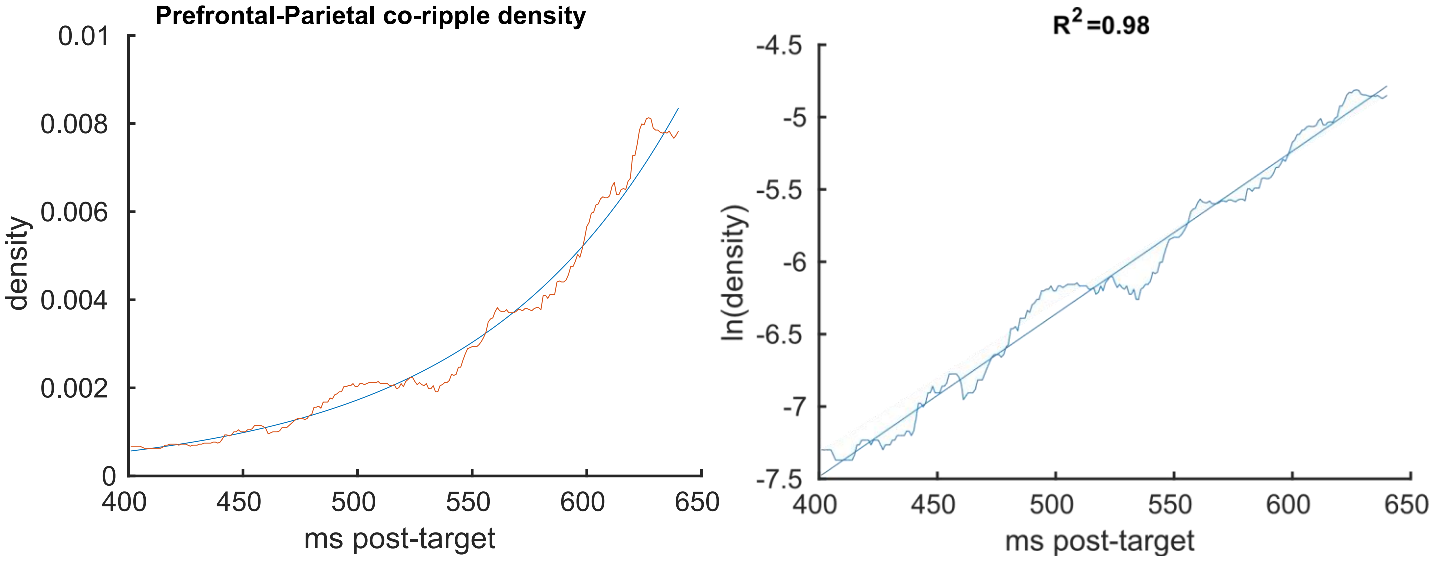


**Fig. S2. Exponential increase in Prefrontal-Parietal co-ripple density following Target stimuli.** Left: Prefrontal-Parietal Co-R density (red) with exponential fit-line (blue). Right: Log-transformed data and corresponding linear fit-line. Fit lines are equivalent; fit parameters and $R^2$ (adjusted) were derived from ordinary least squares regression of log-transformed data (coefficient: .0.113 [0.111, 0.114]).

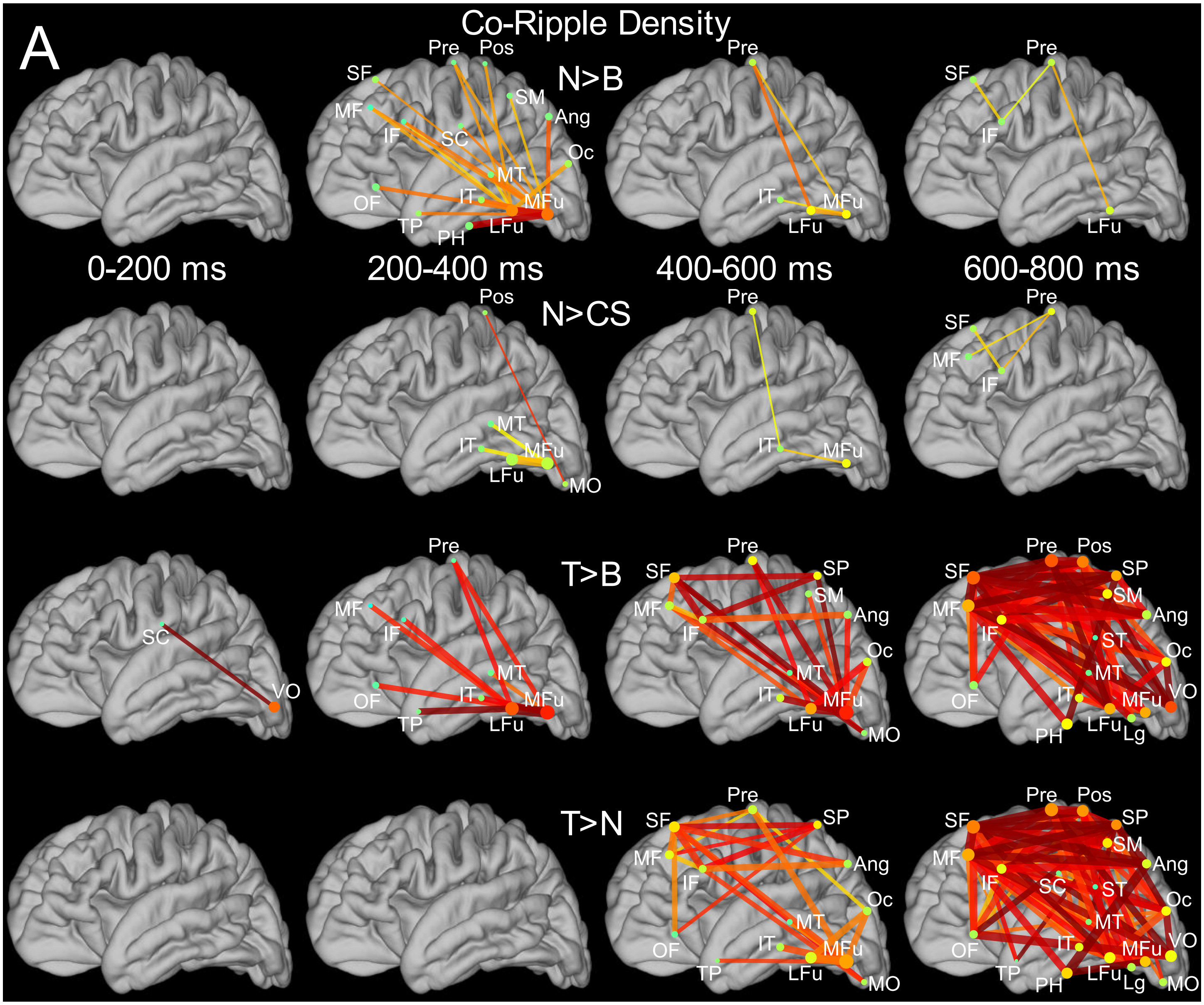


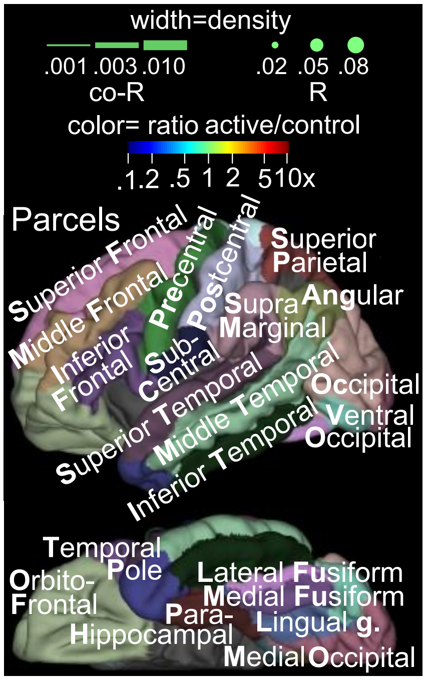


**Fig. S3. Spatiotemporal evolution of co-ripple density evoked by Novel and Target words: Main contrasts and epochs of analysis.** Each row shows connections where the Co-R density was significantly higher in the indicated main condition relative to the control condition (stimulus-type permutation test, $p<0.01$ FDR corrected). Columns correspond to each 200 ms latency window from stimulus onset to 800 ms post-stimulus. Condition B denotes the 100 ms pre-stimulus common Baseline, N non-target Novel words, CS Consonant Strings, and T Target words. Density values represent the sum-total time occupied by Co-R over channel-pairs, trials, and subjects, relative to the theoretical maximum value. See **Materials and Methods, Co-ripples or co-high-gamma bursts** for more detail. Identical to **Fig. 3A-D** in main text. At right is the plot scaling for all such 'connection plots' (including **Figs. S3-S11**) and locations of regions of interest (ROIs). See **Table S2** for further detail on specification of ROIs. **Note that the identical comparisons and analysis but conducted on co-high-gamma bursts rather than Co-R density revealed no significant effects** (Fig. S5).

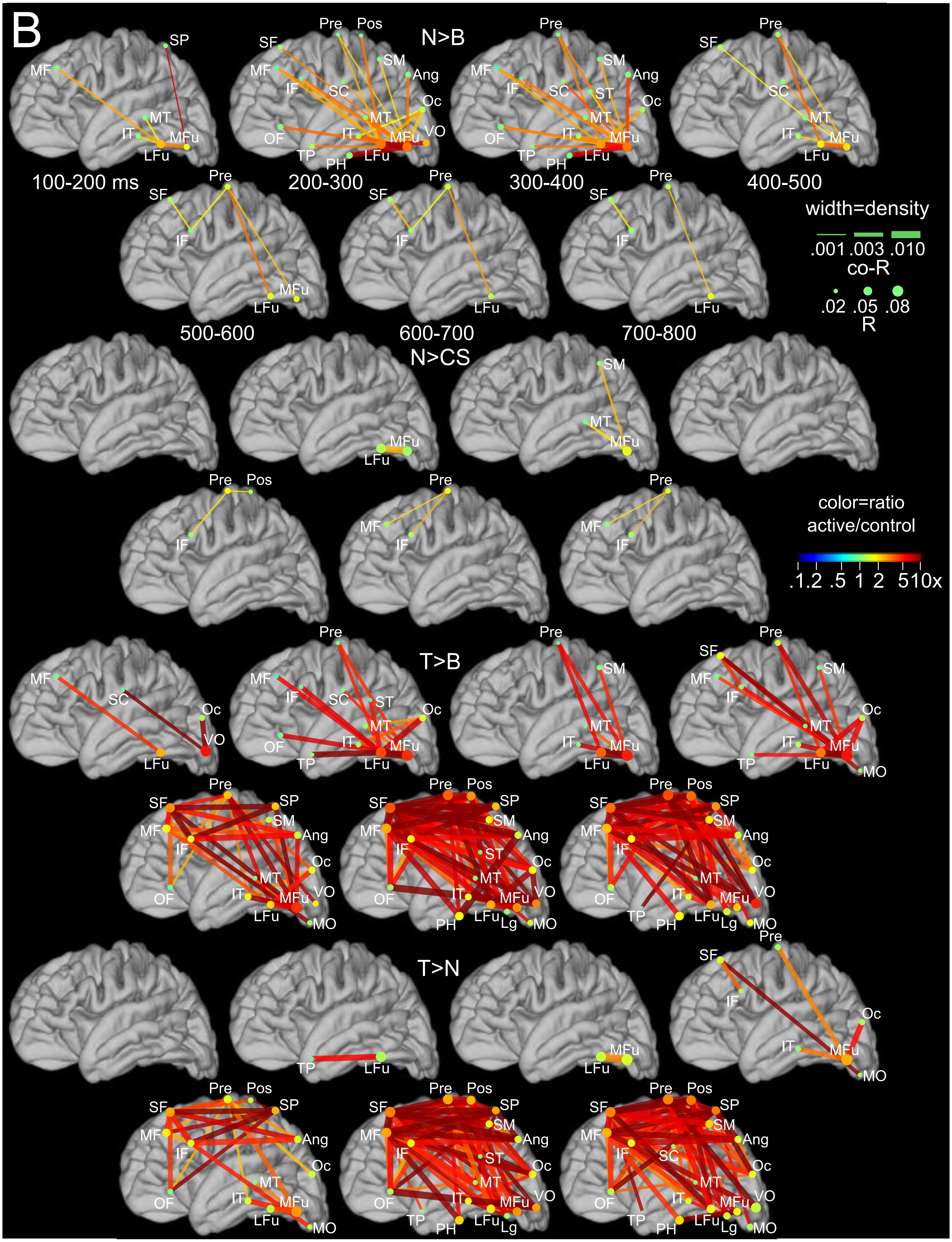


**Fig. S4. Spatiotemporal evolution of co-ripple density evoked by Novel and Target words: 100ms latency windows.** Identical in all respects to Fig. S3 but with seven 100ms long analysis windows from 100-800ms post-word onset, rather than four 200ms long windows from 0-800ms.

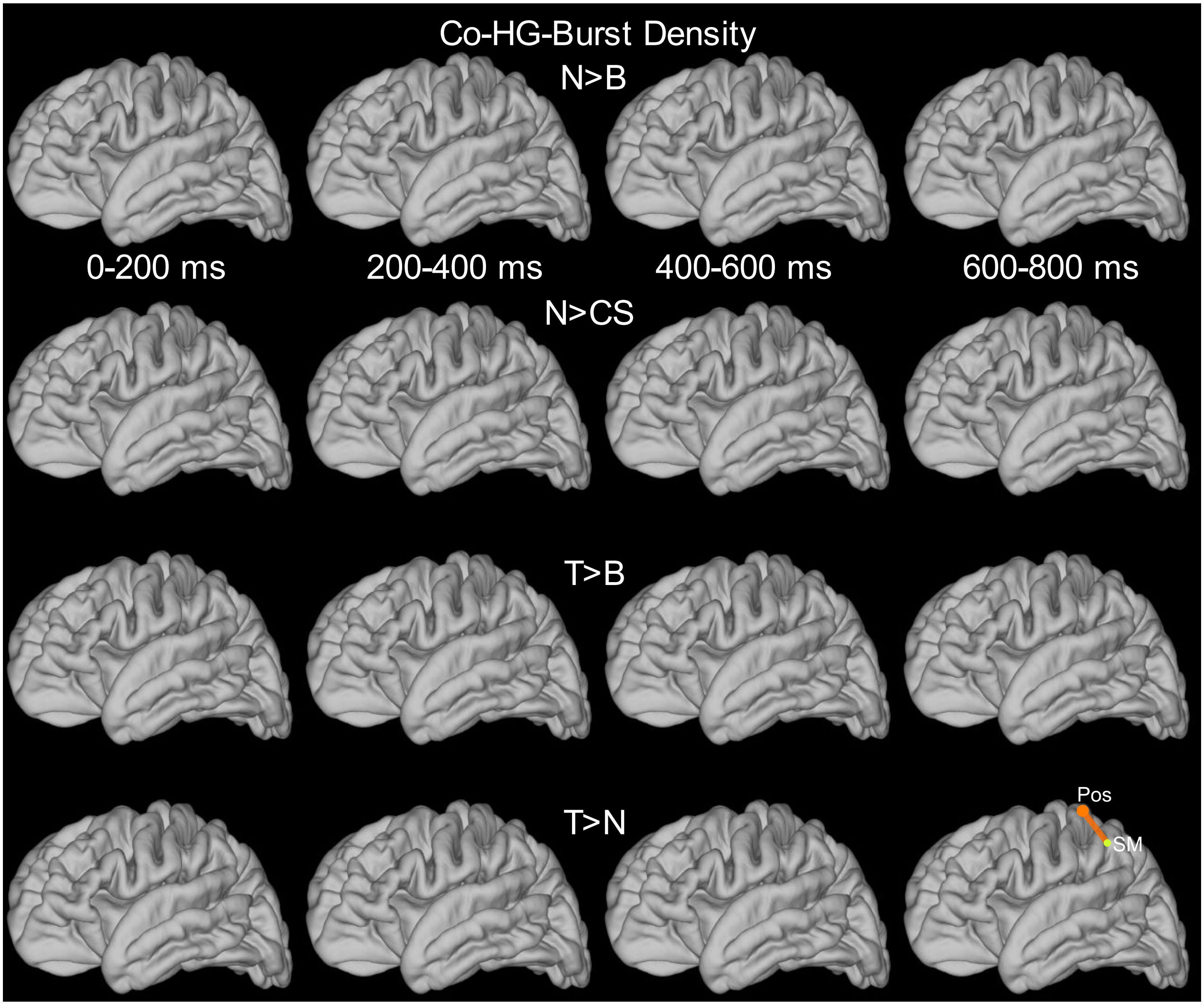


**Fig. S5. Co-high-gamma burst density is generally not modulated by stimulus condition.** Analysis and presentation identical to Fig. S3, except applied to Co-HG density rather than co-ripple density.

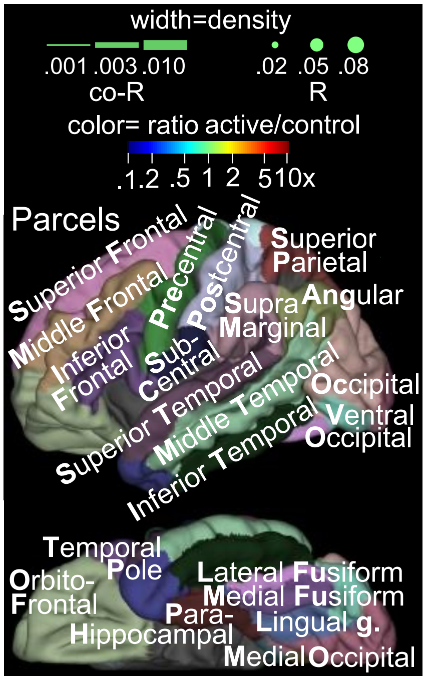

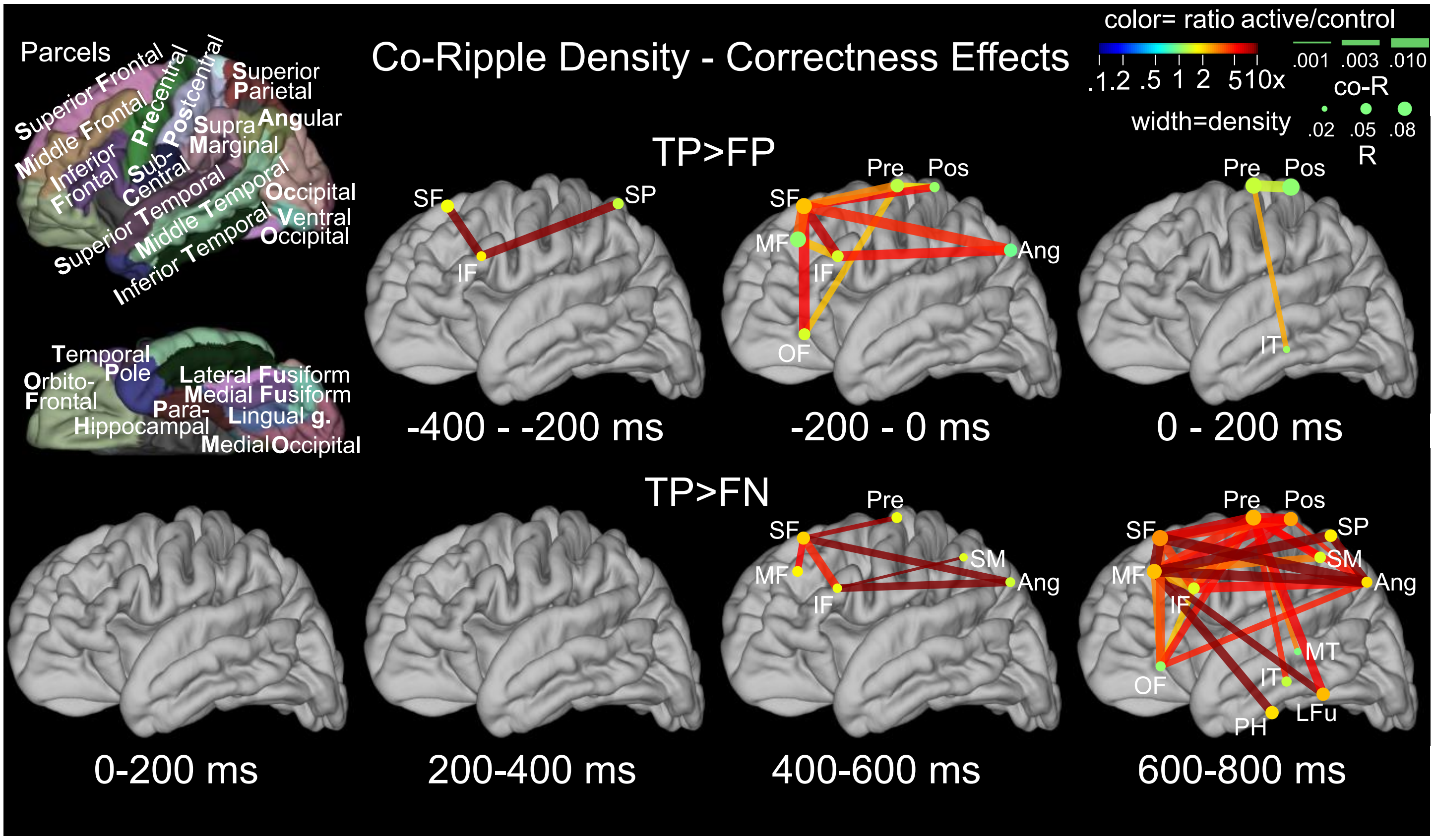


**Fig. S6. Selective executive-semantic co-ripple engagement during correct task performance.** Each row shows connections where the Co-R density was significantly higher in the indicated main condition relative to the control condition (stimulus-type permutation test, p<0.01 FDR corrected; procedures as described for Fig. 3). **TP>FP** (True Positive > False Positive) contrast for 200 ms latency windows, spanning 400 ms before to 200 ms after task responses. **TP > FN** (True Positive > False Negative) contrast from stimulus onset to 800 ms post-stimulus. Task responses typically occurred at approximately 800 ms post-stimulus. Correct task responses are specifically characterized by engagement of Co-R in an executive-semantic network prior to task responses. Incorrect responses did not show significantly greater Co-R than correct responses in any connection or latency. Density values represent the sum-total time occupied by Co-R over channel-pairs, trials, and subjects, relative to the theoretical maximum value. See **Materials and Methods, Co-ripples or co-high-gamma bursts** for more detail. Line and dot width represent Co-R and R density, respectively, in the main condition. **Note that the identical comparisons and analysis but conducted on co-high-gamma bursts rather than Co-R density revealed no significant effects** (data not shown).

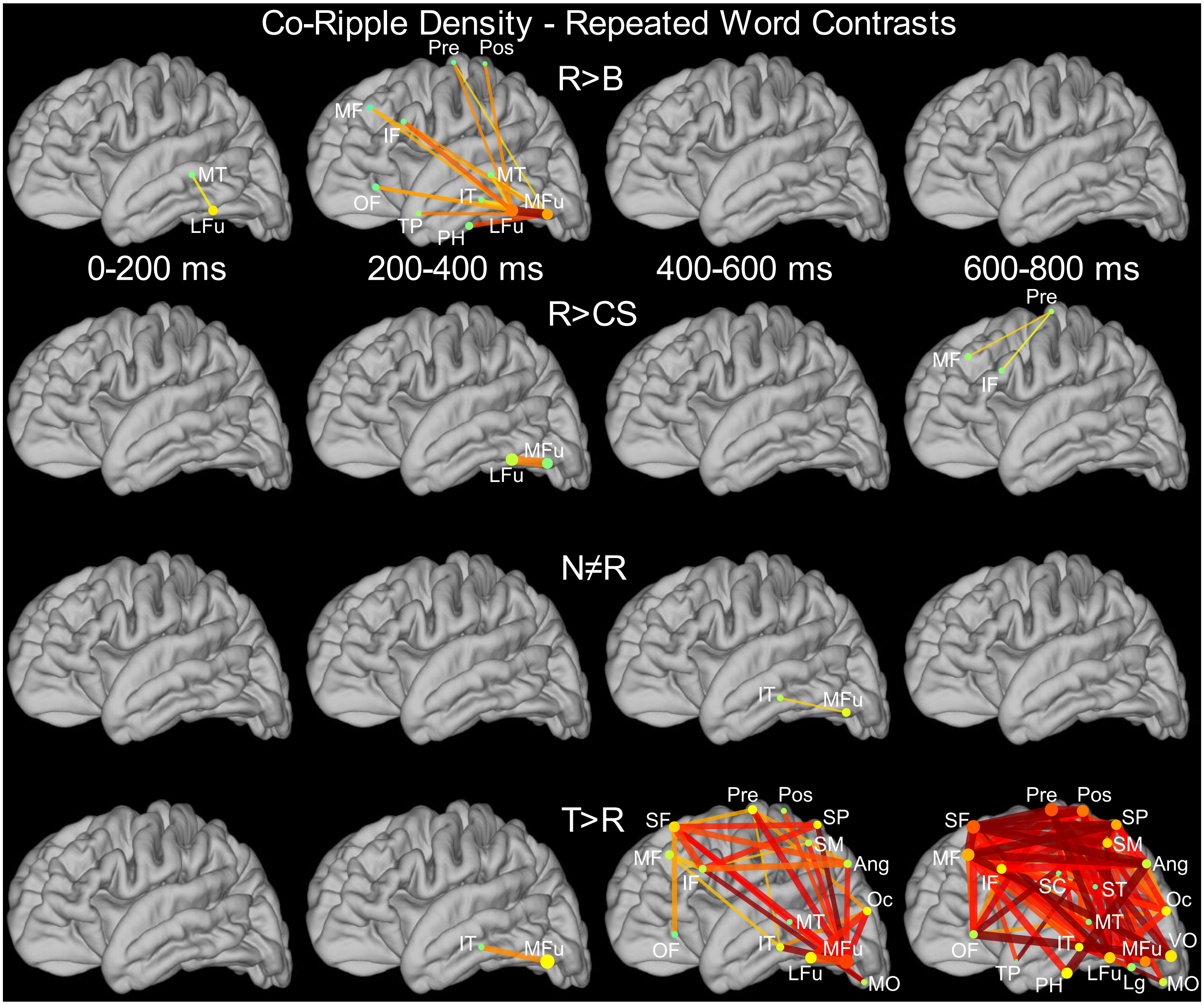


**Fig. S7. Repeated and Novel nontarget words evoke largely equivalent co-ripple density patterns.** Analysis and presentation of Rows 1, 2 and 5 of this figure are identical to those of Rows 1, 2, and 4 of Fig S3, except that comparisons are to Repeated nontarget words (R) rather than Novel nontarget words (N). Direct comparisons of N and R show only one weak difference, where N is greater than R. Connections are shown where the Co-R density was significantly higher in the indicated main condition relative to the control condition (stimulus-type permutation test, R>B, R>CS, T>R: $p<0.01$ FDR corrected; 1-tailed except N≠R which is 2-tailed). Density values represent the sum-total time occupied by Co-R over channel-pairs, trials, and subjects, relative to the theoretical maximum value.

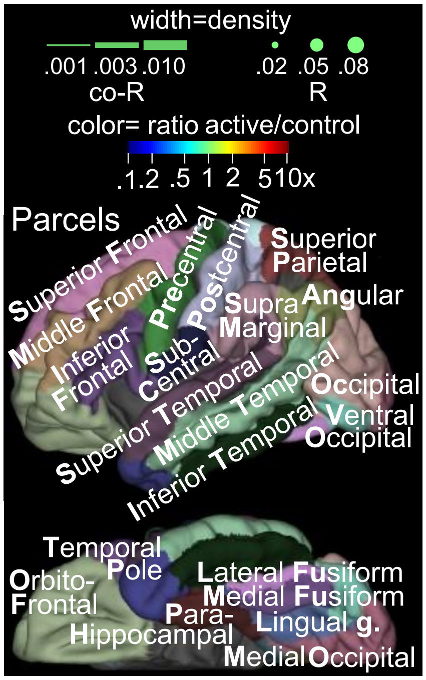

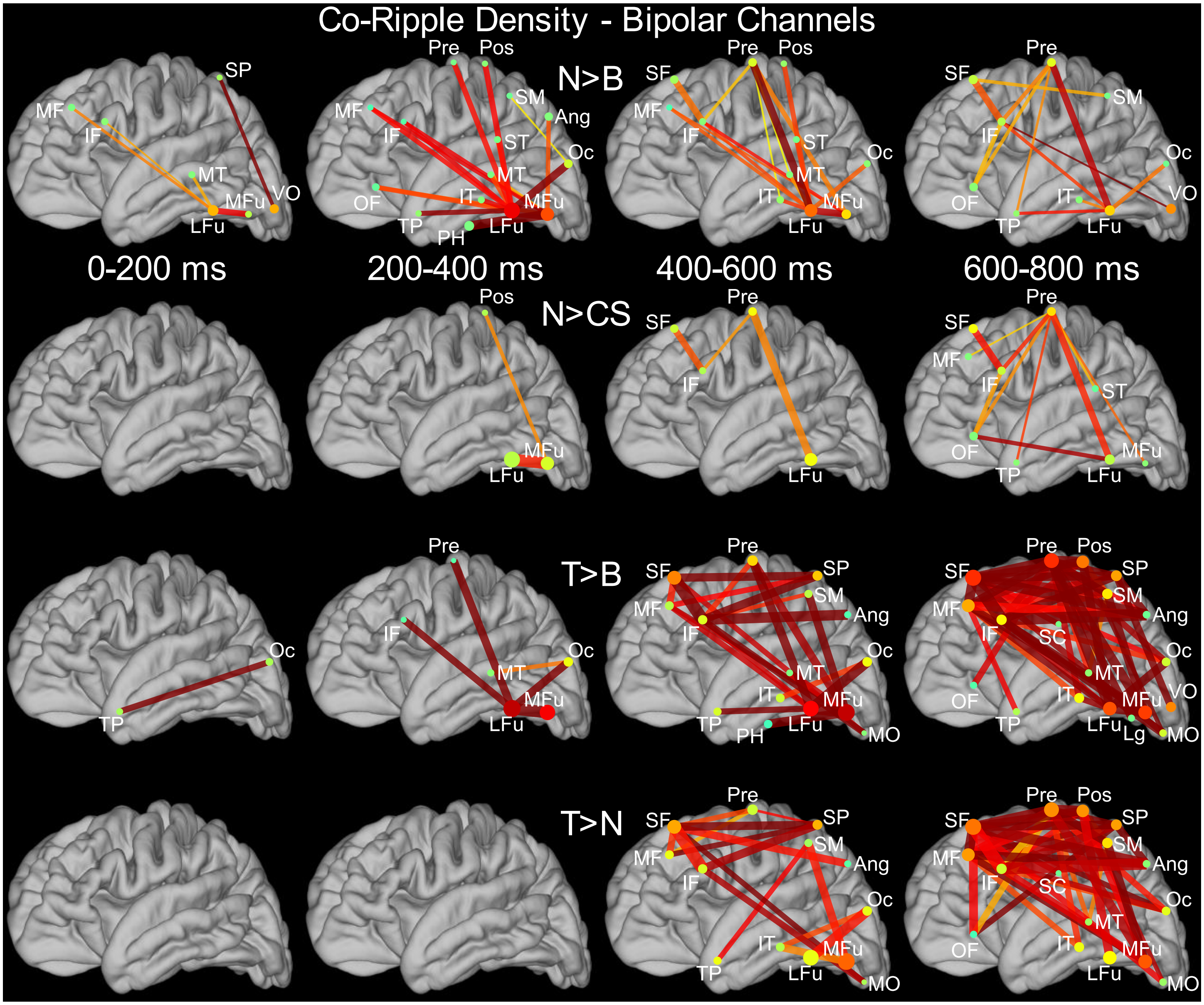


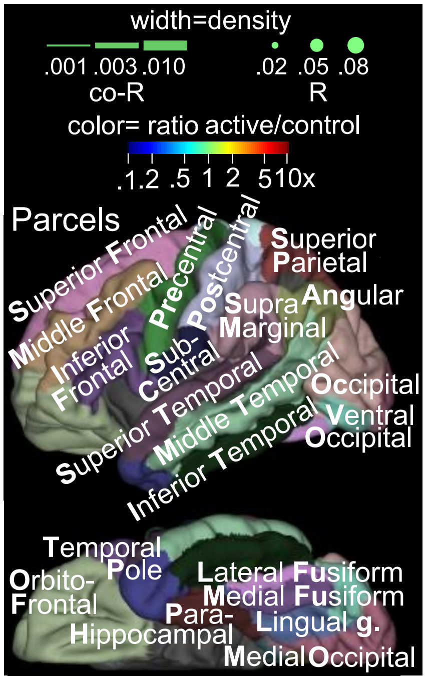


**Fig. S8. Major task effects are robust to bipolar re-referencing.** Re-referencing to bipolar channels was performed as described in **Materials and Methods, Bipolar re-referencing for co-ripple density analyses.** Analysis and presentation identical to Fig S3, except applied to co-ripples (Co-R) detected in bi-polar re-referenced recordings rather than the original referential recordings. Each row shows connections where the Co-R density was significantly higher in the indicated main condition relative to the control condition (stimulus-type permutation test, $p<0.01$ FDR corrected). Density values represent the sum-total time occupied by Co-R over channel-pairs, trials, and subjects, relative to the theoretical maximum value. See **Materials and Methods, Co-ripples or co-high-gamma bursts** for more detail. Line and dot width represent Co-R and R density, respectively, in the main condition.

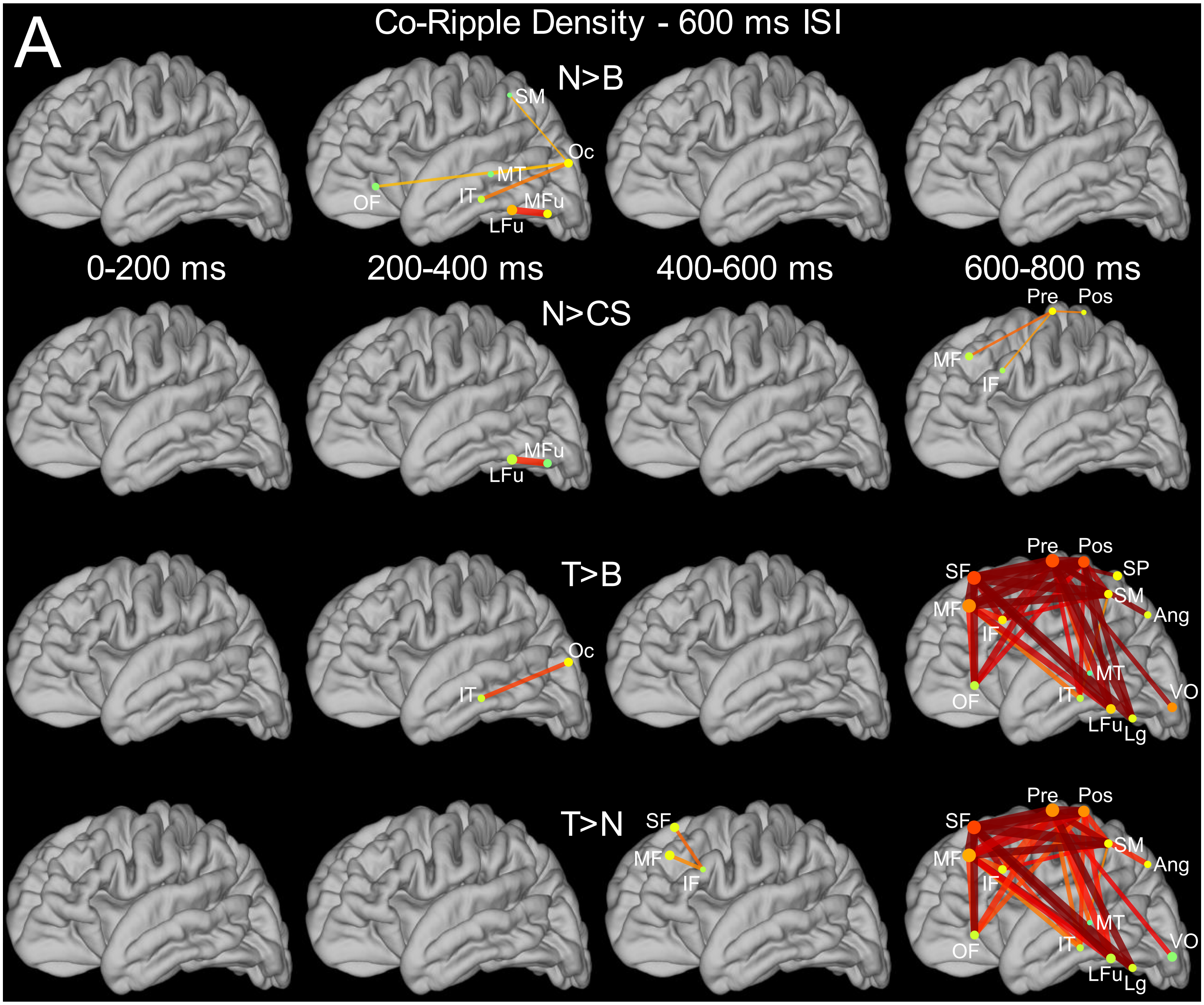


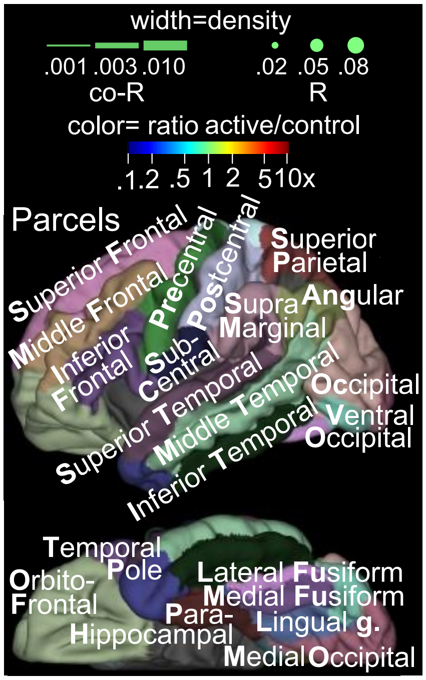


**Fig. S9. Spatiotemporal patterns of co-rippling are consistent between trials with 600 ms and 1000-1400 ms inter-stimulus intervals: A. 600ms duration trials in 7 subjects.** Analysis and presentation identical to Fig S3, except that only 600ms duration trials from Subjects P1-P6, and the relevant subset of trials from P8 are included. Each row shows connections where the Co-R density was significantly higher in the indicated main condition relative to the control condition (stimulus-type permutation test, $p<0.01$ FDR corrected). Density values represent the sum-total time occupied by Co-R over channel-pairs, trials, and subjects, relative to the theoretical maximum value. See **Materials and Methods, Co-ripples or co-high-gamma bursts for more detail**. Line and dot width represent Co-R and R density, respectively, in the main condition.

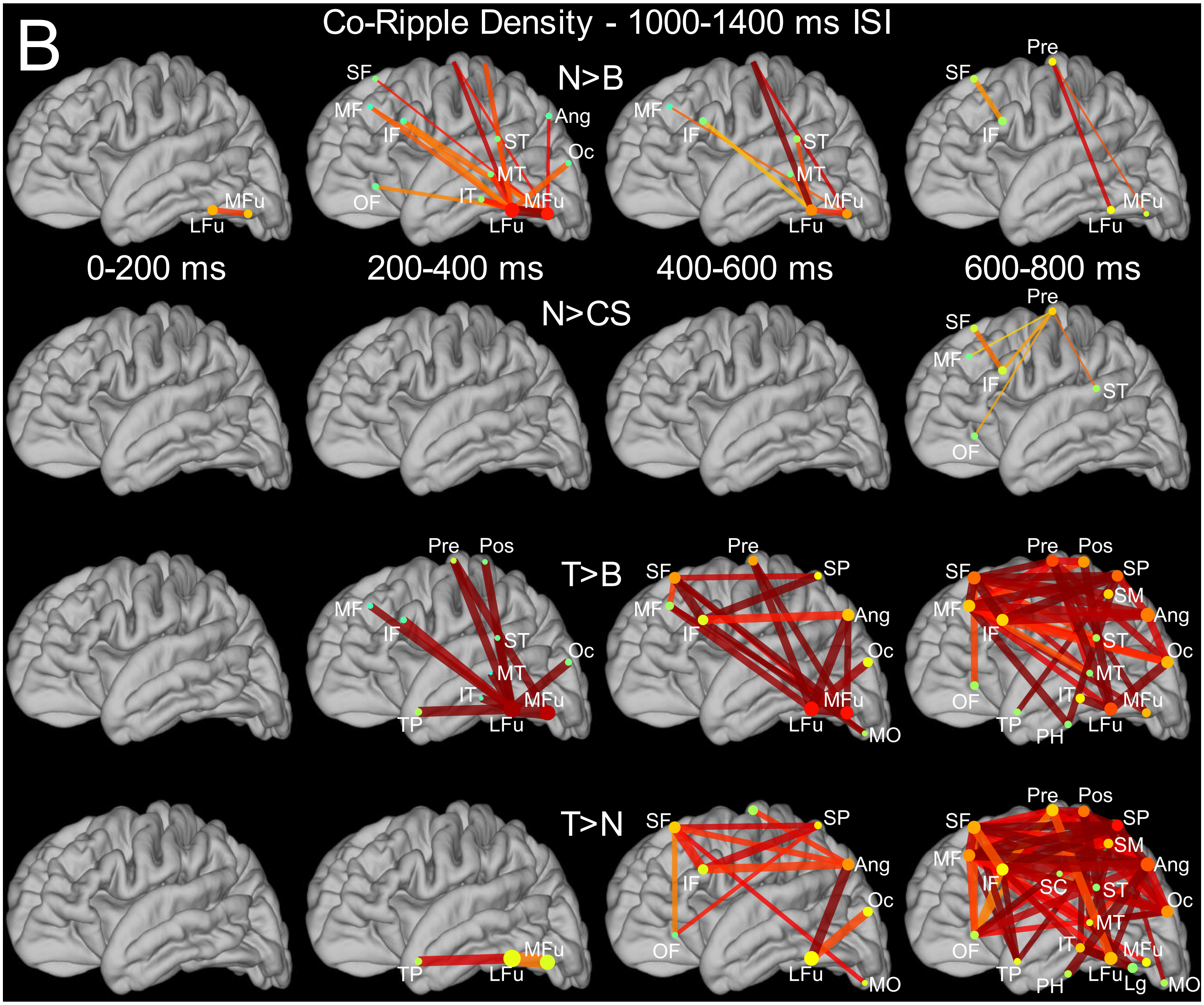


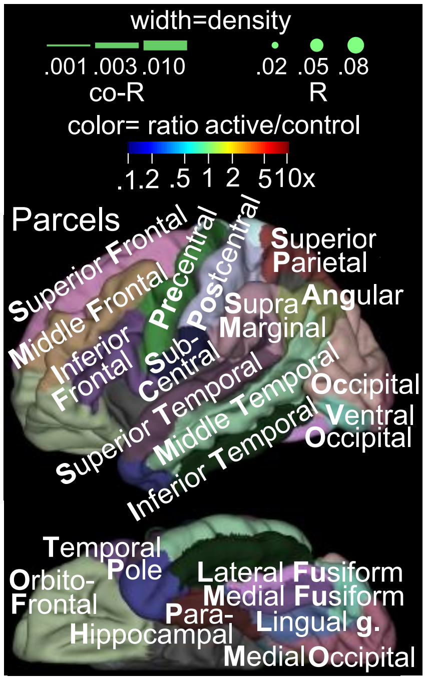


**Fig. S9. Spatiotemporal patterns of co-rippling are consistent between trials with 600 ms and 1000-1400 ms inter-stimulus intervals: B. 1000-1400ms duration trials in 4 subjects.** Analysis and presentation identical to Fig S3, except that only 1000-1400ms duration trials from Subjects P7, P9, P10, and the relevant subset of trials from P8 are included.

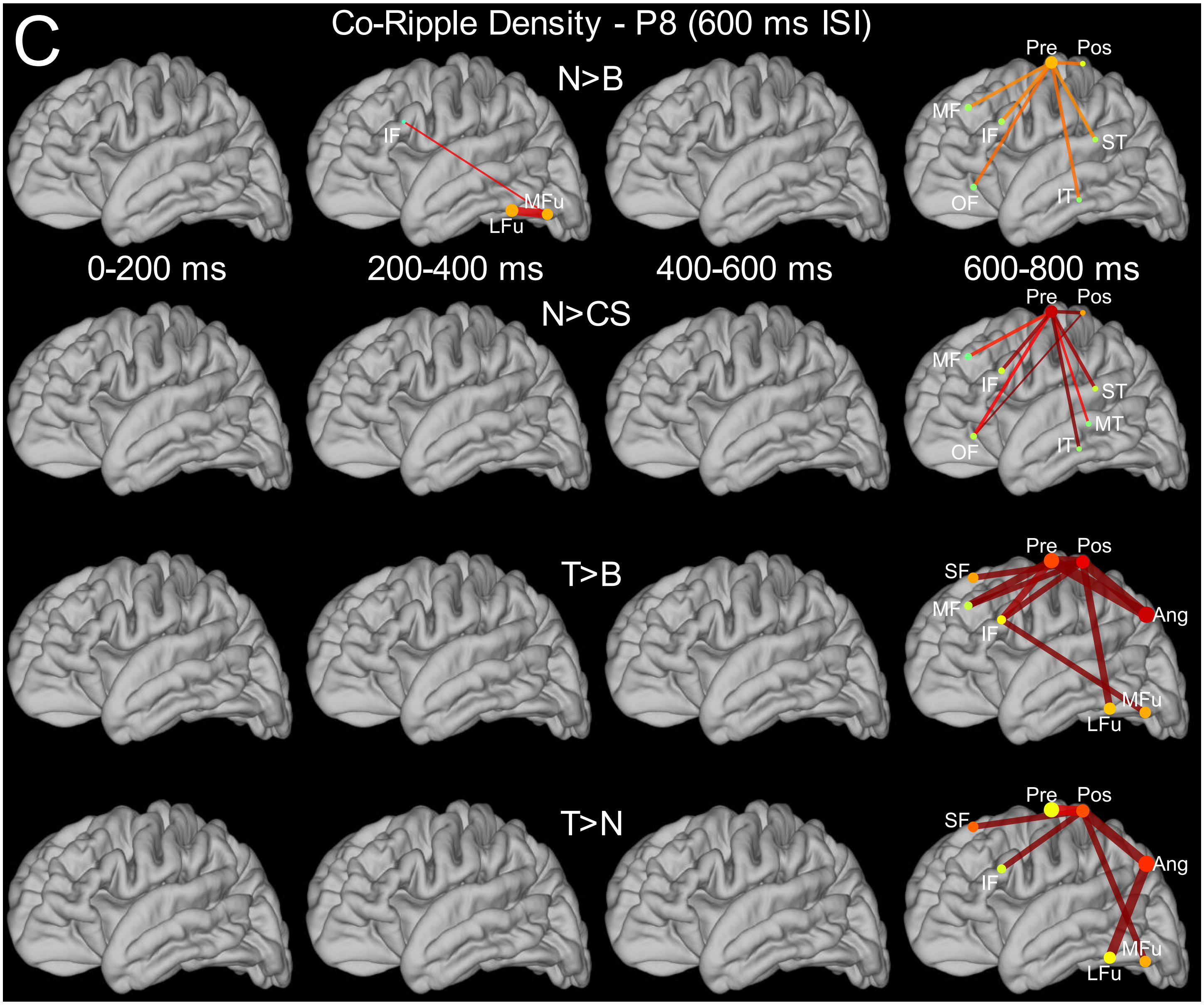


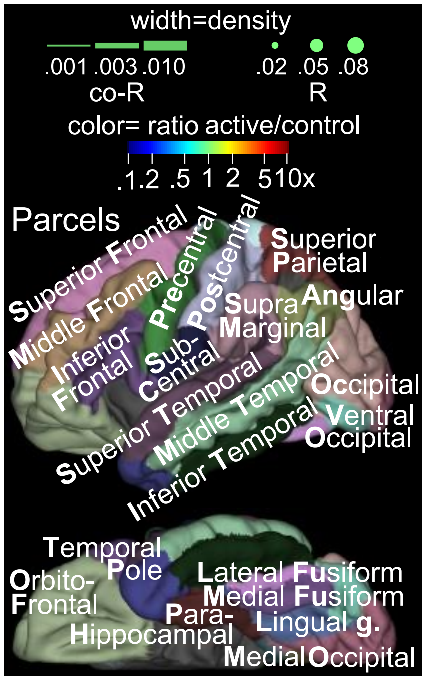


**Fig. S9. Spatiotemporal patterns of co-rippling are consistent between trials with 600 ms and 1000-1400 ms inter-stimulus intervals: C. 600ms duration trials in subject P8.** Analysis and presentation identical to Fig S3, except that only 600ms duration trials from Subject P8 are included.

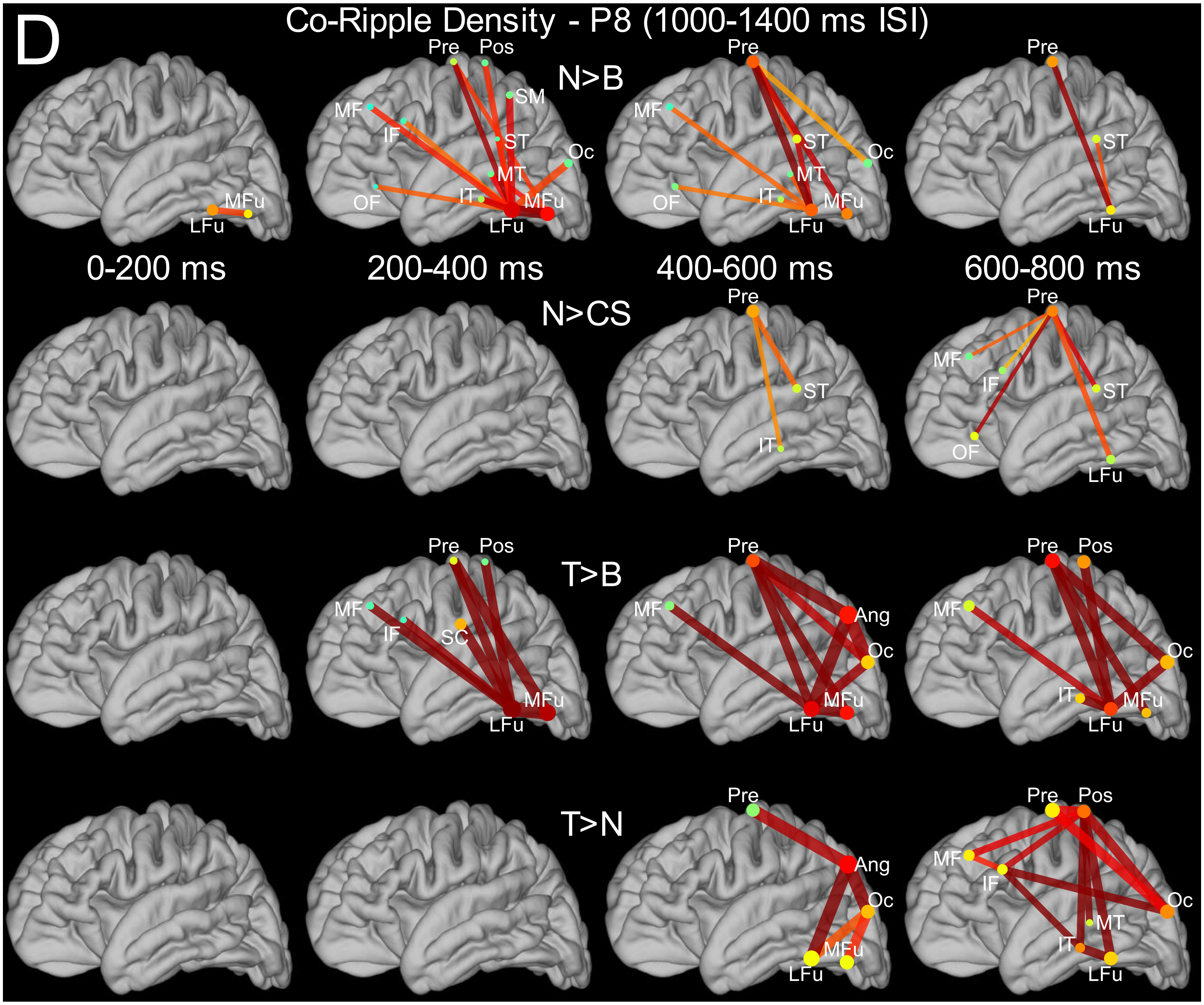


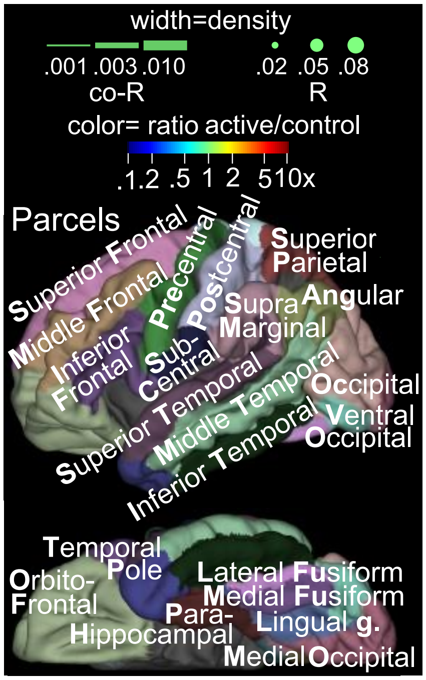


**Fig. S9. Spatiotemporal patterns of co-rippling are consistent between trials with 600 ms and 1000-1400 ms inter-stimulus intervals: D. 1000-1400ms duration trials in subject P8.** Analysis and presentation identical to Fig S3, except that only1000-1400ms duration trials from Subject P8 are included.

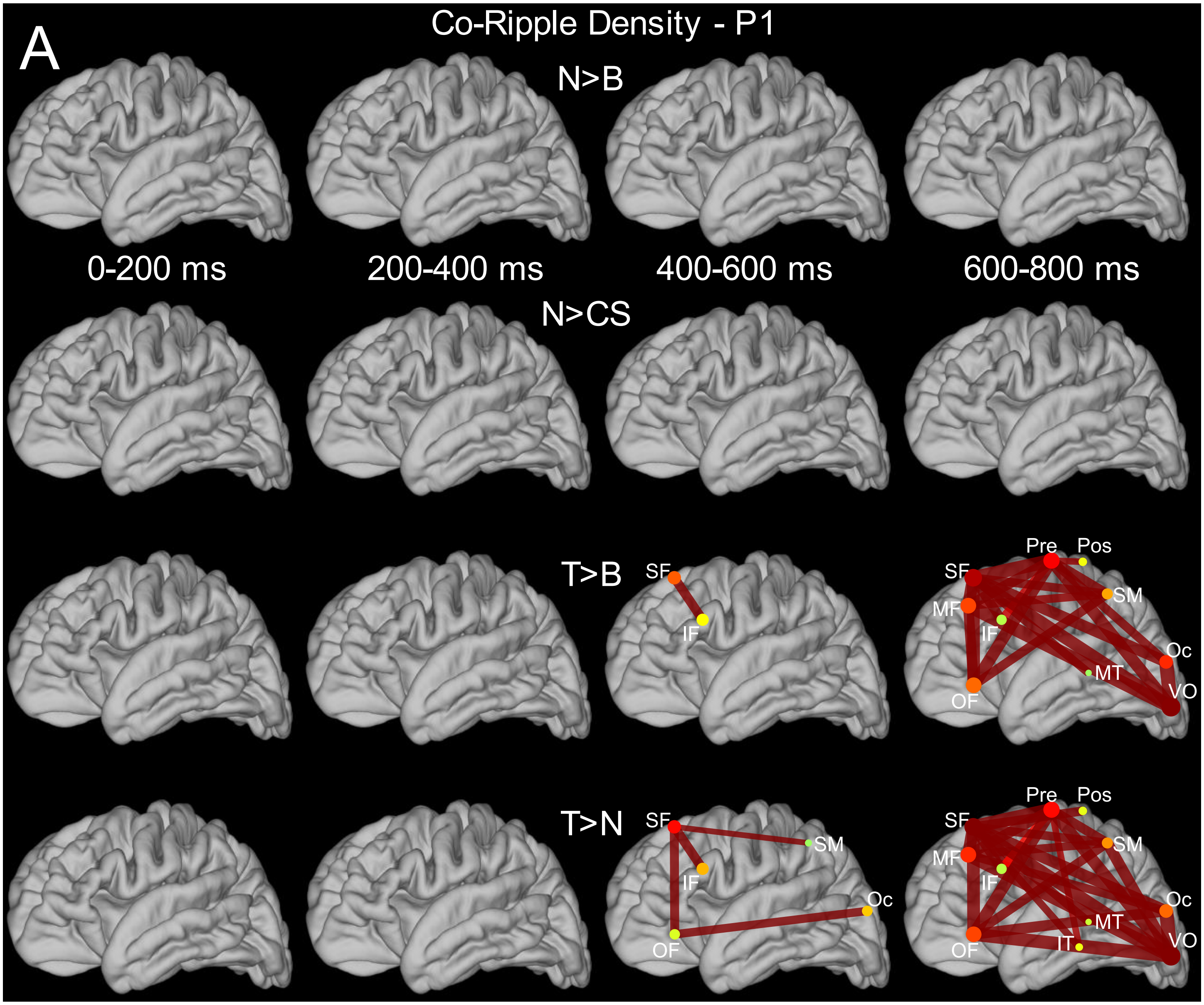


**Fig. S10. Major co-ripple density modulations resolve within individual subjects. A. Region-to-region evolution of Co-R density in subject P1.** Each row shows connections where the Co-R density was significantly higher in the indicated main condition relative to the control condition (stimulus-type permutation test, p<0.01 FDR corrected). Density values represent the sum-total time occupied by Co-R over channel-pairs, trials, and subjects, relative to the theoretical maximum value. Analysis and presentation identical to Fig S3, except that only trials from Subject P1 are included. See **Materials and Methods, Co-ripples or co-high-gamma bursts** for more detail. Line and dot width represent Co-R and R density, respectively, in the main condition.

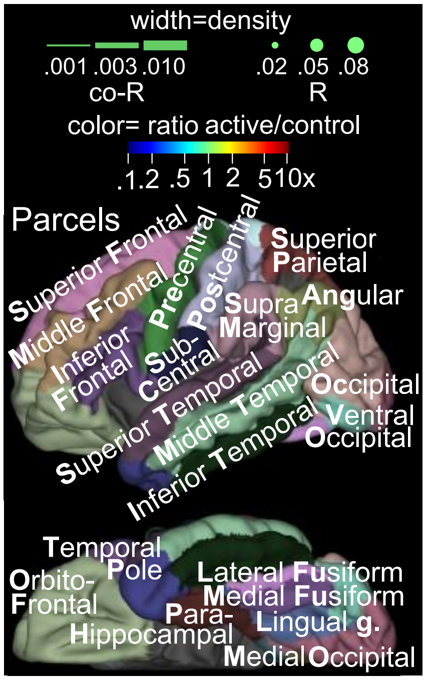

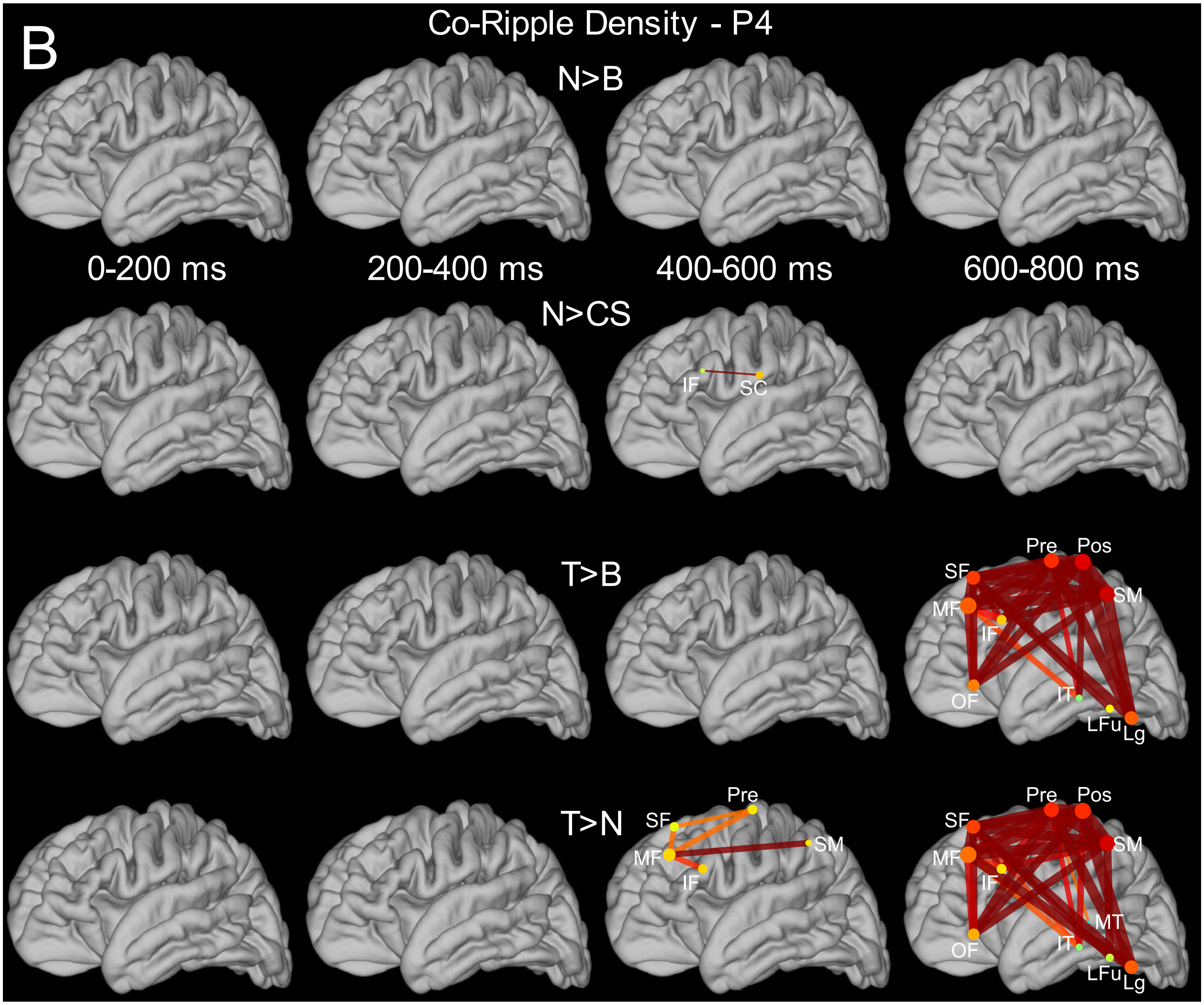


**Fig. S10B. Region-to-region evolution of co-ripple density in subject P4.** Analysis and presentation identical to Fig S3, except that only trials from Subject P4 are included.

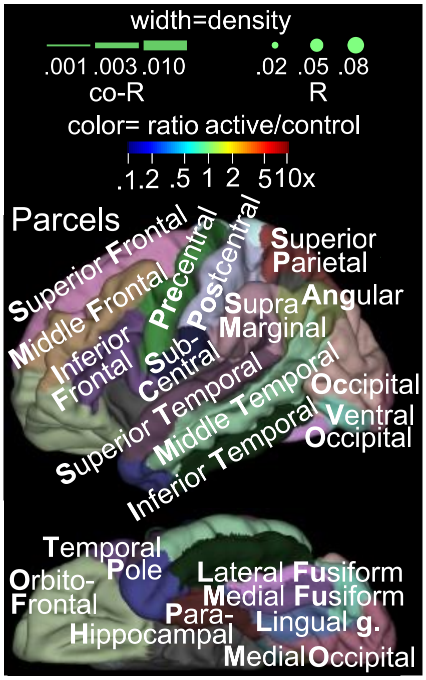

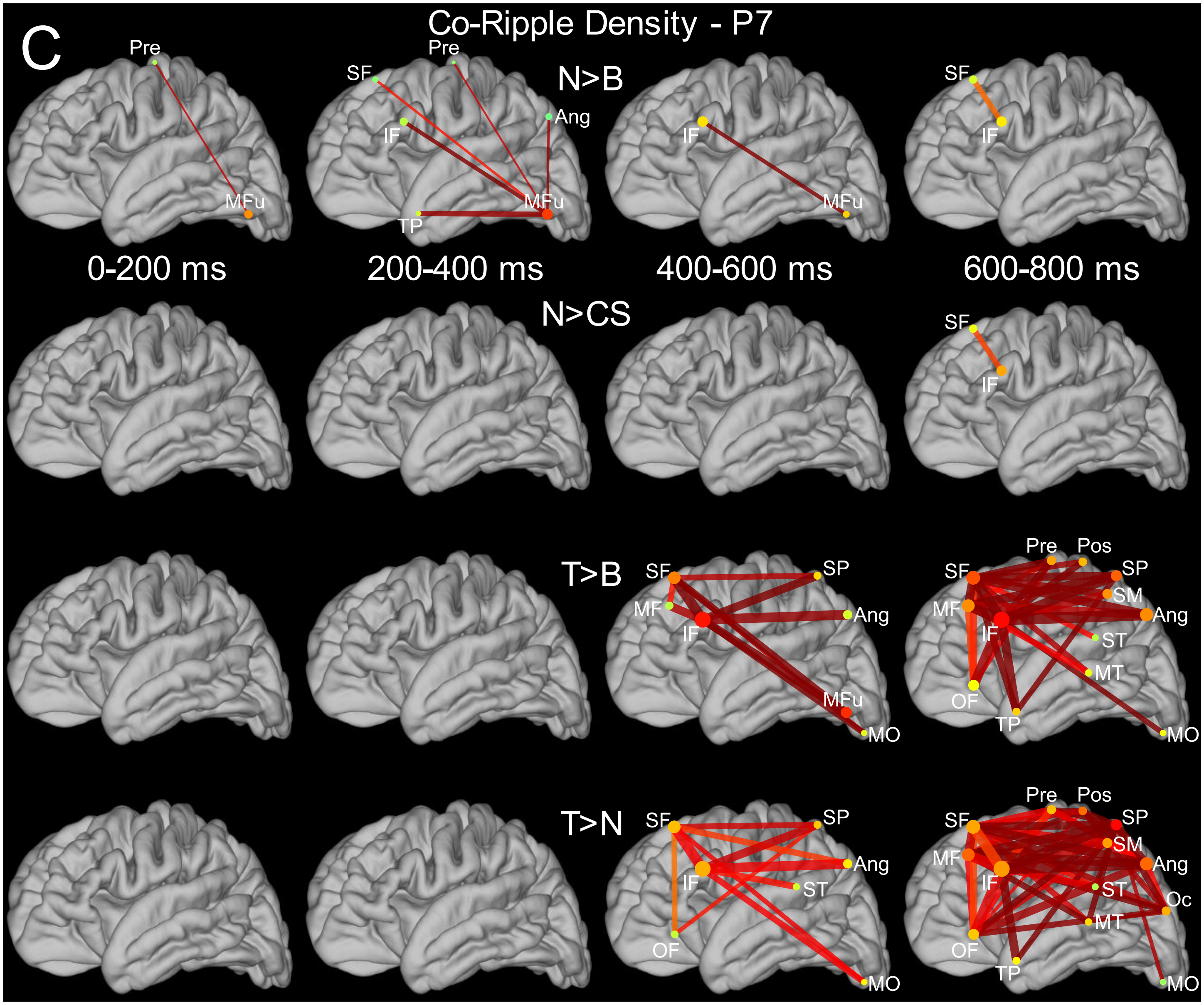


**Fig. S10C. Region-to-region evolution of co-ripple density in subject P7**. Analysis and presentation identical to Fig S3, except that only trials from Subject P7 are included.

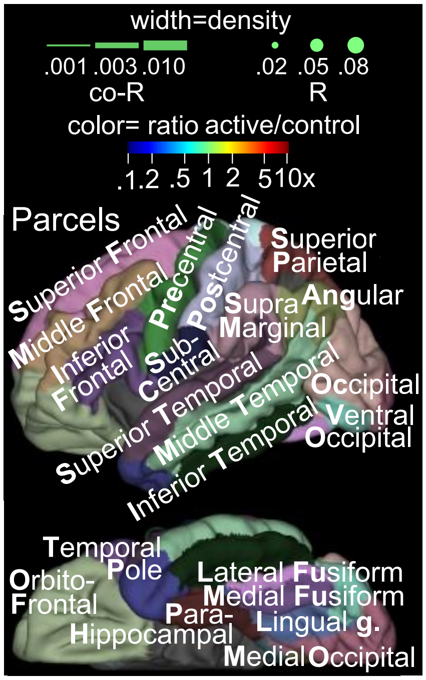

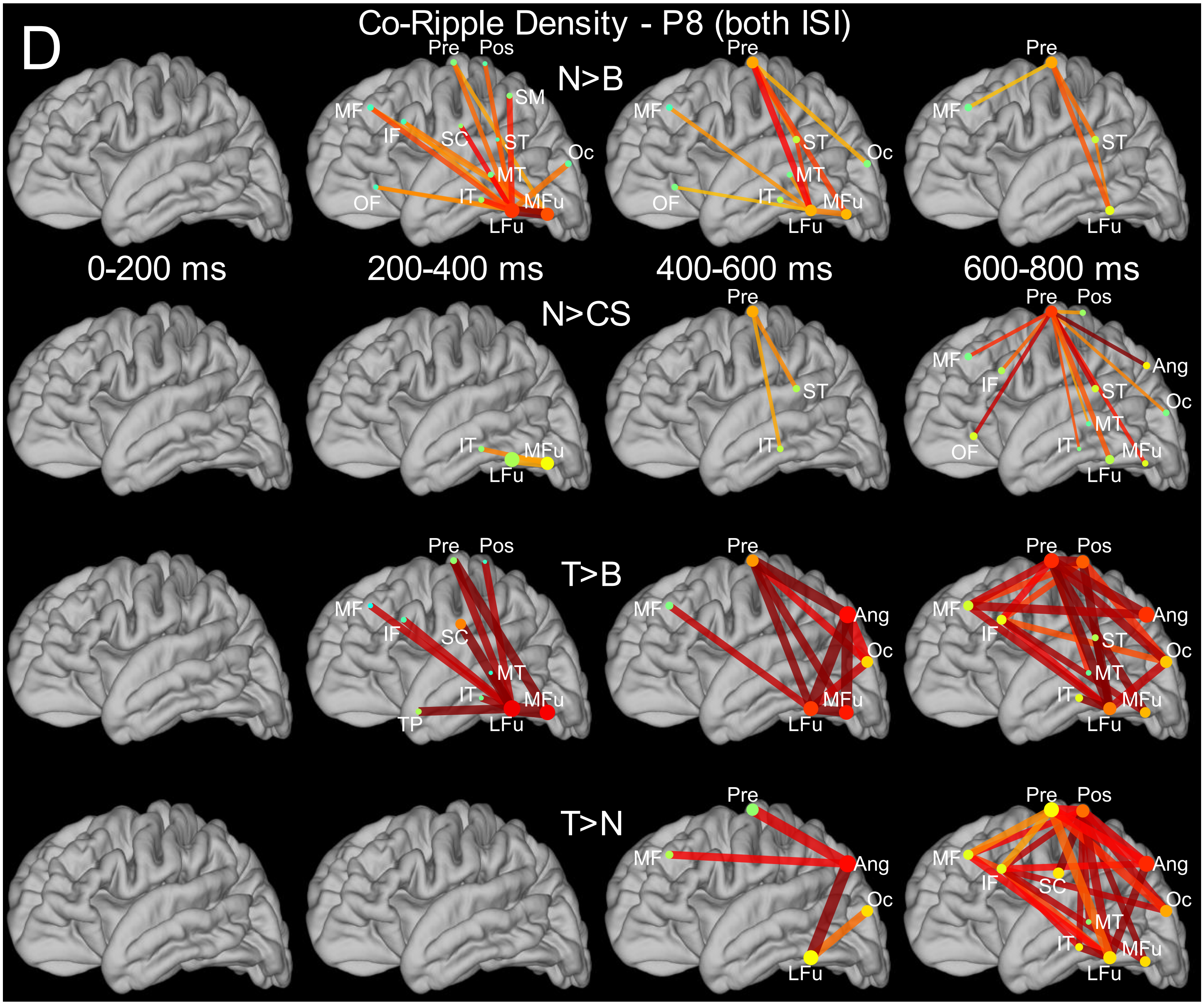


**Fig. S10D. Region-to-region evolution of co-ripple density in subject P8**. Analysis and presentation identical to Fig S3, except that only trials from Subject P8 are included. All trials (both 600 ms and 1000-1400ms ISI) are included (separate analyses of the different trial durations from P8 are shown in Fig. S9CD).

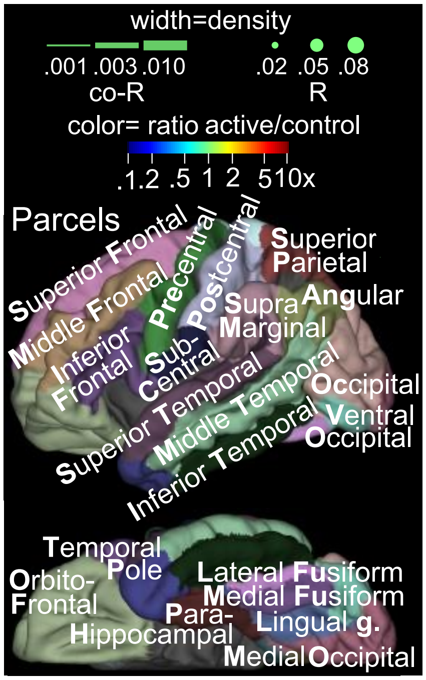

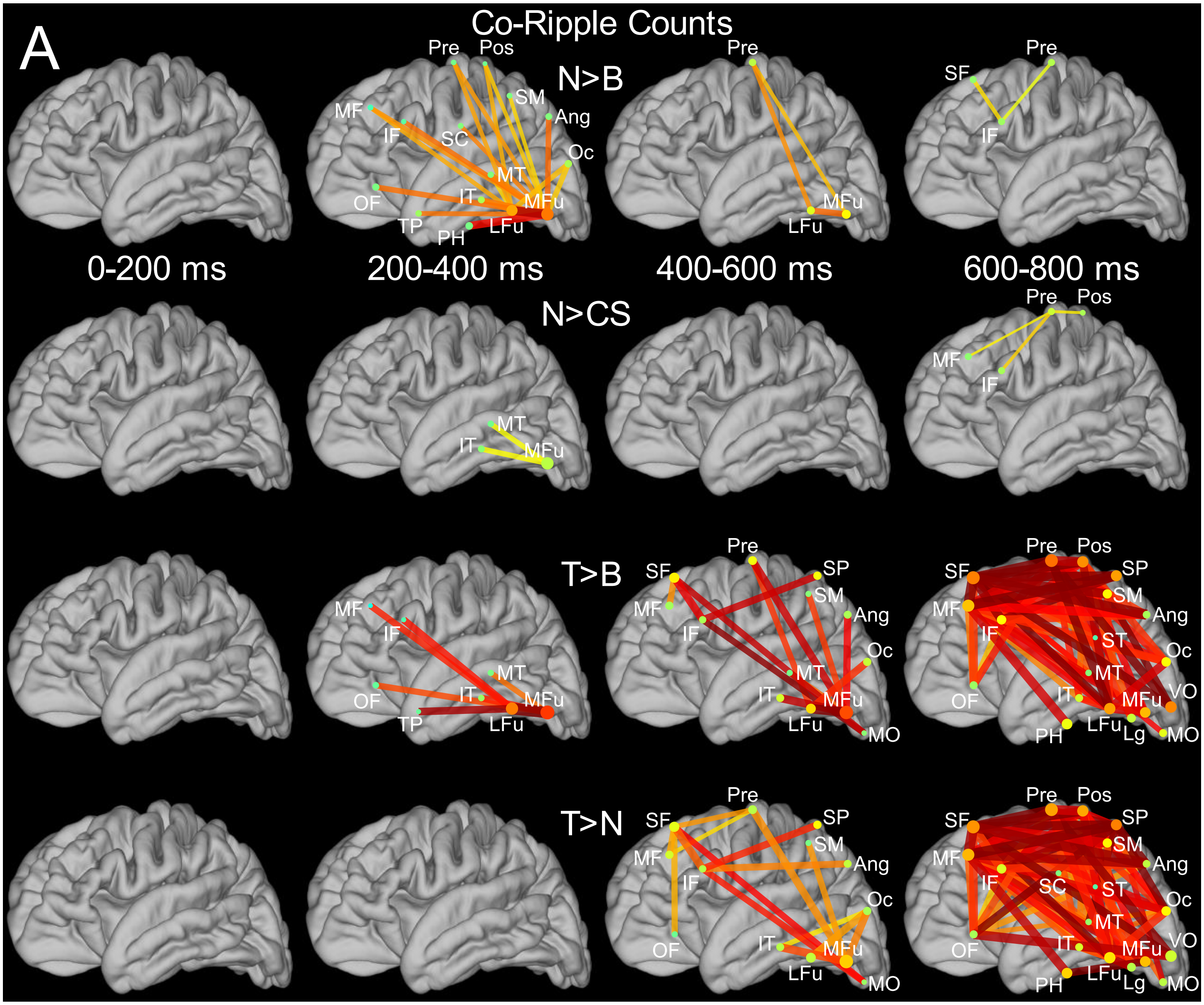


**Fig. S11. Quantification employing counts of co-ripple or co-high-gamma burst events yield similar results to density-based analysis. A. Co-R.** Analysis and presentation are identical to Fig S3, except that instead of quantifying the density of Co-R in bins defined by their latency, regions and task condition, here we quantify by counting the number of Co-R in the same bins. Cortico-cortical connecting lines indicate where the Co-R rate was significantly higher in the indicated main condition relative to the control condition (stimulus-type permutation test, p<0.01 FDR corrected). Line and dot width represent Co-R and R rate, respectively, in the main condition, reported in units of $s^{-1}\cdot$channel(-pair)$^{-1}$. Rate was calculated analogously to density values used elsewhere (see **Materials and Methods, Co-ripples or co-high-gamma bursts**), with the modification that counts of Co-R are summed, rather than Co-R durations, such that rate values shown here correspond to the average count of Co-R over all exemplar pairs and trials during the indicated latency. Results are similar to those obtained with the main analyses employing Co-R density.

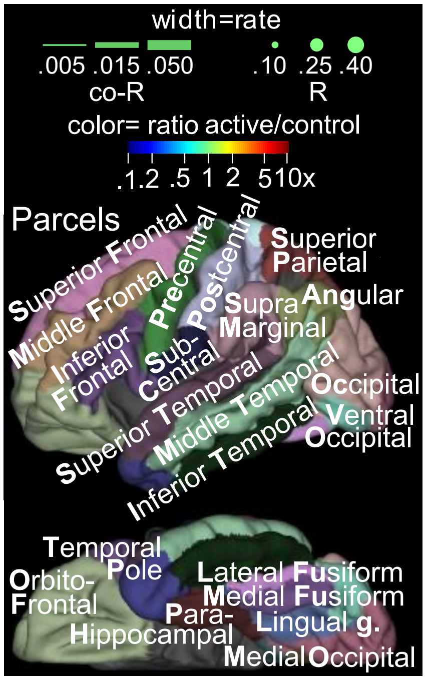

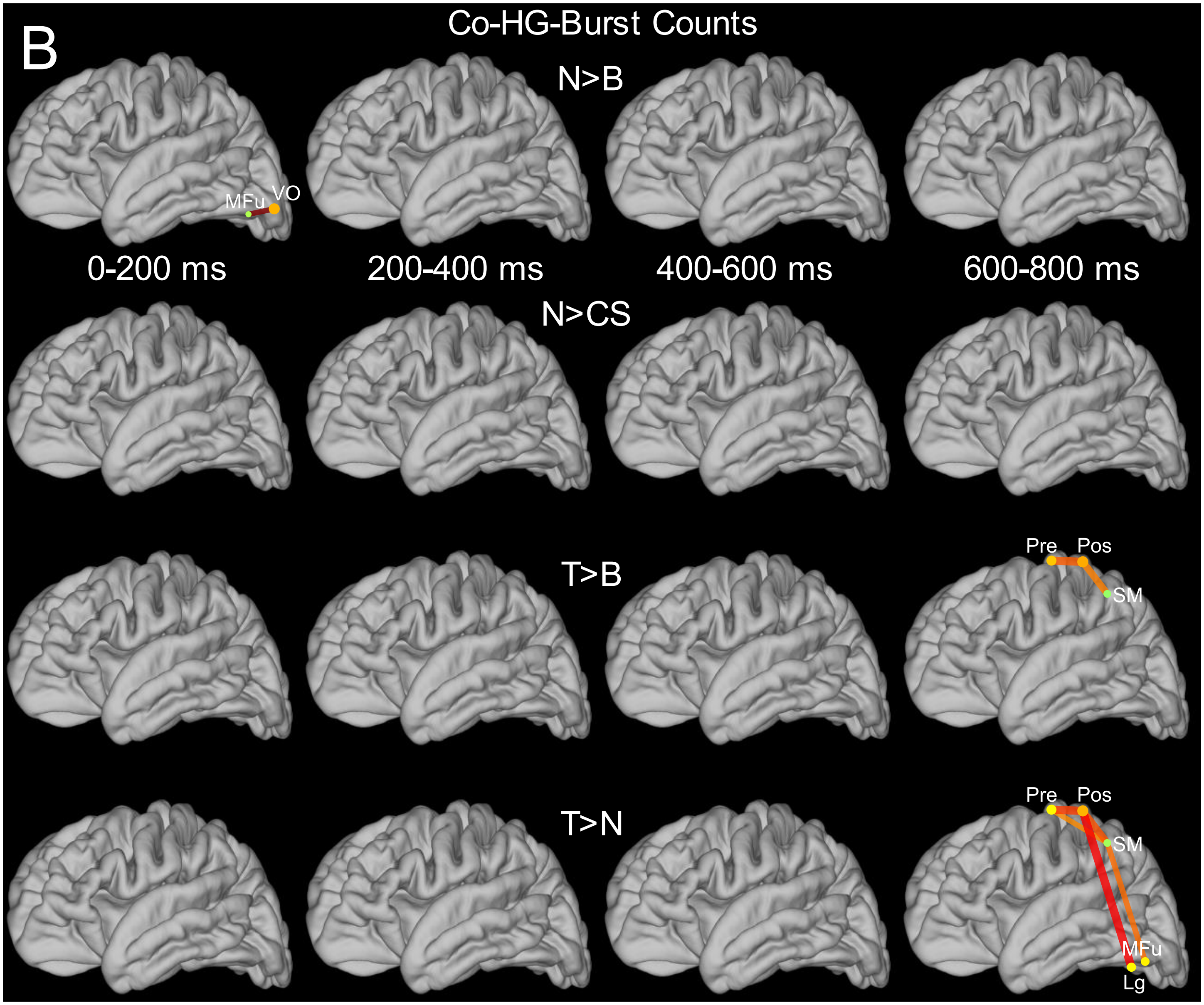


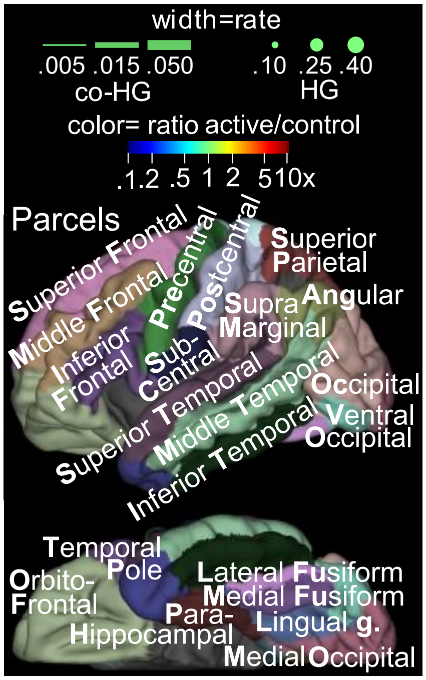


**Fig. S11. Analysis of task-modulation with counts instead of density. B. Co-high-gamma burst rate.** Each row shows connections where the Co-HG rate was significantly higher in the indicated main condition relative to the control condition (stimulus-type permutation test, p<0.01 FDR corrected). Line and dot width represent Co-HG and HGB rate, respectively, in the main condition, reported in units of $s^{-1}$·channel(-pair)$^{-1}$. As observed in analyses employing Co-HG density, task modulation is limited to early visual processing and to a small number of connections, mostly involving Rolandic cortex, approximately coinciding with task responses. Analysis and presentation are identical to Fig. S11A, except that Co-HG are quantified instead of Co-R.

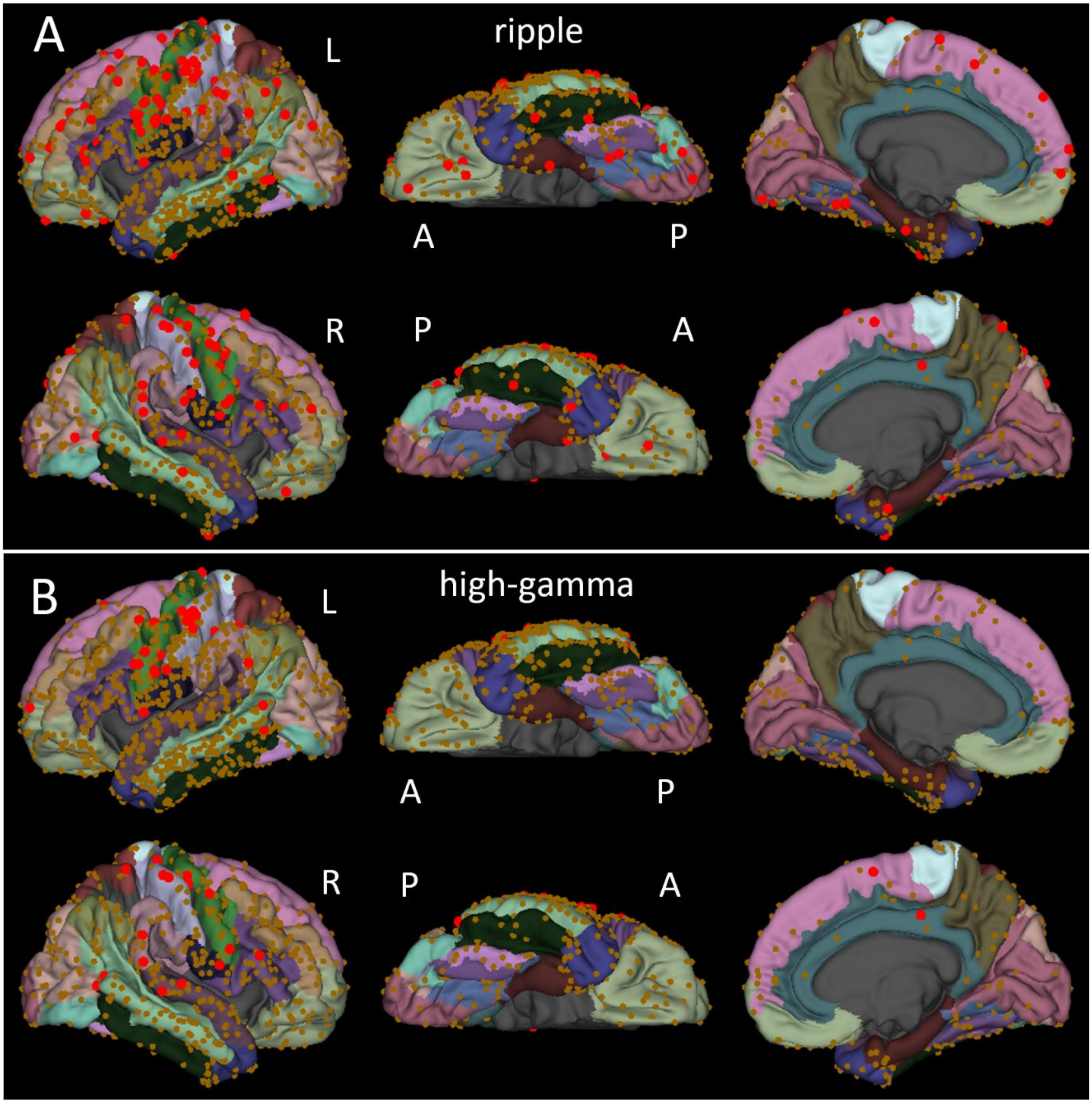


**Fig. S12. Ripple density and high gamma burst density show limited site-wise modulation by Target words. A.** Site-wise significant R density differences, Target Word (T)>Novel Word (N). Red dots indicate sites where R density was significantly different at some latency between active and control conditions (stimulus-type permutation test, one-tailed, FDR correction across sites and latencies, $p<0.01$). Brown dots indicate sites which were accepted for analysis but did not show a significant response. For each site, the density of R was calculated for each 200 ms period from 200 ms to 1000 ms post-stimulus. **B.** Site-wise significant HGB density (same analysis as A). See Fig. 3I for region labels.

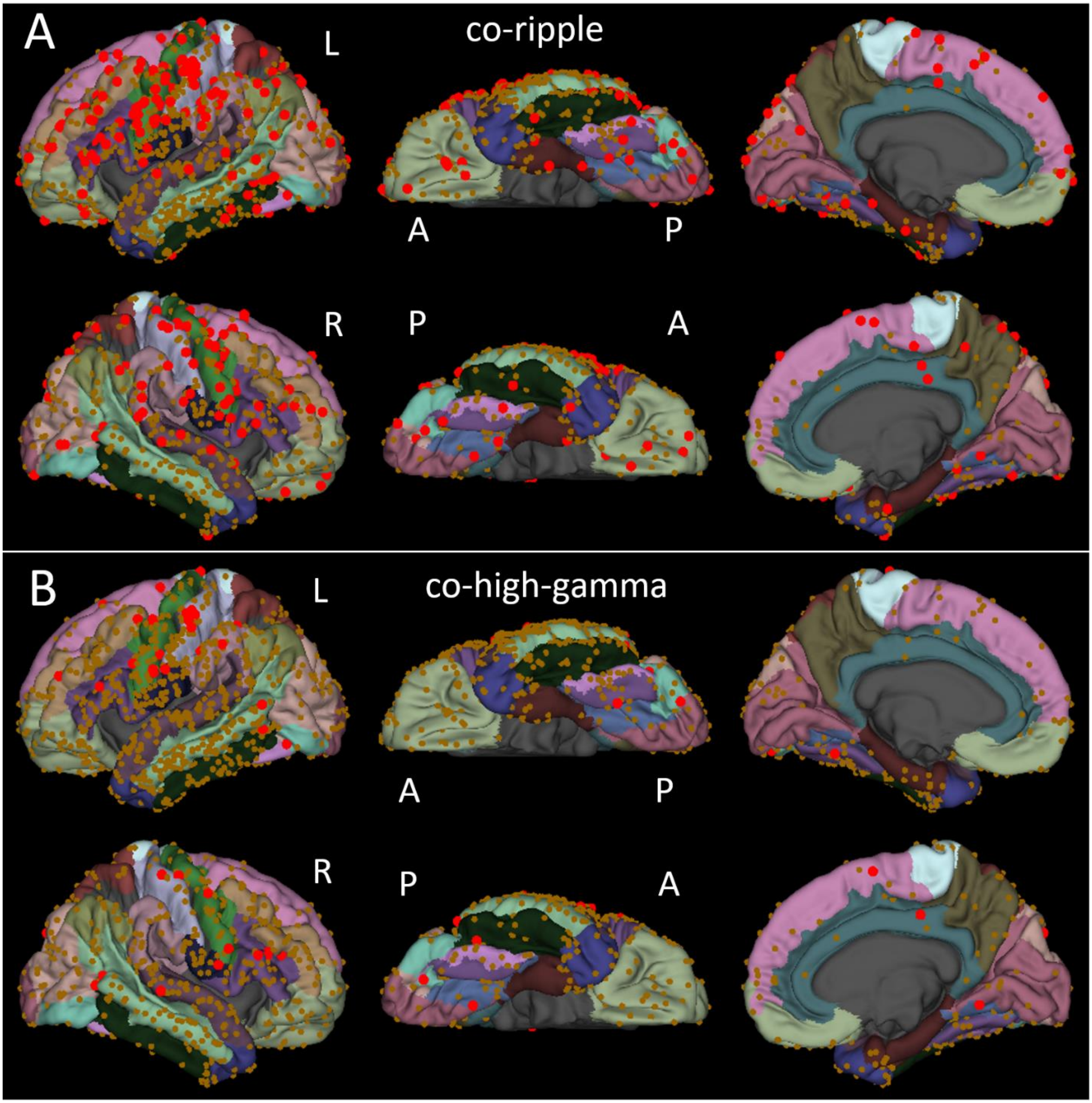


**S13. Site-wise co-ripple density modulation by Target words is more widespread than that of co-high-gamma bursts or ripples. A.** Site-wise significant Co-R density, Target Word (T)>Novel Word (N). Red dots indicate sites where Co-R density was significantly different at some latency between active and control conditions (stimulus-type permutation test, one-tailed, FDR correction across sites and latencies, p<0.01). Brown dots indicate sites which were accepted for analysis but did not show a significant response. For each site, the density of Co-R with any other site was calculated for each 200 ms period from 200 ms to 1000 ms post-stimulus. **B.** Site-wise significant Co-HG density, T>N. See Fig. 3I for region labels.

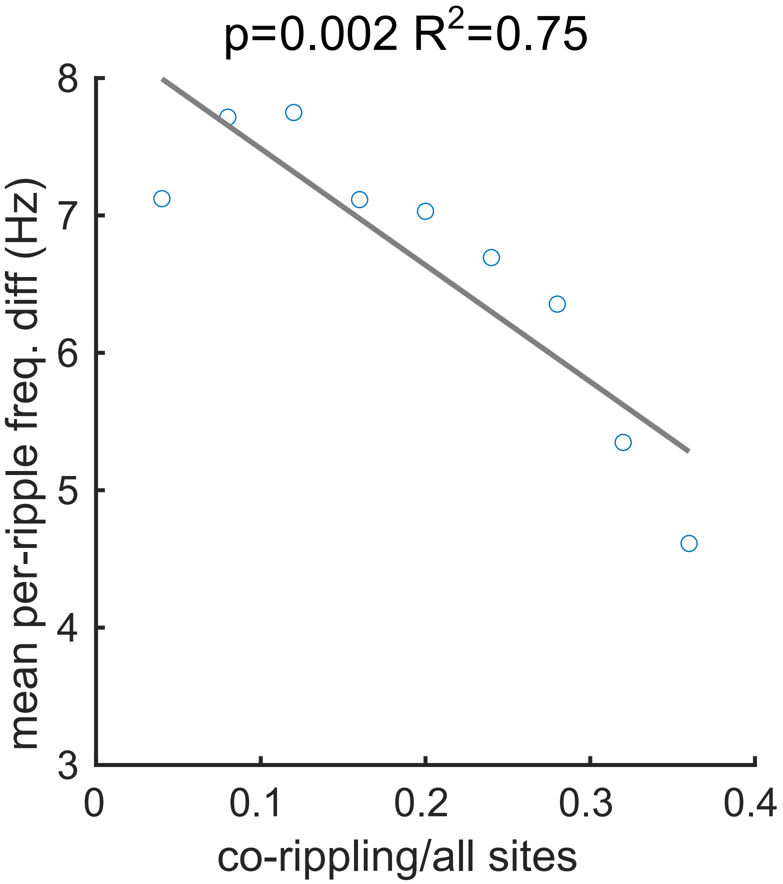


**Fig. S14. Widespread co-rippling is associated with more consistent ripple frequency across sites.**
The average difference between ripple frequencies in each pair of co-rippling sites is plotted against the proportion of the recorded sites which are co-rippling with them. As more sites co-ripple simultaneously, ripples exhibit more consistent frequencies (ordinary least squares regression; -8.5 Hz [-12.5, -4.4], two-tailed t-test, p=0.002, t=-5.0, df=7).

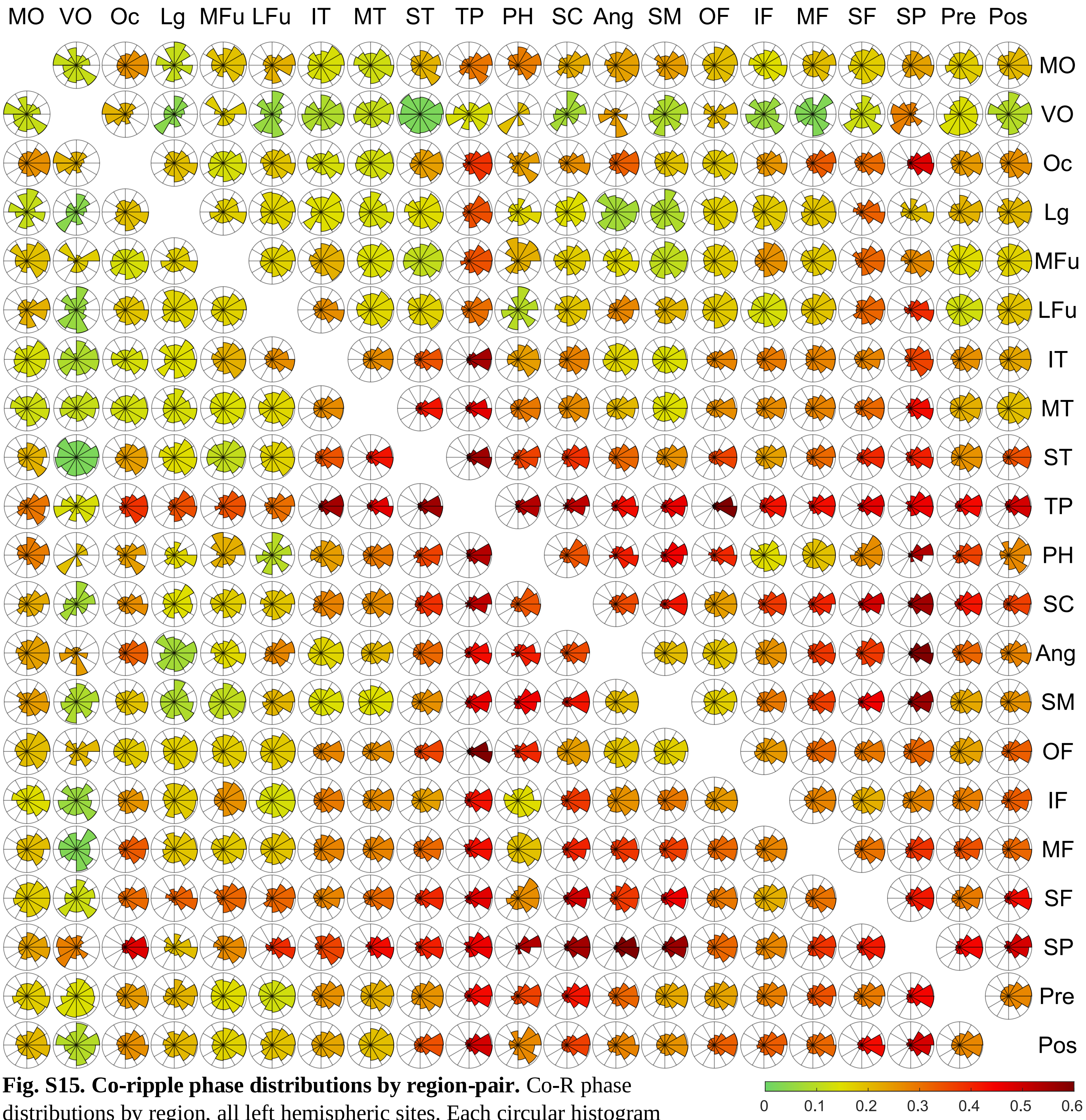


**Fig. S15. Co-ripple phase distributions by region-pair.** Co-R phase distributions by region, all left hemispheric sites. Each circular histogram represents counts of Co-R in each phase bin between the regions noted to the top and right. The range of 0 - 2 pi is divided into 12 equal bins, incrementing counterclockwise. Regions are ordered *a priori* along a visual-semantic-executive-response axis. Color of each histogram indicates the Phase-Locking Value (PLV) of the phase-distribution. The number of Co-R in each rosette, PLV value and significance, and zero-tendency, for each region-pair are shown in Tables S8 and S9. Note that the retinotopic visual areas (MO, VO) and to a lesser degree other ventral stream areas have lower PLV values than other cortical regions. (**MO**: medial occipital (early visual) cortex, **VO**: Ventral occipital cortex, **Oc**: dorsal and lateral occipital cortex, comprising the inferior occipital gyrus, **Lg**: Lingual gyrus excluding V1/V2, **mFu**: medial fusiform cortex including HCP-defined regions V8 and VVC, **lFu**: lateral fusiform cortex comprising the fusiform face complex, **IT**: inferior temporal gyrus, **MT**: middle temporal gyrus, **ST**: superior temporal gyrus, **TP**: temporal pole, **PH**: parahippocampal gyrus, **SC**: subcentral gyrus, **Ang**: angular gyrus**, SM**: supramarginal gyrus, **OF**: orbitofrontal cortex, **IF**: inferior frontal gyrus, **MF**: middle frontal gyrus, **SF**: superior frontal gyrus, **SP**: superior parietal lobule, **Pre**: precentral gyrus, **Pos**: postcentral gyrus)

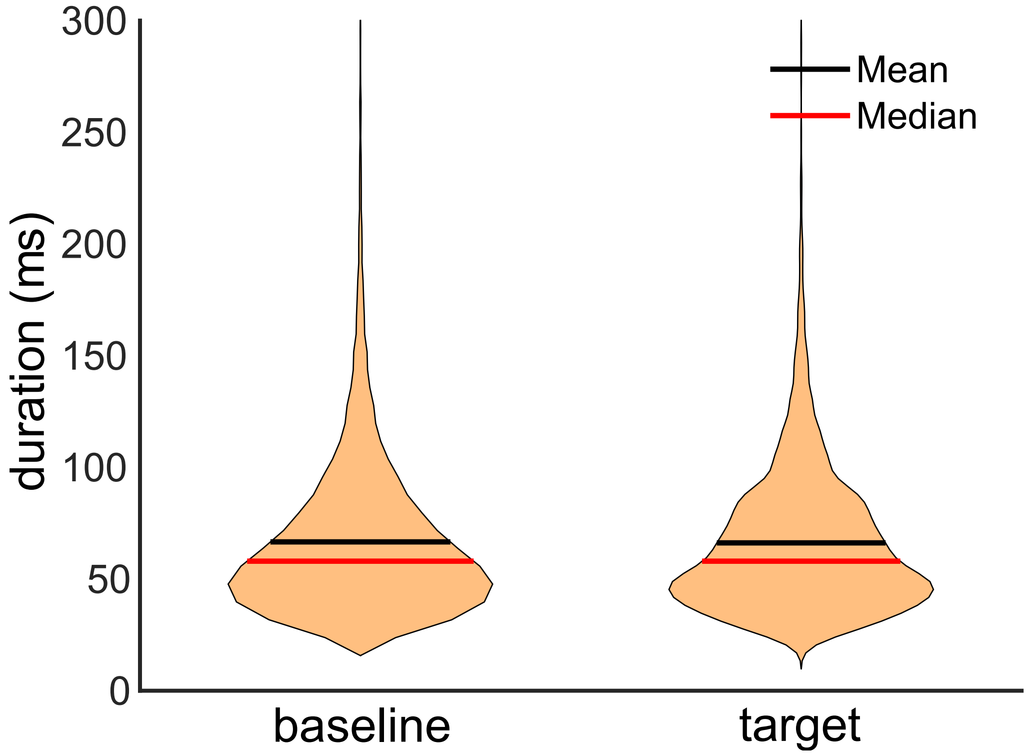


**Fig. S16. Target-evoked co-ripple durations do not differ from Baseline.** Violin plots represent the distributions of Co-R duration, at 100 ms pre-stimulus Baseline (left), and 700-800 ms post-stimulus on Target trials, the period of maximal Co-R density. Mean and median are represented by black and red horizontal lines, respectively. Target-evoked increases in Co-R density were therefore driven entirely by increase in the occurrence of Co-R, not by an increase in Co-R duration. Number of co-ripples: Baseline, Target = 49,220, 5,483; mean = 66.6, 66.2; median = 58, 58. Note that this is the overlap time of ripples in two locations, and thus is shorter than the duration of single ripples (Fig. S1).

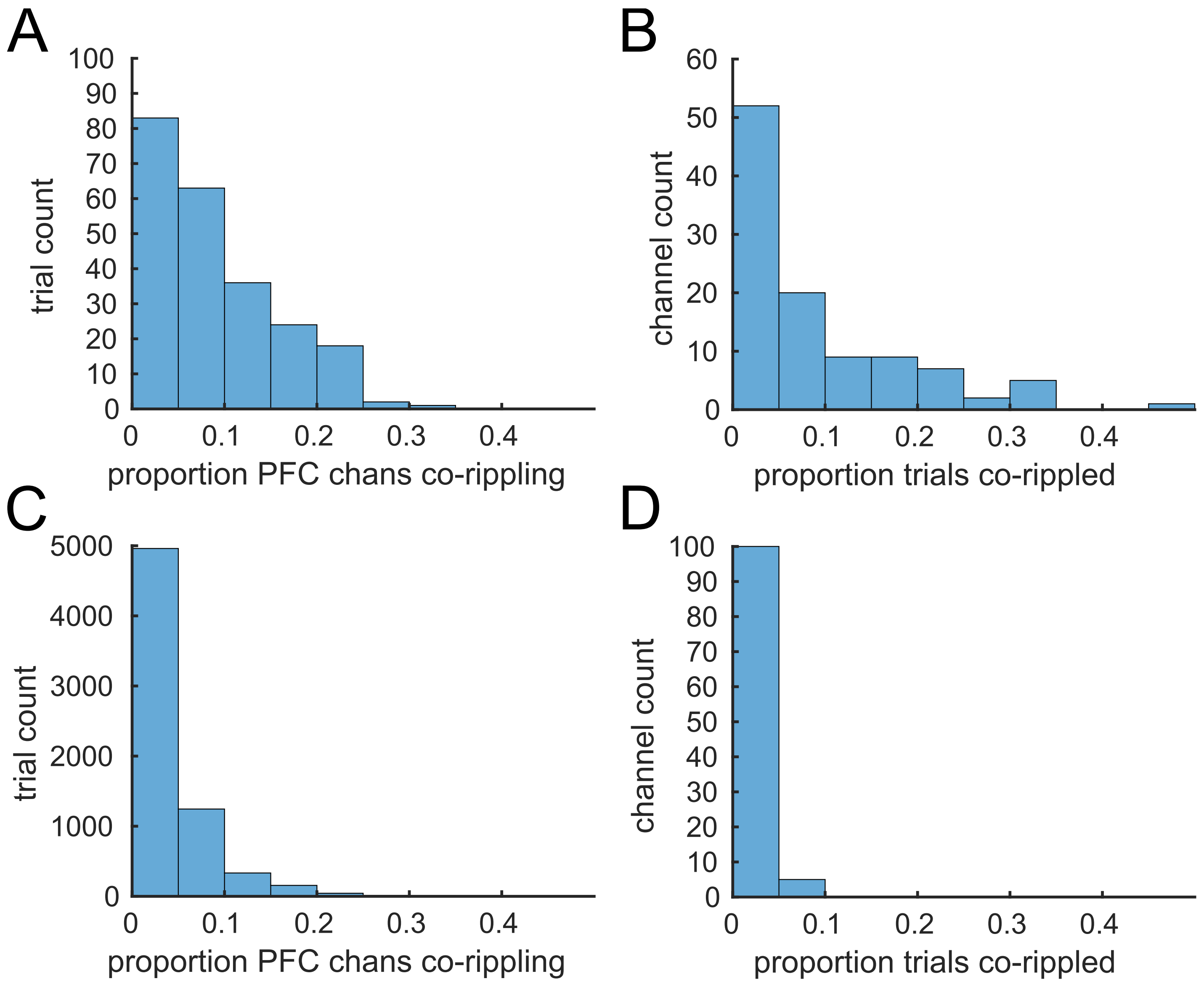


**Fig. S17. Distributions of Target-evoked co-ripple occurrence across both trials and channels displayed unimodal distributions.** Distributions are generated from the sample of channels in Prefrontal Cortex (PFC, including SFG, MFG, and IFG, see Table S2 for further detail) in subjects with at least 10 such channels each (P1,3 4,7,8; 105 total channels), and include only Co-R involving channels outside of PFC. Proportions reflect the number of PFC sites co-rippling with outside sites during this period. **A.** The proportion of channels with any Target-evoked Co-R showed a unimodal distribution over trials. **B.** Same as A for 100 ms pre-stimulus Baseline, provided for comparison. **C.** The proportion of Target trials (N = 227) in which Co-R were evoked showed a unimodal distribution across channels. **D.** Same as C, but for 100 ms pre-stimulus Baseline (N trials = 6766), provided for comparison.

### Tables

**Table S1. Patient characteristics.**

| Code | Age | Onset | Sex | Handed-ness | Wada language | Most contacts | #LH sites | #RH sites | Connection plots | Site plots | ISI |
|---|---|---|---|---|---|---|---|---|---|---|---|
| P1 | 27 | 15 | M | R | L | L | 94 | 7 | yes | yes | F |
| P2 | 40 | 3.5 | F | R | L | L | 86 | 0 | yes | yes | F |
| P3 | 51 | 37 | F | R | L | L | 97 | 0 | yes | yes | F |
| P4 | 45 | 4 | F | R | - | L | 100 | 0 | yes | yes | F |
| P5 | 23 | 14 | M | L | L | L | 98 | 0 | yes | yes | F |
| P6 | 24 | 4 | F | R | - | L | 106 | 0 | yes | yes | F |
| P7 | 27 | 4.5 | F | R | L | L | 172 | 62 | yes | yes | V |
| P8 | 39 | 5 | F | R | L | L | 104 | 0 | yes | yes | F,V |
| P9 | 48 | 3 | F | R | L | B | 64 | 79 | yes | yes | V |
| P10 | 16 | 11 | F | R | L | R | 28 | 158 | yes | yes | V |
| P11 | 50 | 8 | F | - | L | R | 0 | 142 | no | yes | V |
| P12 | 51 | 38 | F | R | L | R | 0 | 118 | no | yes | V |
| P13 | 18 | 7 | M | R | L | R | 0 | 74 | no | yes | F |

ISI: F= fixed at 600 ms; V= variable between 1000 and 1400 ms

**Table S2. Regions of interest**

| Region | Region short | Parcels |
|---|---|---|
| Angular gyrus | Ang | G_pariet_inf-Angular, S_interm_prim-Jensen |
| Dorsal and Lateral Occipital Cortex | Oc | G_cuneus, G_occipital_middle, G_occipital_sup, Pole_occipital, S_oc_middle_and_Lunatus, S_oc_sup_and_transversal, S_occipital_ant |
| Inferior Frontal Gyrus | IF | G_front_inf-Opercular, G_front_inf-Triangul, Lat_Fis-ant-Horizont, Lat_Fis-ant-Vertical, S_front_inf |
| Inferior Temporal Gyrus | IT | G_temporal_inf, S_collat_transv_ant, S_oc-temp_lat, S_temporal_inf |
| Lateral Fusiform Gyrus | lFu | G_oc-temp_lat-fusifor (fraction containing HCP FFC and portions of TF) |
| Lingual Gyrus | Ling | G_oc-temp_med-Lingual (excluding HCP V1 and V2), S_oc-temp_med_and_Lingual |
| Medial Fusiform Gyrus | mFu | G_oc-temp_lat-fusifor (fraction containing HCP VVC and V8, and portions of PHA3 and VMV3) |
| Medial Occipital Cortex | MO | S_calcarine, S_parieto_occipital, G_oc-temp_med-Lingual (including HCP V1 and V2) |
| Middle Frontal Gyrus | MF | G_front_middle, S_front_middle |
| Middle Temporal Gyrus | MT | G_temporal_middle, S_temporal_sup |
| Orbitofrontal Cortex | OF | G_and_S_frontomargin, G_and_S_transv_frontopol, G_front_inf-Orbital, G_orbital, G_rectus, G_subcallosal, S_orbital_lateral, S_orbital_med-olfact, S_orbital-H_Shaped, S_suborbital |
| Parahippocampal Gyrus | PH | G_oc-temp_med-Parahip |
| Postcentral Gyrus | Pos | G_postcentral, S_postcentral |
| Precentral Gyrus | Pre | G_precentral S,_precentral-inf-part, S_precentral-sup-part |
| Subcentral Gyrus | SC | G_and_S_subcentral |
| Superior Frontal Gyrus | SF | G_and S front_sup |
| Superior Temporal Gyrus | ST | G_temp_sup-G_T_transv, G_temp_sup-Lateral, G_temp_sup-Plan_polar, G_temp_sup-Plan_tempo, S_temporal_transverse |
| Supramarginal Gyrus | SM | G_pariet_inf-Supramar, Lat_Fis-post |
| Temporal Pole | TP | Pole_temporal |
| Ventral Occipital Cortex | VO | G_and_S_occipital_inf, S_collat_transv_post |

**Table S3. Ripple characteristics by region.**

| ROI | amp. mean (µV) | amp. SD (µV) | freq. mean (Hz) | freq. SD (Hz) | dur. mean (ms) | dur. SD (ms) | rate mean ($min^{-1}$) | rate SD ($min^{-1}$) |
|---|---|---|---|---|---|---|---|---|
| MO | 16.5 | 7.6 | 90.5 | 5.9 | 122 | 95 | 13.7 | 5.9 |
| VO | 10.4 | 2.8 | 90.5 | 5.7 | 96 | 52 | 19.7 | 5.1 |
| Oc | 15.0 | 8.9 | 90.5 | 5.9 | 114 | 85 | 16.3 | 8.8 |
| Lg | 14.4 | 5.9 | 91.0 | 6.1 | 113 | 74 | 17.9 | 7.8 |
| mFu | 12.2 | 4.3 | 90.3 | 6.0 | 107 | 64 | 22.3 | 8.0 |
| lFu | 16.4 | 9.1 | 91.8 | 6.4 | 125 | 84 | 22.6 | 16.1 |
| IT | 9.2 | 3.7 | 89.1 | 5.8 | 110 | 72 | 14.4 | 4.9 |
| MT | 9.3 | 4.1 | 89.3 | 6.0 | 114 | 77 | 15.2 | 5.4 |
| ST | 9.4 | 4.2 | 90.1 | 5.8 | 126 | 93 | 15.7 | 6.0 |
| TP | 8.1 | 2.9 | 90.4 | 6.2 | 105 | 87 | 13.2 | 4.1 |
| PH | 17.6 | 11.4 | 87.9 | 6.2 | 111 | 69 | 19.7 | 6.7 |
| SC | 7.7 | 3.5 | 89.9 | 5.9 | 105 | 77 | 13.7 | 4.7 |
| Ang | 11.2 | 4.5 | 90.0 | 5.6 | 136 | 109 | 17.2 | 7.1 |
| SM | 9.2 | 4.5 | 90.0 | 5.8 | 116 | 78 | 13.9 | 5.9 |
| OF | 10.4 | 3.8 | 89.7 | 6.1 | 128 | 103 | 17.7 | 4.9 |
| IF | 10.5 | 4.0 | 89.8 | 5.9 | 117 | 83 | 15.6 | 6.4 |
| MF | 11.0 | 3.9 | 89.8 | 5.9 | 114 | 74 | 16.0 | 4.2 |
| SF | 10.1 | 4.2 | 90.5 | 6.1 | 104 | 70 | 17.0 | 6.8 |
| SP | 8.8 | 3.0 | 91.1 | 5.7 | 102 | 57 | 12.2 | 4.8 |
| Pre | 10.5 | 3.5 | 90.1 | 5.9 | 106 | 69 | 13.3 | 4.9 |
| Pos | 10.2 | 4.0 | 90.0 | 5.9 | 123 | 102 | 12.3 | 4.4 |

Frequency was measured by the spacing of zero crossings in 70-110 Hz bandpass. Amplitude was quantified as baseline to peak of the Hilbert analytic amplitude. See Table S2 for region labels.

**Table S4. Region-region co-ripple relative density in Novel>Baseline contrast, 200 – 400 ms post-stimulus.**
Corresponds to Fig,. 3A.

| | MO | VO | Oc | Lg | mFu | lFu | IT | MT | ST | TP | PH | SC | Ang | SM | OF | IF | MF | SF | SP | Pre | Pos |
|---|---|---|---|---|---|---|---|---|---|---|---|---|---|---|---|---|---|---|---|---|---|
| MO | | | | | | | | | | | | | | | | | | | | | |
| VO | | | | | | | | | | | | | | | | | | | | | |
| Oc | | | | | | 2.6 | | | | | | | | | | | | | | | |
| Lg | | | | | | | | | | | | | | | | | | | | | |
| mFu | | | | | | 7.4 | 3.5 | 3.2 | | 3.3 | 7.3 | 3.3 | 3.8 | 2.4 | 3.4 | 3.4 | 3.1 | 2.9 | | 2.7 | |
| lFu | | | 2.6 | | 7.4 | | 2.5 | 2.2 | | | | | | | | 2.2 | 2.7 | | | 2.8 | 2.7 |
| IT | | | | | 3.5 | 2.5 | | | | | | | | | | | | | | | |
| MT | | | | | 3.2 | 2.2 | | | | | | | | | | | | | | | |
| ST | | | | | | | | | | | | | | | | | | | | | |
| TP | | | | | 3.3 | | | | | | | | | | | | | | | | |
| PH | | | | | 7.3 | | | | | | | | | | | | | | | | |
| SC | | | | | 3.3 | | | | | | | | | | | | | | | | |
| Ang | | | | | 3.8 | | | | | | | | | | | | | | | | |
| SM | | | | | 2.4 | | | | | | | | | | | | | | | | |
| OF | | | | | 3.4 | | | | | | | | | | | | | | | | |
| IF | | | | | 3.4 | 2.2 | | | | | | | | | | | | | | | |
| MF | | | | | 3.1 | 2.7 | | | | | | | | | | | | | | | |
| SF | | | | | 2.9 | | | | | | | | | | | | | | | | |
| SP | | | | | | | | | | | | | | | | | | | | | |
| Pre | | | | | 2.7 | 2.8 | | | | | | | | | | | | | | | |
| Pos | | | | | | 2.7 | | | | | | | | | | | | | | | |

**Table S5. Region-region co-ripple relative density in Target>Baseline contrast, 400 – 600 ms post-stimulus.**
Corresponds to Fig. 3C.

| | MO | VO | Oc | Lg | mFu | lFu | IT | MT | ST | TP | PH | SC | Ang | SM | OF | IF | MF | SF | SP | Pre | Pos |
|---|---|---|---|---|---|---|---|---|---|---|---|---|---|---|---|---|---|---|---|---|---|
| MO | | | | | | | | | | | | | | | | 8.4 | | 6.5 | | | |
| VO | | | | | | | | | | | | | | | | | | | | | |
| Oc | | | | | 4.4 | 5.6 | | | | | | | | | | | | | | | |
| Lg | | | | | | | | | | | | | | | | | | | | | |
| mFu | | | 4.4 | | | 8.6 | 8.4 | 4.7 | | | | | 5.3 | 4.5 | | | 4.4 | 9.4 | 20 | 7.9 | |
| lFu | | | 5.6 | | 8.6 | | | | | | | | | | | | 4.0 | | | 5.6 | |
| IT | | | | | 8.4 | | | | | | | | | | | | | | | | |
| MT | | | | | 4.7 | | | | | | | | | | | | | | | | |
| ST | | | | | | | | | | | | | | | | | | | | | |
| TP | | | | | | | | | | | | | | | | | | | | | |
| PH | | | | | | | | | | | | | | | | | | | | | |
| SC | | | | | | | | | | | | | | | | | | | | | |
| Ang | | | | | 5.3 | | | | | | | | | | | 3.9 | | | | | |
| SM | | | | | 4.5 | | | | | | | | | | | | | | | | |
| OF | | | | | | | | | | | | | | | | | | | | | |
| IF | 8.4 | | | | | | | | | | | | 3.9 | | | | 2.5 | 7.5 | 7.9 | | |
| MF | | | | | 4.4 | 4.0 | | | | | | | | | | 2.5 | | 3.7 | | | |
| SF | 6.5 | | | | 9.4 | | | | | | | | | | | 7.5 | 3.7 | | 7.3 | | |
| SP | | | | | 20 | | | | | | | | | | | 7.9 | | 7.3 | | | |
| Pre | | | | | 7.9 | 5.6 | | | | | | | | | | | | | | | |
| Pos | | | | | | | | | | | | | | | | | | | | | |

**Table S6. Region-region co-ripple relative density in Target>Novel contrast, 600 – 800 ms post-stimulus.** Corresponds to Fig. 3D.

| | MO | VO | Oc | Lg | mFu | lFu | IT | MT | ST | TP | PH | SC | Ang | SM | OF | IF | MF | SF | SP | Pre | Pos |
|---|---|---|---|---|---|---|---|---|---|---|---|---|---|---|---|---|---|---|---|---|---|
| MO | | | | | | | | | | | | | | | | | | | 5.0 | 3.8 | |
| VO | | | 8.9 | | | | 5.1 | | | | | | | | 7.9 | | 8.4 | 11 | | 12 | |
| Oc | | 8.9 | | | | 4.2 | 3.5 | | | | | | | 3.6 | 3.5 | 5.3 | 6.8 | 7.5 | 12 | 4.8 | 5.2 |
| Lg | | | | | | | 6.3 | 5.7 | | | | | | | | | | | | 6.9 | 5.8 |
| mFu | | | | | | 4.7 | 4.6 | 3.7 | | | 16 | | | 3.1 | | 4.1 | 6.2 | | | 4.4 | 5.6 |
| lFu | | | 4.2 | | 4.7 | | 3.9 | | | | | | | | | 3.7 | 5.8 | 13 | | 4.6 | 6.8 |
| IT | | 5.1 | 3.5 | 6.3 | 4.6 | 3.9 | | | | | | | | | | 3.4 | 4.0 | 3.5 | | 3.9 | 3.4 |
| MT | | | | 5.7 | 3.7 | | | | | | | | | | | | 2.9 | 3.9 | 15 | 3.3 | 3.0 |
| ST | | | | | | | | | | | | | | | | | 2.3 | 3.1 | 5.2 | | |
| TP | | | | | | | | | | | | | | | | | 3.2 | 6.0 | 20 | | |
| PH | | | | | 16 | | | | | | | | 13 | | | | 6.4 | | | | |
| SC | | | | | | | | | | | | | | | | | | | 6.1 | | |
| Ang | | | | | | | | | | | 13 | | | | | 6.3 | 9.2 | 9.9 | 13 | 3.7 | |
| SM | | | 3.6 | | 3.1 | | | | | | | | | | | 4.2 | 6.7 | 7.3 | 5.8 | 7.6 | 4.4 |
| OF | | 7.9 | 3.5 | | | | | | | | | | | | | 2.4 | 3.5 | 5.2 | 16 | 4.5 | 2.6 |
| IF | | | 5.3 | | 4.1 | 3.7 | 3.4 | | | | | | 6.3 | 4.2 | 2.4 | | 4.1 | 4.4 | 8.9 | 3.9 | 6.4 |
| MF | | 8.4 | 6.8 | | 6.2 | 5.8 | 4.0 | 2.9 | 2.3 | 3.2 | 6.4 | | 9.2 | 6.7 | 3.5 | 4.1 | | 8.7 | 24 | 6.4 | 7.7 |
| SF | | 11 | 7.5 | | | 13 | 3.5 | 3.9 | 3.1 | 6.0 | | | 9.9 | 7.3 | 5.2 | 4.4 | 8.7 | | 16 | 11 | 13 |
| SP | 5.0 | | 12 | | | | | 15 | 5.2 | 20 | | 6.1 | 13 | 5.8 | 16 | 8.9 | 24 | 16 | | 8.6 | 6.8 |
| Pre | 3.8 | 12 | 4.8 | 6.9 | 4.4 | 4.6 | 3.9 | 3.3 | | | | | 3.7 | 7.6 | 4.5 | 3.9 | 6.4 | 11 | 8.6 | | 7.8 |
| Pos | | | 5.2 | 5.8 | 5.6 | 6.8 | 3.4 | 3.0 | | | | | | 4.4 | 2.6 | 6.4 | 7.7 | 13 | 6.8 | 7.8 | |

**Table S7. Region-region co-ripple relative density in True Positive>False Positive contrast, -200 – 0 ms peri-response.** Corresponds to Fig. 4C.

| | MO | VO | Oc | Lg | mFu | lFu | IT | MT | ST | TP | PH | SC | Ang | SM | OF | IF | MF | SF | SP | Pre | Pos |
|---|---|---|---|---|---|---|---|---|---|---|---|---|---|---|---|---|---|---|---|---|---|
| MO | | | | | | | | | | | | | | | | | | | | | |
| VO | | | | | | | | | | | | | | | | | | | | | |
| Oc | | | | | | | | | | | | | | | | | | | | | |
| Lg | | | | | | | | | | | | | | | | | | | | | |
| mFu | | | | | | | | | | | | | | | | | | | | | |
| lFu | | | | | | | | | | | | | | | | | | | | | |
| IT | | | | | | | | | | | | | | | | | | | | | |
| MT | | | | | | | | | | | | | | | | | | | | | |
| ST | | | | | | | | | | | | | | | | | | | | | |
| TP | | | | | | | | | | | | | | | | | | | | | |
| PH | | | | | | | | | | | | | | | | | | | | | |
| SC | | | | | | | | | | | | | | | | | | | | | |
| Ang | | | | | | | | | | | | | | | | 5.4 | | | | | |
| SM | | | | | | | | | | | | | | | | | | | | | |
| OF | | | | | | | | | | | | | | | | | | 5.7 | | 2.3 | |
| IF | | | | | | | | | | | | | 5.4 | | | | 2.4 | 7.3 | | | |
| MF | | | | | | | | | | | | | | | | 2.4 | | 3.8 | | | |
| SF | | | | | | | | | | | | | | | 5.7 | 7.3 | 3.8 | | | 3.1 | 5.8 |
| SP | | | | | | | | | | | | | | | | | | | | | |
| Pre | | | | | | | | | | | | | | | 2.3 | | | 3.1 | | | 1.9 |
| Pos | | | | | | | | | | | | | | | | | | 5.8 | | 1.9 | |

**Table S8. Region-region co-ripple counts.**

| | MO | VO | Oc | Lg | mFu | lFu | IT | MT | ST | TP | PH | SC | Ang | SM | OF | IF | MF | SF | SP | Pre | Pos |
|---|---|---|---|---|---|---|---|---|---|---|---|---|---|---|---|---|---|---|---|---|---|
| MO | | 91 | 1089 | 100 | 237 | 148 | 551 | 649 | 1153 | 208 | 126 | 601 | 716 | 953 | 1214 | 431 | 1387 | 1583 | 757 | 814 | 740 |
| VO | 91 | | 122 | 53 | 41 | 92 | 199 | 205 | 303 | 44 | 6 | 80 | 18 | 134 | 115 | 183 | 140 | 77 | 88 | 212 | 147 |
| Oc | 1089 | 122 | | 717 | 1229 | 1170 | 2943 | 4268 | 3290 | 700 | 156 | 1631 | 1840 | 3152 | 3769 | 2445 | 3081 | 2019 | 925 | 2741 | 2661 |
| Lg | 100 | 53 | 717 | | 327 | 355 | 762 | 990 | 843 | 288 | 72 | 242 | 244 | 476 | 747 | 904 | 749 | 223 | 77 | 635 | 844 |
| mFu | 237 | 41 | 1229 | 327 | | 785 | 1471 | 1968 | 1500 | 439 | 180 | 619 | 615 | 1174 | 1589 | 1139 | 1348 | 702 | 201 | 1370 | 1116 |
| lFu | 148 | 92 | 1170 | 355 | 785 | | 1362 | 1934 | 1434 | 461 | 51 | 375 | 408 | 696 | 1380 | 1777 | 1298 | 593 | 153 | 1383 | 1335 |
| IT | 551 | 199 | 2943 | 762 | 1471 | 1362 | | 8859 | 5753 | 1766 | 698 | 1841 | 1992 | 4256 | 5069 | 3186 | 3600 | 1922 | 442 | 3592 | 3424 |
| MT | 649 | 205 | 4268 | 990 | 1968 | 1934 | 8859 | | 9546 | 2696 | 1067 | 2763 | 3483 | 6568 | 9497 | 5270 | 5683 | 2980 | 602 | 5392 | 4518 |
| ST | 1153 | 303 | 3290 | 843 | 1500 | 1434 | 5753 | 9546 | | 2499 | 974 | 3236 | 2582 | 4996 | 8323 | 4702 | 5691 | 4833 | 1186 | 4920 | 4235 |
| TP | 208 | 44 | 700 | 288 | 439 | 461 | 1766 | 2696 | 2499 | | 460 | 584 | 393 | 888 | 2394 | 1375 | 1458 | 896 | 183 | 1180 | 1189 |
| PH | 126 | 6 | 156 | 72 | 180 | 51 | 698 | 1067 | 974 | 460 | | 267 | 158 | 342 | 882 | 359 | 783 | 447 | 108 | 371 | 190 |
| SC | 601 | 80 | 1631 | 242 | 619 | 375 | 1841 | 2763 | 3236 | 584 | 267 | | 1356 | 3384 | 2689 | 1278 | 2255 | 2151 | 659 | 2356 | 1872 |
| Ang | 716 | 18 | 1840 | 244 | 615 | 408 | 1992 | 3483 | 2582 | 393 | 158 | 1356 | | 3448 | 3706 | 1221 | 2636 | 2610 | 880 | 1810 | 1613 |
| SM | 953 | 134 | 3152 | 476 | 1174 | 696 | 4256 | 6568 | 4996 | 888 | 342 | 3384 | 3448 | | 5061 | 1683 | 3123 | 2901 | 957 | 3781 | 3445 |
| OF | 1214 | 115 | 3769 | 747 | 1589 | 1380 | 5069 | 9497 | 8323 | 2394 | 882 | 2689 | 3706 | 5061 | | 5630 | 8109 | 6726 | 1378 | 4923 | 3746 |
| IF | 431 | 183 | 2445 | 904 | 1139 | 1777 | 3186 | 5270 | 4702 | 1375 | 359 | 1278 | 1221 | 1683 | 5630 | | 5813 | 3025 | 570 | 4212 | 3408 |
| MF | 1387 | 140 | 3081 | 749 | 1348 | 1298 | 3600 | 5683 | 5691 | 1458 | 783 | 2255 | 2636 | 3123 | 8109 | 5813 | | 7116 | 1812 | 5457 | 3697 |
| SF | 1583 | 77 | 2019 | 223 | 702 | 593 | 1922 | 2980 | 4833 | 896 | 447 | 2151 | 2610 | 2901 | 6726 | 3025 | 7116 | | 1947 | 3640 | 2401 |
| SP | 757 | 88 | 925 | 77 | 201 | 153 | 442 | 602 | 1186 | 183 | 108 | 659 | 880 | 957 | 1378 | 570 | 1812 | 1947 | | 958 | 672 |
| Pre | 814 | 212 | 2741 | 635 | 1370 | 1383 | 3592 | 5392 | 4920 | 1180 | 371 | 2356 | 1810 | 3781 | 4923 | 4212 | 5457 | 3640 | 958 | | 4620 |
| Pos | 740 | 147 | 2661 | 844 | 1116 | 1335 | 3424 | 4518 | 4235 | 1189 | 190 | 1872 | 1613 | 3445 | 3746 | 3408 | 3697 | 2401 | 672 | 4620 | |

**Table S9. Region-region co-ripple PLV and zero-latency bias.** Corresponding co-ripple phase distributions shown in Fig. S15.

| | MO | VO | Oc | Lg | mFu | lFu | IT | MT | ST | TP | PH | SC | Ang | SM | OF | IF | MF | SF | SP | Pre | Pos |
|---|---|---|---|---|---|---|---|---|---|---|---|---|---|---|---|---|---|---|---|---|---|
| MO | | 0.11 | 0.26 | 0.11 | 0.19 | 0.21 | 0.13 | 0.12 | 0.21 | 0.30 | 0.29 | 0.22 | 0.24 | 0.24 | 0.19 | 0.15 | 0.19 | 0.17 | 0.23 | 0.17 | 0.20 |
| VO | 0.11 | | 0.21 | 0.04 | 0.17 | 0.06 | 0.09 | 0.11 | 0.02 | 0.14 | 0.19 | 0.07 | 0.24 | 0.08 | 0.20 | 0.06 | 0.03 | 0.12 | 0.28 | 0.14 | 0.09 |
| Oc | 0.26 | 0.21 | | 0.19 | 0.13 | 0.18 | 0.13 | 0.13 | 0.24 | 0.38 | 0.24 | 0.26 | 0.33 | 0.19 | 0.18 | 0.25 | 0.33 | 0.31 | 0.47 | 0.25 | 0.26 |
| Lg | 0.11 | 0.04 | 0.19 | | 0.17 | 0.16 | 0.15 | 0.13 | 0.15 | 0.35 | 0.16 | 0.15 | 0.07 | 0.08 | 0.17 | 0.18 | 0.18 | 0.32 | 0.19 | 0.21 | 0.20 |
| mFu | 0.19 | 0.17 | 0.13 | 0.17 | | 0.16 | 0.22 | 0.14 | 0.11 | 0.34 | 0.21 | 0.17 | 0.15 | 0.11 | 0.17 | 0.26 | 0.18 | 0.32 | 0.27 | 0.15 | 0.16 |
| lFu | 0.21 | 0.06 | 0.18 | 0.16 | 0.16 | | 0.26 | 0.16 | 0.16 | 0.30 | 0.10 | 0.18 | 0.27 | 0.21 | 0.18 | 0.13 | 0.19 | 0.32 | 0.39 | 0.13 | 0.19 |
| IT | 0.13 | 0.09 | 0.13 | 0.15 | 0.22 | 0.26 | | 0.26 | 0.34 | 0.55 | 0.24 | 0.28 | 0.16 | 0.14 | 0.27 | 0.29 | 0.28 | 0.27 | 0.36 | 0.26 | 0.23 |
| MT | 0.12 | 0.11 | 0.13 | 0.13 | 0.14 | 0.16 | 0.26 | | 0.42 | 0.47 | 0.29 | 0.26 | 0.20 | 0.14 | 0.26 | 0.27 | 0.28 | 0.31 | 0.43 | 0.22 | 0.19 |
| ST | 0.21 | 0.02 | 0.24 | 0.15 | 0.11 | 0.16 | 0.34 | 0.42 | | 0.56 | 0.36 | 0.39 | 0.31 | 0.26 | 0.36 | 0.24 | 0.31 | 0.40 | 0.41 | 0.26 | 0.35 |
| TP | 0.30 | 0.14 | 0.38 | 0.35 | 0.34 | 0.30 | 0.55 | 0.47 | 0.56 | | 0.53 | 0.52 | 0.43 | 0.46 | 0.61 | 0.43 | 0.43 | 0.47 | 0.46 | 0.43 | 0.49 |
| PH | 0.29 | 0.19 | 0.24 | 0.16 | 0.21 | 0.10 | 0.24 | 0.29 | 0.36 | 0.53 | | 0.34 | 0.41 | 0.45 | 0.40 | 0.15 | 0.19 | 0.26 | 0.53 | 0.36 | 0.27 |
| SC | 0.22 | 0.07 | 0.26 | 0.15 | 0.17 | 0.18 | 0.28 | 0.26 | 0.39 | 0.52 | 0.34 | | 0.35 | 0.42 | 0.24 | 0.38 | 0.40 | 0.49 | 0.55 | 0.42 | 0.37 |
| Ang | 0.24 | 0.24 | 0.33 | 0.07 | 0.15 | 0.27 | 0.16 | 0.20 | 0.31 | 0.43 | 0.41 | 0.35 | | 0.20 | 0.18 | 0.25 | 0.38 | 0.37 | 0.61 | 0.32 | 0.28 |
| SM | 0.24 | 0.08 | 0.19 | 0.08 | 0.11 | 0.21 | 0.14 | 0.14 | 0.26 | 0.46 | 0.45 | 0.42 | 0.20 | | 0.17 | 0.29 | 0.37 | 0.45 | 0.56 | 0.22 | 0.26 |
| OF | 0.19 | 0.20 | 0.18 | 0.17 | 0.17 | 0.18 | 0.27 | 0.26 | 0.36 | 0.61 | 0.40 | 0.24 | 0.18 | 0.17 | | 0.25 | 0.31 | 0.29 | 0.31 | 0.23 | 0.33 |
| IF | 0.15 | 0.06 | 0.25 | 0.18 | 0.26 | 0.13 | 0.29 | 0.27 | 0.24 | 0.43 | 0.15 | 0.38 | 0.25 | 0.29 | 0.25 | | 0.27 | 0.22 | 0.27 | 0.28 | 0.33 |
| MF | 0.19 | 0.03 | 0.33 | 0.18 | 0.18 | 0.19 | 0.28 | 0.28 | 0.31 | 0.43 | 0.19 | 0.40 | 0.38 | 0.37 | 0.31 | 0.27 | | 0.30 | 0.38 | 0.34 | 0.31 |
| SF | 0.17 | 0.12 | 0.31 | 0.32 | 0.32 | 0.32 | 0.27 | 0.31 | 0.40 | 0.47 | 0.26 | 0.49 | 0.37 | 0.45 | 0.29 | 0.22 | 0.30 | | 0.42 | 0.29 | 0.43 |
| SP | 0.23 | 0.28 | 0.47 | 0.19 | 0.27 | 0.39 | 0.36 | 0.43 | 0.41 | 0.46 | 0.53 | 0.55 | 0.61 | 0.56 | 0.31 | 0.27 | 0.38 | 0.42 | | 0.44 | 0.48 |
| Pre | 0.17 | 0.14 | 0.25 | 0.21 | 0.15 | 0.13 | 0.26 | 0.22 | 0.26 | 0.43 | 0.36 | 0.42 | 0.32 | 0.22 | 0.23 | 0.28 | 0.34 | 0.29 | 0.44 | | 0.27 |
| Pos | 0.20 | 0.09 | 0.26 | 0.20 | 0.16 | 0.19 | 0.23 | 0.19 | 0.35 | 0.49 | 0.27 | 0.37 | 0.28 | 0.26 | 0.33 | 0.33 | 0.31 | 0.43 | 0.48 | 0.27 | |

**=Sig. PLV & Zero-Latency**

**=Non-sig. PLV**

**=Sig. PLV & Non-zero latency**
